# Ancient DNA reveals that natural selection has upregulated the immune system over the last 10,000 years

**DOI:** 10.64898/2026.04.14.718409

**Authors:** Javier Maravall-López, Buu Truong, Gaspard Kerner, Yujie Zhao, Kangcheng Hou, Annabel Perry, Ali Akbari, David Reich, Alkes L. Price

## Abstract

The specific mechanisms through which human biology and disease susceptibility evolved with major shifts in West Eurasian environments and societies over the last 10,000 years(*1*)—particularly rising infectious burden(*2*)—remain poorly characterized, despite ancient DNA studies(*3-6*) identifying hundreds of candidate loci under positive selection(*6*). Here, we identify specific immune diseases/traits, genes/variants, pathways, and tissues/cell types impacted by natural selection by systematically integrating variant-level selection statistics with genome-wide association study (GWAS), quantitative trait locus (QTL), and molecular bulk/single-cell and gene pathway data. Genome-wide, positively-selected alleles are associated with reduced susceptibility to infectious diseases like tuberculosis (TB), influenza, and intestinal infections; consistent with selection-signal enrichments in immune cells within barrier tissues such as the respiratory tract and gut mucosa. In contrast, positively-selected alleles increase risk of intestinal inflammatory disease and autoimmune hypothyroidism, supportive of a tradeoff between infection and immune-mediated pathology, and consistent with adaptive alleles being QTLs for genes upregulating inflammation and other host-defense pathways. We reveal many novel adaptive loci with convergent signals from selection, infectious disease GWAS and immune-gene QTLs (including at *FUT6* for intestinal infections; at *ASAP1* for TB; and at *LYZ*, an antimicrobial enzyme), fine-mapping selection onto likely causal variants. Surprisingly, adaptive alleles had a protective effect on allergic conditions like asthma and dermatitis, challenging a common view that these conditions arose through evolutionary mismatch of present-day hygienic contexts relative to past, pathogen-rich environments(*7*).

## Introduction

The last 10,000 years of West Eurasian history have witnessed profound changes in the lifestyle, social organization and mobility of human populations(*1*), contributing to rising infectious disease burden(*2*), which was highly lethal within this time transect(*8*). We can thus expect pathogens to be among the strongest selective pressures shaping the genome of ancient West Eurasians. It is widely speculated that adaptations to pathogens may have resulted in a trade-off with immune-mediated disease genetic susceptibility(*4, 9-11*) (through antagonistic pleiotropy, the phenomenon of a genetic variant affecting different traits in opposite directions). At the same time, allergic inflammatory conditions like asthma are hypothesized to arise through a mismatch between a genetic background adapted to past, pathogen-rich environments and modern, hygienic settings(*7*). However, much remains unknown about the particular genetic variations, genes, pathways and tissues/cell types that have been relevant to recent human immune genetic adaptation.

Ancient DNA is emerging as a promising tool to study recent human evolution directly through its unprecedented ability to track allele frequency changes through time(*3-6, 12*). In particular, Akbari et al.(*6*) discovered 479 independent variants with >99% probability of being under positive selection in West Eurasia over the last ten millennia, and found these variants to be enriched for immune-related trait heritability(*6*). However, this study did not investigate immune-related functional effects of adaptation, and did not attempt to identify the specific causal variants involved, limiting biological interpretation. In parallel, the two last decades of human genetics research have provided a wealth of datasets and methods to link genetic variants to biology(*13-15*). Integrating these diverse data modalities with ancient DNA-based selection scans has the potential to yield meaningful biological discoveries.

Here, we characterize the immune landscape of genome-wide signals of positive selection in ancient West Eurasians(*6*) across several levels of biological organization, identifying specific immune diseases/traits, genes/variants, pathways, and tissues/cell types impacted by natural selection by leveraging genome-wide association (GWAS), quantitative trait loci (QTL), curated pathway collections, and molecular bulk/single-cell datasets. We perform both local analyses at loci with evidence for selection, as well as polygenic analyses that take advantage of genome-wide patterns not limited to significant loci(*16*). Our results offer novel insights into the biological mechanisms underlying immune-related adaptations in West Eurasia.

## Results

### Overview of methods

We adapted 4 types of methods from the statistical genetics literature (developed for GWAS data) to selection summary statistics (**Figure 1a**). These methods: (1) jointly analyze the genetic architectures of selection statistics and disease GWAS(*17, 18*) (**Figure 1b**); (2) use molecular data to identify biologically causal genes and variants at loci implicated by selection statistics(*18-22*) (**Figure 1c**); (3) harness pathway-level enrichment and trans-protein QTL analyses of causal genes implicated by selection statistics(*23*) to identify biological processes up- or downregulated by selection (**Figure 1d**); and (4) identify critical tissues and cell types implicated by selection statistics(*22, 24-27*) (**Figure 1e**). A complete description of these methods is provided in the **Methods** section and **Table S3**.

**Figure 1.**
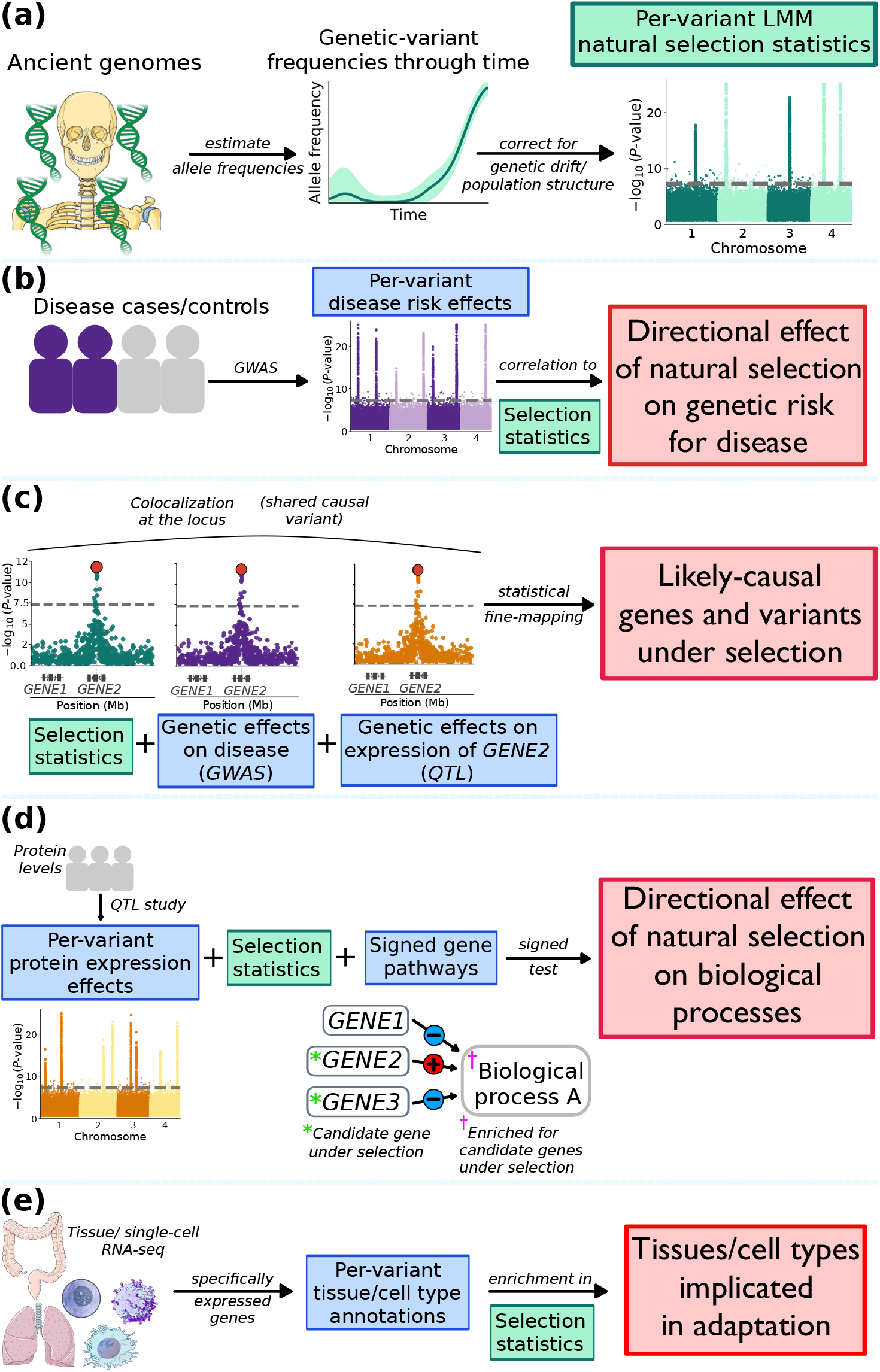
Overview of methods to link natural selection to disease risk, specific genes/variants, biological pathways, and tissues/cell types. **(a)** Per-variant LMM selection statistics are computed from ancient genomes, modifying the method of Akbari et al.(*6*). Illustrations are reused from Servier Medical Art (https://smart.servier.com), licensed under CC BY 4.0 (https://creativecommons.org/licenses/by/4.0/); and the NIAID NIH BioArt Source (https://bioart.niaid.nih.gov/124). **(b)** <u>Directional effects of natural selection on genetic disease risk</u> are estimated from LMM selection statistics and disease GWAS. **(c)** <u>Causal genes and variants targeted by natural selection</u> are identified from convergence of LMM selection statistics, disease GWAS, and expression QTL. **(d)** <u>Directional effects of natural selection on biological processes</u> are identified from LMM selection statistics (encoding positively-selected alleles), protein QTL (encoding protein-increasing alleles), and signed gene pathway annotations (encoding proteins up- or downregulating a given process). **(e)** <u>Tissues and fine-grained cell types in implicated adaptation</u> are identified from LMM selection statistics and bulk/single-cell RNA-seq. Illustrations are reused from Servier Medical Art (https://smart.servier.com), licensed under CC BY 4.0 (https://creativecommons.org/licenses/by/4.0/); and the NIAID NIH BioArt Source (https://bioart.niaid.nih.gov/82,612,376). In **(b)-(e)**, green boxes indicate natural selection inputs, blue boxes indicate auxiliary data inputs, and red boxes indicate outcomes of our analyses. See **Methods** for details of each method.

We jointly analyzed genome-wide selection statistics with GWAS association statistics for 2,989 diseases/traits (**Materials and Methods**) (including infection-related traits, immune-mediated diseases and traits related to immune-response mechanisms) (average *N*=277K); cis-eQTL association statistics from 50 GTEx tissues(*28*) (average *N*=439); cis-pQTL and trans-pQTL association statistics for 2,795 proteins in UK Biobank(*23*) (*N*=35K); and bulk/single-cell molecular datasets(*22, 28-33*) and curated gene pathways(*34-38*) (**Methods** and **Data Availability**). Aside from GWAS association statistics for a smaller number of traits (about a sixth), none of these auxiliary data types were analyzed by Akbari et al(*6*).

We determined that the selection statistics computed by Akbari et al. (using a generalized linear mixed model (GLMM)(*6*) violated some of the statistical assumptions necessary to produce unbiased results when analyzed with the GWAS methods that we applied. Consequently, we modified the Akbari et al.(*6*) approach to instead use a linear mixed model (LMM), and verified that LMM selection statistics constitute appropriate input for GWAS methods (see **Methods** and **Supplementary Text A**). Enrichment of LMM (positive) selection statistics in background selection closely matched that of GWAS (**Figure S18**,) consistent with our previous findings that background selection is not driving positive selection detection(*6*).

### Adaptation reduced infection risk and increased immune-mediated disease risk

We sought to assess whether positively-selected alleles show a genome-wide pattern of impacting risk for infectious and immune-mediated disease. We thus applied cross-trait LD score regression(*17*) (LDSC) to estimate genetic correlations between LMM selection statistics and GWAS association statistics (**Methods**), analyzing many new diseases/traits related to infection and immunity (vs. Akbari et al.(*6*) analyses using GLMM selection statistics). This method uses a stringent statistical-significance criterion that only identifies relationships that hold throughout the genome, not just at a subset of loci. Results are in **Figure 2a** and **Data S1**.

**Figure 2.**
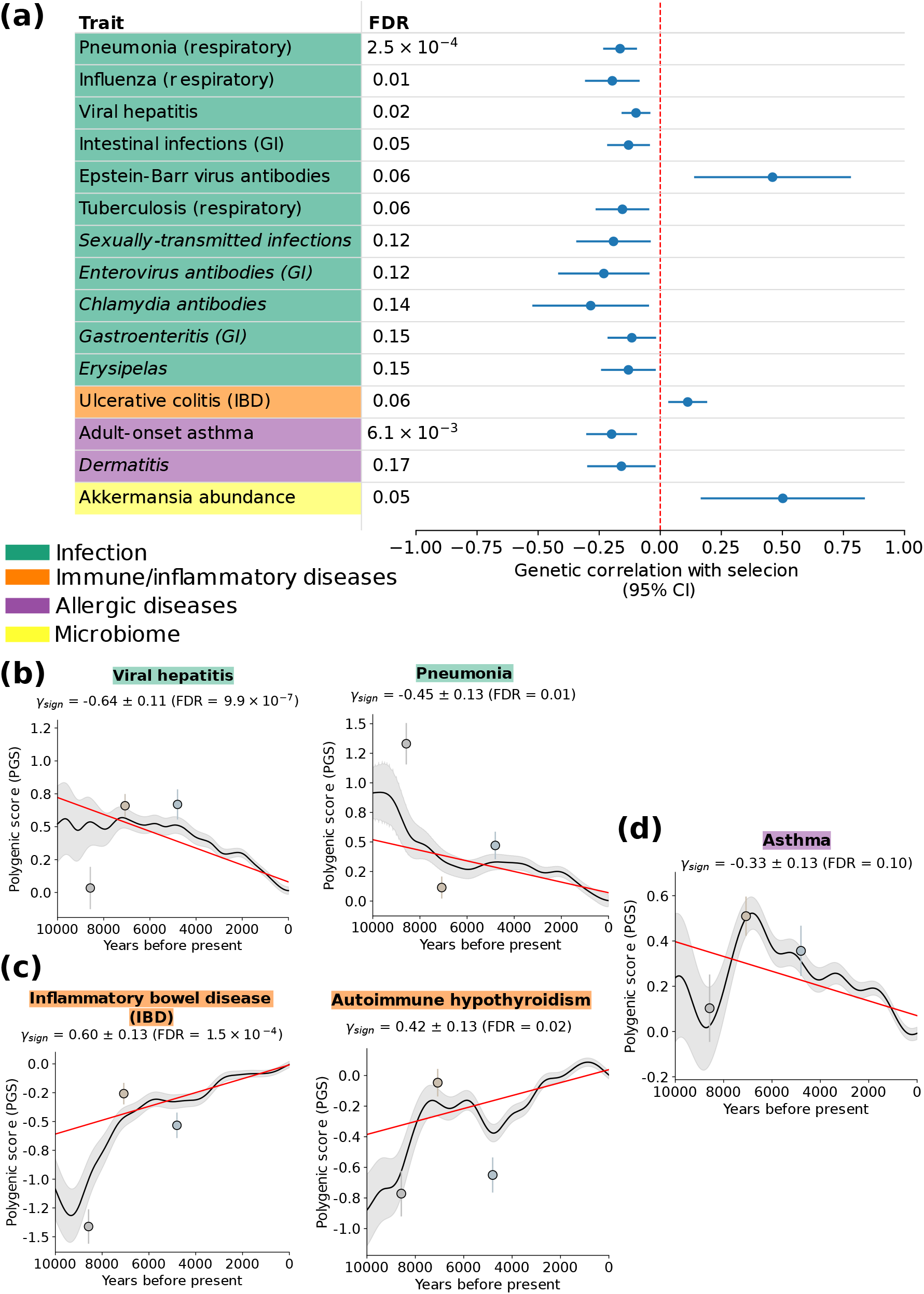
Natural selection decreased risk of infections and increased risk of immune-mediated diseases. **(a)** <u>Genetic correlations between natural selection and selected GWAS traits</u>. We report cross-trait LDSC genetic-correlation point estimates with 95% confidence intervals (CI) (1.96 × SE, computing standard errors (SE) via a genomic-block jackknife), for selected traits from immune-related categories. This analysis depends on statistical power, which is largely a function of sample size. Suggestive results (10% < FDR < 20%) are denoted in italics. GI: gastrointestinal. *Chlamydia trachomatis* causes a common, sexually-transmitted bacterial infection; erysipelas is a bacterial skin infection(*41*). **(b)** <u>Significant decreases over time in genetically-predicted infection risk</u>. Decreases were also detected for tuberculosis; and *Streptococcus pneumoniae*, a respiratory pathogen that is a major cause of pneumonia in humans(*41*) (**Figure S19a**). **(c)** <u>Significant increases over time in genetically-predicted immune-mediated disease risk</u>. An increase was also detected for a biomarker of autoimmune thyroid disease(*41*) (**Figure S19b**). **(d)** <u>Significant decrease over time in genetically-predicted asthma risk</u>. An increase was also detected for additional allergy-related phenotypes (**Figure S19c**). In **(b)**-**(d)**, the solid black line shows, for selected GWAS traits, the mean polygenic score (PGS) (an individual’s effect size-weighted dosage of risk-increasing alleles that is used as a genetic predictor of a trait, trained in modern European individuals) of western Eurasian individuals over the past 10,000 years, with the 95% confidence band in grey. The red line indicates a polygenic test that assesses whether this average genetic trait predictor has shifted in magnitude beyond the neutral expectation (via a linear mixed model regression adjusted for population structure, with slope γ_sign_, see **Methods** for details). Circles with error bars denote the mean PGS with 95% confidence intervals for representative groups of the three ancestral populations to modern Europeans(*64*), in order of temporal proximity to the present: Western hunter–gatherers (light gray; n = 131), early European farmers (light beige; n = 452), and steppe pastoralists (light blue; n = 293). In **(a)-(d)**, nominal statistical significance for a difference from 0 was assessed via a two-tailed *Z* test; FDR refers to Benjamini-Hochberg(*65*) (BH) *q*-values. See **Data S1** for complete numerical results.

Significant (FDR < 10%) results for infection-related traits included negative genetic correlations for susceptibility to pneumonia; influenza; intestinal infections; viral hepatitis; and tuberculosis (TB, widely considered to be among the deadliest infectious diseases in human history(*39*)), consistent with the strongest known homozygote genetic risk factor for TB(*40*) being reportedly under negative selection in the last three millennia(*6, 12*). A covariance-aware meta-analysis of 12 approximately genetically independent infection-disease traits (**Methods**) produced a significantly negative estimate: −0.0445 (SE 0.0162, one-tailed *p <* 0.003 from a *Z*-test), highlighting a general pattern of positively-selected alleles being associated with decreased infection susceptibility. The single significant exception was antibody levels against Epstein-Barr (EB) virus. Because infection by this virus is almost universal in human cohorts, it is possible that inter-individual differences in antibody levels are driven by overall strength of the immune response rather than susceptibility; in that case, the positive correlation would indicate a broad enhancer effect of adaptation on immune response. Additional analyses also supported genetic overlap between selection and aggregate infectious diseases (**Figure S22a**).

For immune-mediated diseases, we identified a positive genetic correlation with ulcerative colitis (UC), a common form of inflammatory intestinal bowel disease (IBD)(*41*), which could be consistent with antagonistic pleiotropy with infection-related adaptation(*4, 9-11*), and falls in line with previous reports of IBD genetic risk having increased over time in West Eurasia(*4*). For allergic diseases, we identified a negative genetic correlation with adult-onset asthma risk, an allergic inflammatory trait characterized by hyperreactivity of the airway immune response(*41*). This challenges the commonly-held view that these types of conditions arise through alleles that were protective against past, pathogen-rich environments becoming maladaptive in modern, hygienic contexts(*7*). For microbiome traits, an unexpected result was a positive genetic correlation with abundance of *Akkermansia*, a bacterium in the human microbiome(*42*) (plausibly related to adaptation enhancing gut barrier mucosal immunity, see below).

As an alternative way to study directional effects of natural selection on genetic disease risk, we applied a polygenic test developed by Akbari et al.(*6*). This test assesses whether alleles that predict disease risk in present-day populations are collectively increasing or decreasing in frequency across time in ancient individuals, beyond what can be explained by genetic drift/population structure; this test can be better powered than cross-trait LDSC under sparse genetic architectures (**Methods**). Results (**Data S1**) confirmed a trend of significant decreasing risk for infectious diseases (**Figure 2b** and **Figure S19a**), of increasing risk for immune-mediated diseases (including for autoimmune hypothyroidism, not detected by cross-trait LDSC, **Figure 2c** and **Figure S19b**), and of decreasing risk for allergic conditions (**Figure 2d** and **Figure S19c**).

We applied coloc(*18*) to assess colocalization (shared causal variants) at loci with evidence of both selection and GWAS association (**Methods**) (this analysis depends on statistical power, which is largely a function of sample size). We identified 413 unique loci colocalizing between LMM selection statistics and GWAS (**Figures S20**,**21**); these included 72 for autoimmune diseases and 41 for infection-related traits. Complete results are reported in **Data S3**, and notable specific loci are discussed in the next section. We also performed statistical fine-mapping(*18, 43*) to identify likely causal variants under selection (**Table S4**) (without regard to GWAS) (**Methods**).

We performed a secondary analysis to identify GWAS traits that may have been causally under selection, by applying the Latent Causal Variable method(*19*) (LCV; **Methods; Figure S22b** and **Data S1**). Roughly, LCV assesses whether variants that impact an upstream trait impact a downstream trait, extending Mendelian Randomization (MR) methods, which may be confounded by horizontal pleiotropy(*44*). Traits inferred to be causal for selection included lighter skin pigmentation (FDR *<* 10^−20^) and higher vitamin D levels (FDR *<* 10^−20^) (widely reported to be under selection(*3, 5, 6, 45, 46*) and hence viewed as positive controls); and a suggestive result for lower antibody levels (likely proxying lower susceptibility) for *Helicobacter pylori*, a gastric pathogen detected in the ancient DNA record >5,000 years ago(*47*). Conversely, selection was suggestively causal for higher abundance of *Akkermansia* (also see **Figure 2a**). Because this commensal is a mucin specialist that lives in the gut mucus layer such that increased mucin availability tends to increase its levels(*42*), its abundance is likely proxying stronger gut-barrier immunity (see below).

We conclude that selection favored alleles associated with genetic resistance to infections in present-day populations, as well as risk alleles for immune-mediated diseases and alleles protective against allergies.

### Fine-mapping pinpoints immune genes and variants under selection

Identifying the effector genes underlying selection signals is not always straightforward (analogous to disease GWAS). In order to identify specific genes impacted by adaptation, we used two complementary approaches. First, we applied coloc(*18*) to assess colocalization between loci implicated as significant by both selection statistics and molecular QTL association statistics, including cis-eQTL for 18,597 protein-coding genes across 50 GTEx(*28*) tissues (average *N*=439); cis-eQTL for 5,459 protein-coding genes across 5 immune cell types from the eQTL Catalogue(*48*) (average *N*=130) (**Data S3**); and cis-pQTL and trans-pQTL for 2,795 proteins in plasma in the UK Biobank(*23*) (*N*=35K) (**Methods**). Second, we applied CALDERA(*20*), a method for fine-mapping causal genes using fine-mapped coding variants at a candidate locus, distance to genes, and genome-wide patterns of enrichment for particular gene features(*21, 22*). Each of these approaches was previously developed for GWAS association statistics, but applied here to LMM selection statistics.

We identified colocalizations between selection statistics and molecular QTL at 464 unique loci (including at 65% of genome-wide significant loci for selection, **Figure S23a**), implicating 1,719 protein-coding genes (885 in *cis*), and 71 genes with CALDERA posterior inclusion probability (PIP_CALDERA_)>0.5 (for which power is likely limited by variant-level selection fine-mapping resolution) (**Data S3**). There was substantial overlap between the respective sets of implicated genes (**Data S4**).

We highlight 6 examples of selection-GWAS colocalization (**Data S3**) in **Figure 3**, focusing on infection-related and immune-response GWAS traits; these examples implicate genes (and, in most cases, a specific variant) related to infection and host-microbe interactions impacted by adaptation, with most including eQTL or pQTL colocalization. We are not currently aware of previous evidence of selection impacting these 6 loci.

**Figure 3.**
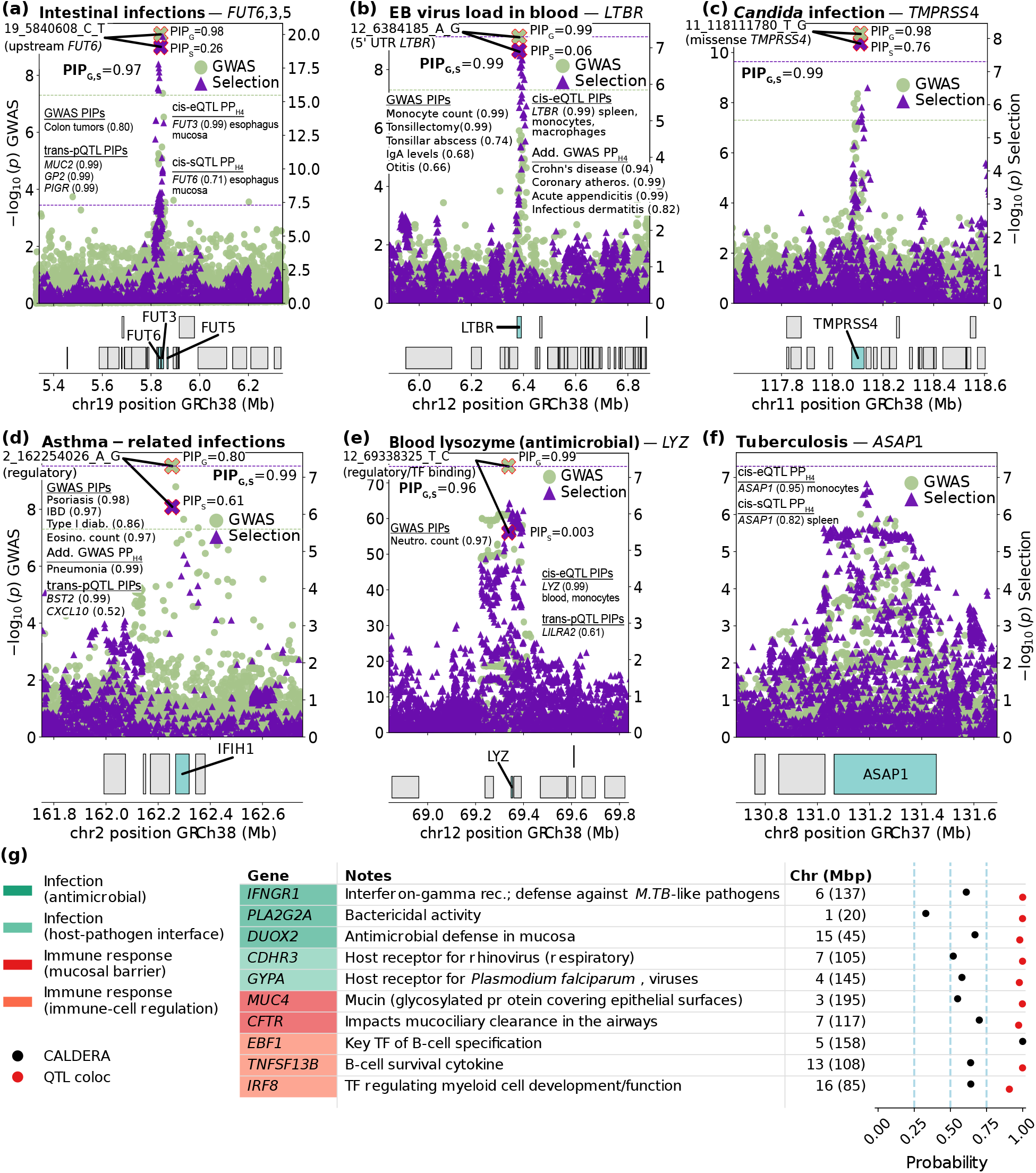
Examples of colocalizations of selection with infection-related GWAS and immune-related QTLs. **(a-f)** <u>Examples of colocalizations between selection and GWAS related to infection</u>. In most cases (**(a-e)**), we were able to nominate specific likely causal variants through fine-mapping, and link them to a biologically plausible gene through QTL colocalization or coding variation. PP_H4_: posterior probability of a shared causal variant between selection and the listed GWAS trait or QTL gene, as assessed by coloc(*18*) (**Methods**). e,s,pQTL: expression, splicing, protein QTL (in plasma), respectively. PIP_S_=posterior probability of a variant being causal for selection. PIP_G_=posterior probability of a variant being causal for the focal GWAS trait. PIP_S,G_=posterior probability of a variant being the shared causal variant for selection and the focal GWAS trait conditional on positive colocalization (computed by coloc, **Methods**), multiplied by PP_H4_. PIP=posterior probability of a variant being causal for the listed GWAS trait or QTL gene. We note that 19_5840608_C_T (from **(a)**) is also a fine-mapped *trans*-pQTL (PIP>0.99 in each case) for *FAM3D, REG1A, REG1B* and *CEACAM5*; and that 12_6384185_A_G (from **(b)**) is also a fine-mapped *trans*-pQTL (PIP>0.97 in each case) for *CCL21* and *TNFRSF9*. Information on the function of these and other genes mentioned in **(a)**-**(f)** is given in **Table S5**. In **(d)**, we caution that rs1990760, an *IFIH1* missense variant absent from the selection scan, belonged to fine-mapped credible sets for FinnGen(*66*) infectious endpoints at this locus. **(g)** <u>Examples of adaptive genes prioritized by QTL colocalization and/or gene-level fine-mapping with CALDERA</u>. We report 10 genes prioritized for adaptation, together with probabilities of being causal for selection according to gene-level fine-mapping with CALDERA(*20*) (**Methods**); or probabilities of adaptation colocalizing with QTLs for that gene as assessed by coloc(*18*) (**Methods**). We made an effort to group genes into functional categories, but we caution that some genes could plausibly be assigned to more than one category. Notes were adapted from protein-function descriptions in ref.(*51*). Some genes are discussed in more detail in **Supplementary Text B**. Rec.: receptor; *M. TB: Mycobacterium tuberculosis*, bacterium causing tuberculosis(*39*). *Plasmodium falciparum* is a protozoon causing malaria(*67*); TF: transcription factor, a DNA-binding protein that regulates gene transcription(*51*). We note that *PLA2G2A* and *TNFSF13B* are upregulated in infectious disease(*68*). In the case of *PLA2G2A*, we were able to nominate a specific 5’ UTR variant at this gene as the likely causal selected variant (**Table S6**).

First, we identified a selection-intestinal infection colocalization (PP_H4_>0.99, where PP_H4_ is the posterior probability of a shared causal variant) upstream of the *FUT6* gene (**Figure 2a**), strongly expressed in apical epithelial cells(*49*) and part of a family of genes involved in post-translational modifications in mucosal surfaces known to impact interactions with pathogens(*50*); selection at this locus also colocalized with the expression of *FUT3* in esophagus mucosa, an adaptation-critical tissue (see below). We implicate a single SNP with posterior inclusion probability (PIP) for selection (PIP_Sel_)=0.26, PIP for GWAS (PIP_GWAS_)=0.98, and joint PIP_Sel,GWAS_=0.97, which is also a fine-mapped *trans*-pQTL (PIP>0.99 in each case) for 7 genes involved in gut immunity (**Table S5**); this variant was also fine-mapped (PIP=0.80) for colon-tumor GWAS.

Second, we identified a selection-EB virus load colocalization (PP _H4_>0.99) at the *LTBR* gene (**Figure 2b**), modulating immune response and lymphoid tissue development(*51*). We implicate a single 5’ UTR SNP with PIP_Sel_=0.06, PIP_GWAS_=0.99, and joint PIP_Sel,GWAS_=0.99, which is also a fine-mapped *cis-*eQTL (PIP>0.99) for *LTBR* and a fine-mapped *trans*-pQTL (PIP>0.97 in each case) for 2 host-defense genes (**Table S5**). This locus also colocalized with Crohn’s disease (a form of IBD) risk.

Third, we identified a selection-fungal infection colocalization (PP_H4_>0.99) inside the *TMPRSS4* gene (**Figure 2c**), expressed in epithelial mucosa and previously shown to impact viral infection susceptibility through its role as a host receptor(*52*). We implicate a single missense SNP with PIP_Sel_=0.76, PIP_GWAS_=0.98, and joint PIP_Sel,GWAS_>0.99 (**Methods**).

Fourth, we identified a selection-asthma infections colocalization (PP _H4_>0.99) on chromosome 2 (162.2 Mbp) (**Figure 2d**). We implicate a single regulatory SNP with PIP_Sel_=0.61, PIP_GWAS_=0.80, and joint PIP_Sel,GWAS_>0.99, which is also a fine-mapped *trans-*pQTL (PIP>0.99) for *BST2*, an interferon-induced antiviral protein(*51*). This variant was also fine-mapped (PIP>0.85 in each case) for 3 autoimmune diseases, including IBD. We were not able to implicate a particular gene in *cis* through QTL colocalization; the proximal interferon-inducing *IFIH1*, involved in innate antiviral immunity(*51*), is a plausible candidate.

Fifth, we identified a selection-lysozyme (an antimicrobial enzyme(*41*)) blood levels colocalization (PP _H4_=0.96) at the *LYZ* gene (**Figure 2e**), coding for this enzyme(*51*). In this case, we implicate a single regulatory SNP with PIP_Sel_=0.003, PIP_GWAS_=0.99, and joint PIP_Sel,GWAS_=0.96, which is also a fine-mapped *cis-* eQTL (PIP>0.99) for *LYZ* and a fine-mapped *trans*-pQTL (PIP=0.61) for *LILRA2*, a gene involved in response to microbial infection(*51*) (**Table S5**). The top PIP_Sel_ at the locus was only 0.04, within a 123-member 95% credible set (**Data S2**), highlighting how incorporation of additional data layers beyond selection statistics alone may advance resolution of adaptive loci.

Sixth, we identified a selection-TB colocalization (posterior probability of colocalization (PP_H4_ = 0.80) at the *ASAP1* gene (**Figure 2f**), a gene implicated in susceptibility to TB in European- and Han Chinese-ancestry populations as well as zebrafish models(*53-55*); we also identified a selection-*ASAP1* splicing colocalization at this locus in 3 GTEx tissues including the immune-related spleen, and a selection-*ASAP1* expression colocalization in two immune cell types, T cells and monocytes (**Data S3**). We were not able to nominate a specific causal variant at this locus (the top shared PIP was only 0.098).

We also highlight one example involving a variant widely known to be under selection at *TLR1(56)* (**Figure S24a**), and a plausible example of two independent signals at the same selected locus (**Figure S24b**).

Many QTL colocalizations did not coincide with infectious GWAS colocalizations but nonetheless implicated genes related to infectious disease and immune response. We highlight 10 such examples in **Figure 3g** (see **Supplementary Text B**; additional examples are reported in **Table S5**); given our implication of TB, *IFNGR1*, the gene encoding the IFN‐γ (interferon gamma) receptor component 1 (PIP_CALDERA_=0.61) was noteworthy. The interferon gamma signaling axis sits at the heart of host immune response to mycobacterial pathogens(*57, 58*) (the genus containing *M. tuberculosis*) and other intramacrophagic pathogens: IFN‐γ binding to its receptor on macrophages triggers these cells to clear mycobacteria(*41*). We also identified many other examples for which selection-QTL colocalization for immune genes without infectious-disease GWAS colocalization implicated specific variants (**Table S6**), and other loci for which selection-GWAS (for non-immune traits) and selection-QTL colocalization implicated specific variants at genes not primarily related to immunity (**Table S7**).

As a secondary analysis, we quantified the proportion of selection signals explained by variants near immune genes to be roughly one third of that explained by variants near all genes (**Figure S23b**). This suggests that even though immune-related biology is enriched for selection signals (the per-variant selection-signal variance explained was roughly 50% greater on average near immune genes than near all genes, see **Figure S23c**), adaptation also occurred along other biological axes (potentially, metabolism or reproduction, since such traits produced highly significant genetic correlations with selection, **Data S1**; see also **Table S7**).

These results identify many genes related to infections and immune response that were impacted by adaptation, and demonstrate how integration of multiple data modalities nominates likely causal variants at adaptive loci.

### Natural selection upregulated immune gene pathways

We sought to assess whether particular gene pathways were overrepresented among genes impacted by adaptation, using 10,480 pathways from gene ontology(*34, 35*); 4,023 canonical pathways(*35*); 5,748 pathways from human phenotype ontology(*35, 36*); 50 pathways from the Hallmark collection(*35*); and 46 pathways from Inborn Errors of Immunity, a curated library of gene sets derived from the IUIS classification of monogenic immune disorders(*37*). We assessed enrichment of each pathway for 936 genes with GTEx QTL colocalization (PP_H4_>0.8) or 516 genes with CALDERA PIP>0.1 (156 genes were included in both gene sets; excess overlap 133.6x, two-tailed hypergeometric *p* < 10^−20^) (we excluded plasma pQTL and immune QTL colocalization due to differences in available gene sets; **Data S4**); we assessed the statistical significance of enrichment using a hypergeometric test (**Methods**). Importantly, we further assessed whether adaptation increased or decreased the activity of immune processes.

We identified 1,469 pathways enriched for GTEx QTL colocalization genes and 868 pathways enriched for CALDERA PIP>0.1 genes (FDR<10%) (**Data S4**). Both sets of enriched pathways exhibited excess overlap with a set of 153 immune-related pathways curated by ImmPort(*38*) (**Figure 4a, Figure S25a** and **Data S4**), with odds ratios of 6.41 (one-tailed permutation *p* < 10^−6^) and 1.98 (one-tailed permutation p < 9.3 × 10^−3^), respectively, in a logistic regression adjusted for pathway size (**Methods**). Pathway enrichments for GTEx QTL colocalization genes and CALDERA PIP>0.1 genes were concordant (Pearson correlation of *r =* 0.70 of log odds ratios across all pathways tested, two-tailed permutation *p* < 10^−6^), consistent with significant overlap between these sets of genes (**Data S4**).

**Figure 4.**
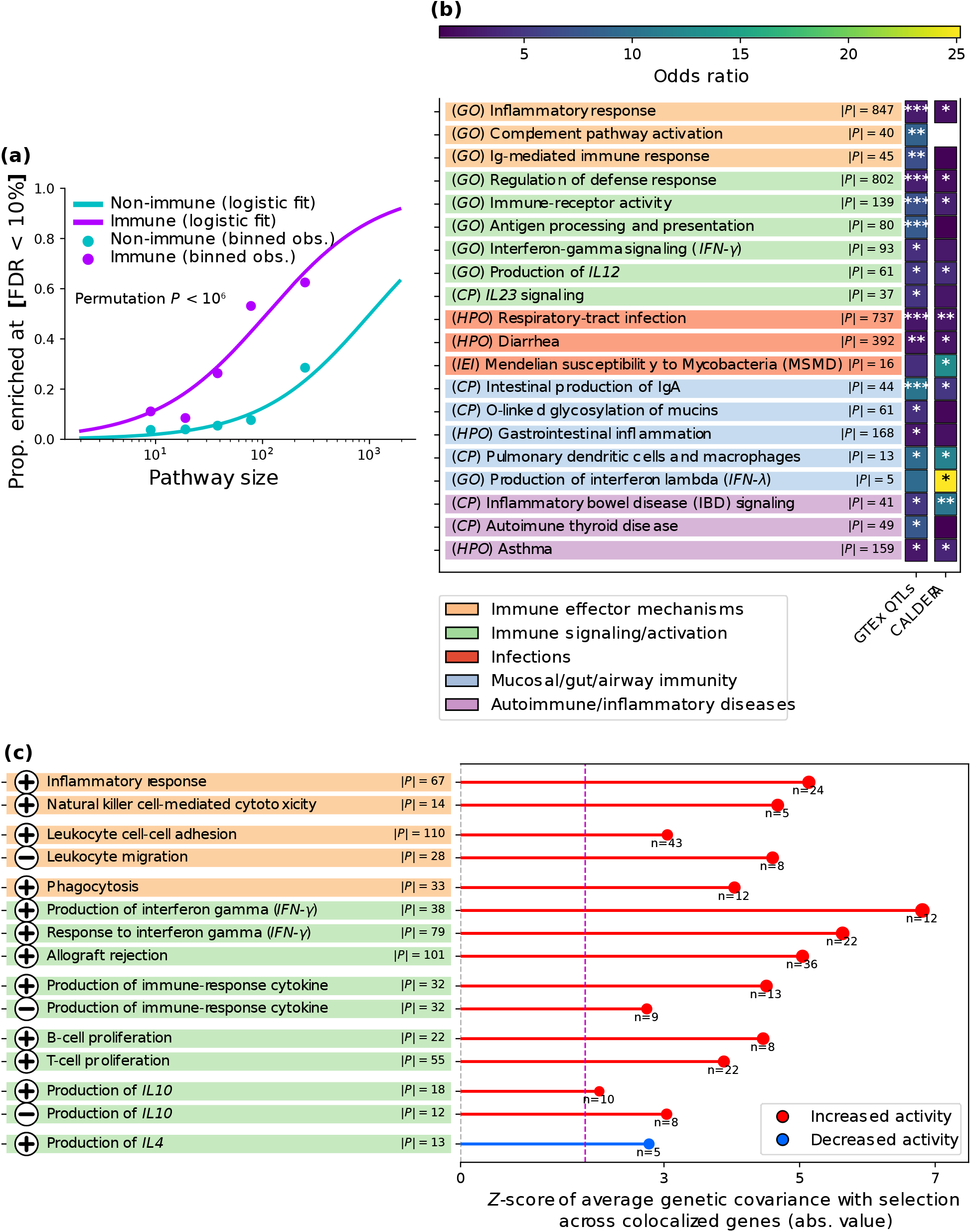
Prioritized genes for selection are enriched for immune-related pathways. **(a)** <u>Immune-related pathways are significantly more likely to be enriched for genes with a selection-GTEx QTL colocalization</u>. Dots represent proportion of pathways with enrichment FDR < 10% for 5 size quintiles (teal for non-immune, purple for immune). Curves represent a logistic regression fit adjusting for pathway size. *P*-values are computed via permutations to take into account redundancy in gene content across pathways. Analogous results for CALDERA>0.1 genes are reported in **Figure S25a**. Numerical results are reported in **Data S4. (b)** <u>Examples of pathways that are significantly enriched for genes prioritized for selection</u>. We report enrichment results for genes with a GTEx QTL colocalization, and genes with CALDERA PIP > 0.1. The color of each square represents the odds ratio (*: FDR<0.10, **: FDR<10^−3^, ***: FDR<10^−5^). For each pathway, |*P*| indicates the number of genes in the pathway intersecting the background gene universe (the set of genes that could have been included in the prioritized set) in each case. IL12 is a key upstream cytokine produced by activated macrophages/monocytes and conventional dendritic cells that drives *IFN-γ* production (supporting *IFN-γ*–centered host defense central to defense against mycobacteria(*41*)); *IL23* shares the p40 subunit with *IL12*, and can also impact anti-mycobacterial defense(*69*). **(c)** <u>Examples of pathways for which selection increased or decreased activity</u>. The color code of pathway names is the same as in **(b)**. For each pathway, we report the *Z*-score for the average genetic covariance with selection across selection-pQTL colocalized genes (genetic correlations are not available in general due to some genes having very noisy heritability estimates), for variants with |*Z*| > 3 for association with selection and plasma levels of the corresponding protein (genome-wide estimates were required to be qualitatively consistent, see **Methods**). To account for covariance in estimation errors or gene co-regulation, averages were jackknifed with 200 blocks of approximately equally-sized adjacent variants(*70*) (**Methods**). For each pathway, |*P*| indicates the number of genes in the pathway intersecting the background gene universe of plasma pQTLs; *n* indicates the number of genes used for the |*Z*| > 3 computation (some genes were dropped if enforcing that condition led to at least one of the 200 jackknife blocks used having no data). The dashed purple vertical line indicates a significantly (FDR < 10%) non-zero average genetic covariance. Z-scores are colored by the predicted effect of positively-selected alleles on pathway activity (i.e., increased for a positive average genetic covariance with a set of positive regulators or for a negative one with a set of negative regulators; conversely for decreased activity). Symbols indicate whether the inference was obtained using a set of positive (plus sign) or negative (minus sign) regulators of a given process (these gene sets have low overlap, **Data S4**). In some cases (e.g., production of immune-response cytokine), we show our inference using both positive and negative regulators of the same process; the 6 such pairs all produced directionally-concordant conclusions (permutation one-tailed *p* < 0.07), indicating the robustness of our predictions.

We highlight 15 examples of enriched immune-related pathways in **Figure 4b**, spanning 5 categories (including respiratory/gut infections, mucosal immunity and immune-mediated diseases, consistent with results implicating the corresponding GWAS trait categories (**Figure 2**) and gene categories (**Figure 3g**) in adaptation). Five of these examples are particularly notable. First, we implicated pathways related to IBD signaling (FDR < 1.7 × 10^−4^), autoimmune thyroid disease (FDR < 1.1 × 10^−3^) and asthma (FDR < 0.006); consistent with GWAS results (**Figure 2**). Second, we implicated an immune gene network for IgA (a mucosal-immunity antibody(*41*)) production in the intestines (FDR < 7.5 × 10^−6^). Third, we implicated the O-linked glycosylation of mucins pathway (FDR < 0.026), consistent with previous results implicating mucin genes (**Figures 2c,g**), and indicating a broader adaptive role of mucin biology(*59, 60*). Fourth, we implicated the Mendelian Susceptibility to Mycobacterial Disease (MSMD) pathway (FDR < 0.02); MSMD genes have a causal role in susceptibility to mycobacterial infection (including TB), established through loss-of-function carriers(*58*), and they were suggestively enriched for TB GWAS (**Data S4**). Fifth, we implicated the *IFN*‐*γ* signaling pathway (FDR < 3.3 × 10^−4^), central to host defense against mycobacteria and other intramacrophagic pathogens(*57, 58*).

Alternative approaches to linking selection to gene sets also highlighted immune-related pathways (**Figure S25b**,**c**).

We sought to further assess whether adaptation systematically upregulated or downregulated specific biological processes. We sub-selected 158 pathways that (i) were significantly enriched for GTEx QTL colocalization or CALDERA>0.1, and (ii) consisted of either genes that are all positive or all negative regulators of a given process, or genes that are all upregulated or all downregulated under a particular biological condition (from the Hallmark collection(*35*)). For these pathways, we inferred increased activation if adaptive alleles on average increase the expression of their positive regulators (or decrease the expression of their negative regulators; analogously for decreased activation; **Methods**) according to genome-wide (including *trans*) plasma pQTLs (*N*=50K).

A total of 74 of the 158 pathways had an average genetic covariance significantly different from 0 (FDR < 10%) (**Data S4**). Of these, 35 pathways were immune (33 with predicted increased activation, one-tailed permutation *p <* 7.0 × 10^−4^), and 39 were non-immune (22 with predicted increased activation, one-tailed permutation *p <* 0.41); the odds ratio for immune process enrichment for increased activation was 12.75 (one-tailed permutation *p <* 6.5 × 10^−4^). These results highlight a pervasive pattern of adaptation increasing immune pathway activity. This falls in line with genetically-predicted spleen volume having increased over time (FDR < 8.1 × 10^−11^, **Data S1**), since this organ plays a central role in immune function(*41*).

We highlight 15 of the 35 immune pathways (14 upregulated, 1 downregulated) in **Figure 4c**. Of these, 5 are particularly notable. First, adaptation increased inflammatory response, consistent with previous reports that pro-inflammatory eQTLs are enriched for positive-selection signals(*61*). This also falls in line with our polygenic test highlighting a significant increase over time in genetically-predicted levels of IL8 (**Figure S19c**), a major mediator of inflammation(*41*). Second, adaptation increased immune effector processes, including phagocytosis (the cellular uptake of microbes or debris by macrophages or other immune cells(*41*)) and natural-killer-cell-mediated cytotoxicity (the immune cell-mediated directed killing of infected or malignant cells(*41*)). Third, adaptation increased proliferation of B and T cells (types of leucocytes that help fight infections(*41*)), suggesting that selection enhanced the capacity for a rapid immune response. Fourth, adaptation increased *IFN*‐*γ* production, which could be related to selection pressure from intramacrophagic pathogens(*57, 58*); alternatively, it may just be part of this pro-inflammatory trend. Fifth, adaptation decreased production of *IL4*, an interleukin that promotes allergic immunity and is closely linked to asthma and atopic dermatitis(*41*); this result offers a (perhaps partial) molecular explanation of adaptive alleles being risk-decreasing for these diseases (**Figure 2a**).

We conclude that adaptation impacted many pathways related to immunity and infection, with a general trend towards increased activation.

### Immune cells in pathogen-interfacing tissues contributed to adaptation

We hypothesized that signals of selection might be driven by genes that impact particular tissues, and sought to identify these tissues using methods previously applied to GWAS. We first examined the gene features (such as expression in a given tissue or membership of a given pathway) for which PoPS(*22*) identified gene-level polygenic enrichment for signals of selection (**Methods**). We clustered gene features implicated by PoPS into orthogonal clusters (following ref.(*22*), in which PoPS clusters were informative about trait-specific biology), and manually annotated the top 25 clusters (of which 22 could be unambiguously annotated) (**Methods**). The top 10 (and 18 of 22 annotated) clusters were immune-related, and the 22 annotated clusters were all related to blood/immune, respiratory, digestive and barrier tissues (**Figure 5a, Data S5**). The implication of immune cells in the airway epithelium was of particular interest, and consistent with previous results implicating respiratory infections via (i) GWAS (**Figure 2**), (ii) genes (**Figure 3g**) and pathways (**Figure 4b**) and (iii) selection-GWAS colocalizations with antibody reactivity to respiratory viruses (**Data S3**).

**Figure 5.**
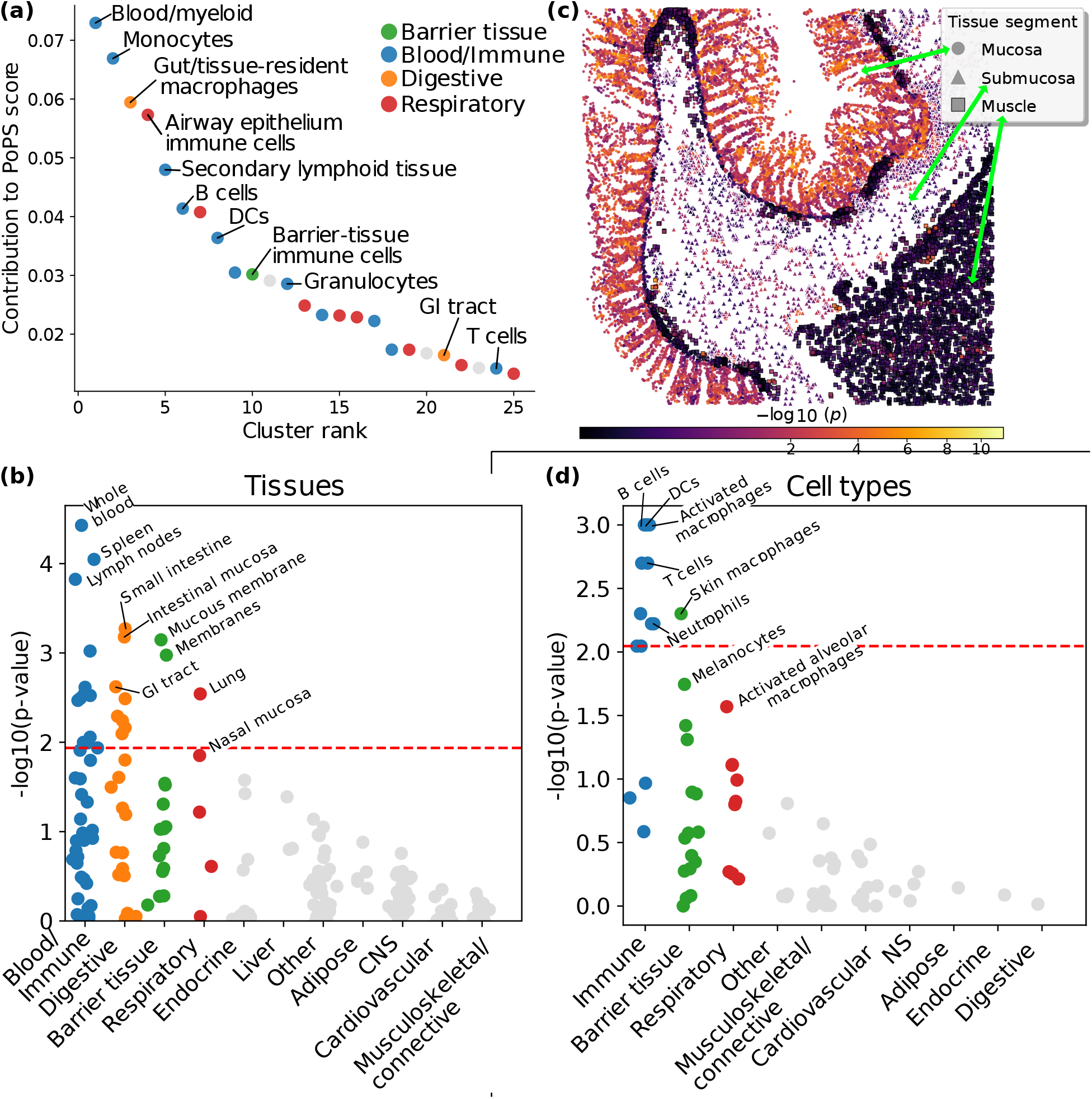
Enrichment of specific tissues and cell types for signals of selection. **(a)** <u>Top 25 PoPS clusters of orthogonal gene features selected for gene-level polygenic enrichment in signals of selection</u>. Clusters were manually annotated and we show selected labels (generally chosen to be non-redundant). Complete numerical results including selected features with names, cluster membership and justification of label choices are reported in **Data S5**. Gray clusters could not be unambiguously annotated. Remaining clusters are colored according to tissue category as indicated by the legend. DCs: dendritic cells, GI: gastrointestinal. **(b)** <u>Tissues with enrichment for signals of selection inferred by LDSC-SEG</u>. Dots represent tissues, partitioned into tissue categories (columns). Only categories with at least one significant (FDR < 10%, dashed red line) tissue are colored, with the same color code as in **(a)**. Selected tissues are labeled. Numerical results are reported in **Data S5. (c)** <u>Spatial patterns of individual-cell enrichment in gut tissue inferred by gsMap</u>. Dots represent spatially-arranged intestinal cells(*32*) from a selected sample, colored by *p*-value for enrichment for selection signals. The tissue segment that each cell belongs to is indicated. See **Figures S33**-**36** for results for other samples, and **Data S5** for complete numerical results. Formal category-level *p*-values are reported in **Figure S27a**,**b. (d)** <u>Cell types with enrichment for signals of selection inferred by scDRS</u>. Dots represent cell types, partitioned into cell-type categories (columns) chosen to reflect tissue-specific cell types. A mapping between cell-type label and tissue of origin is provided in **Figure S38**. Only categories with at least one significant (FDR < 10%, dashed red line) or suggestive (FDR < 15%) cell type are colored, with the same color code as in **(a)**. Selected cell types are labeled. Numerical results are reported in **Data S6**.

We next applied LDSC-SEG(*25*) to selection statistics using expression data from GTEx(*28*) (human) and the Franke lab(*30, 31*) (human, mouse, rat), as in ref.(*25*). LDSC-SEG applies S-LDSC(*24*) to evaluate whether variants proximal to genes specifically expressed in a particular tissue are enriched for disease heritability (or in our case, signals of selection), controlling for proximity to any gene (**Methods**). Consistent with PoPS results, blood/immune, digestive, barrier and respiratory tissues showed significant enrichment (**Figure 4b, Data S5**). Analyses of additional functional data sets further corroborated the enrichment of blood/immune and digestive tissues (**Figure S26**).

We next applied gsMap(*26*) to a single-cell spatial transcriptomics dataset across 8 distinct regions from the human small intestine and colon(*32*); gsMap applies S-LDSC(*24*) to evaluate spatial patterns of enrichment for disease heritability (or in our case, signals of selection) in individual cells (**Methods**). We pursued this analysis because PoPS specifically implicated immune cells in respiratory tissue, such that we sought to refine our implication of digestive tissues. We reached two main conclusions. First, aggregating across intestinal regions, gsMap detected enrichment specific to mucosal cells (**Figure 4c, Data S5, Figure S27a**). This is consistent with top LDSC-SEG tissues being mucosal (**Figure 5b**). Second, when stratifying mucosal cells by major communities — immune (51% of mucosal cells, **Figure S28**) and epithelial (49% of mucosal cells)— gsMap detected specific enrichment of immune cells (**Figure S27b**); results were qualitatively consistent for individual intestinal regions and when stratifying cells by finer spatial neighborhoods or finer cell type (**Supplementary Text C**). These findings linking adaptation to gut immune cells are consistent with several of our previous results (**Supplementary Text C.1**).

We next assessed whether fine-grained cell types across tissue categories are enriched for selection signals. We applied scDRS(*27*) to scRNA-seq data from scGTEx across 8 tissues(*33*); scDRS is a method that leverages scRNA-seq data to associate fine-grained cell types to disease GWAS (or in our case, adaptation) (**Methods**). Cell type-level scDRS results are reported in **Figure 4d** and **Data S6**. All cell types implicated by scDRS (FDR < 10%) were immune-related, including (i) cells from the mononuclear phagocyte system (e.g., activated macrophages and dendritic cells), including skin-resident macrophages, consistent with phagocytosis as a selection-implicated pathway (**Figure 4b**); (ii) adaptive-immunity lymphocytes, including B cells and T cells, consistent with selection increasing proliferation of these cell types according to our pathway analyses (**Figure 4c**); and (iii) granulocytes (e.g., neutrophils, cells that engulf microbes and release antimicrobial enzymes(*41*), consistent with our pathway analyses highlighting immune effector mechanisms (**Figures 4b**,**c**)). All of these cell types were also implicated by PoPS clusters (**Figure 4a**). Suggestive associations included melanocytes (FDR < 0.11), consistent with selection for pigmentation(*3, 5, 6, 45*); and alveolar (lung air sac) macrophages (FDR < 0.15), consistent with PoPS implicating immune cells in respiratory tissue (**Figure 4a**). The latter is consistent with our implication of TB above, since alveolar macrophages are the first immune cells *M. tuberculosis* meets in the lungs(*62*), though macrophage enrichment is also broadly consistent with infectious processes. Alternative approaches to link adaptation to specific cell types reached concordant conclusions (**Figure S39**).

Given that positively-selected alleles tend to reduce susceptibility to respiratory and intestinal infections (**Figure 2**), these results provide support for adaptation having enhanced front-line defense at the primary bodily interfaces of interaction with pathogens, such as the respiratory epithelium and the gut mucosa.

## Discussion

We have presented multiple lines of evidence revealing that natural selection over the last millennia has conferred resistance to infectious disease by widely upregulating the human immune system, implicating many previously-unknown specific variants and genes under positive selection. The coordinated trends across traits and pathways, as well as the consistent directionality of the observed signals, are not plausibly explained by confounders like background selection, and are instead a signature of directional adaptation.

Our findings are broadly consistent with the notion that adaptation to infection has increased genetic risk for immune-mediated diseases. However, we also obtained results that challenged common narratives about how risk for human diseases today might be enhanced due to adaptation to past environments in which humans no longer live. For example, we found a significant trend of positively-selected alleles increasing risk of inflammatory bowel disease as well as protecting against intestinal infections. It is tempting to posit a simple scenario of a tradeoff between these two patterns. However, the part of selection that tracks intestinal infection protection predicts *lower*, not higher, UC risk (**Supplementary Text D**), such that this positive correlation does not arise from selection acting through intestinal infection protection specifically. Rather, a component of adaptation enhancing gut mucosal barrier immunity, and a component promoting inflammatory tone, appear to have opposing effects on UC risk. Separately, we found favored alleles to have a significant protective effect against allergic inflammatory conditions like asthma, plausibly related to adaptation downregulating IL4 production. This is an unexpected finding because it runs counter to a common view that these conditions arose through evolutionary mismatch of present-day hygienic contexts relative to genetic adaptations to past environments where pathogens were abundant(*7*). These observations highlight potentially conflicting effects of natural selection both within and across immune-mediated diseases, in tension with simple narratives of evolutionary trade-offs or mismatches.

There are multiple directions for future studies. Functional assays(*63*) could help identify more precise biological mechanisms through which our curated fine-mapped candidate variants were adaptive. Infection-related trait GWAS (for example, mapping antibody levels of specific infectious agents, instead of broad syndromic phenotypes like pneumonia) currently have limited sample sizes, and larger sample sizes would shed more light on the relative importance of the selective pressures posed by specific pathogens. Complementary data modalities, such as time-series of ancient pathogen genomes, might be able to elucidate human-pathogen co-evolution. Finally, analyses of ancient DNA data from other locations and epochs will make it possible to assess which of the findings here are particular to the West Eurasian Holocene, and which are more general patterns.

## Acknowledgements

We thank Jordan Rossen, Elizabeth Dorans, Ben Strober, Xilin Jiang, Alison R. Barton, Sarah Fortune, Abigail Frey, Peter Culviner, Xin Wang, Mark Daly and members of the Price and Reich labs for scientific discussions on various aspects of this study. We thank Luke O’Connor for help interpreting LCV results. We thank Nolan Kamitaki and Po-Ru Loh for sharing microbiome and virome GWAS summary statistics ahead of publication. We thank Etienne Patin for sharing antibody reactivity GWAS summary statistics ahead of publication. We thank Karl Heilbron for help running CALDERA. We thank Hilary Finucane and Jacob C. Ulirsch for help running PoPS and interpreting PoPS results. We thank Bokai Zhu for sharing MAXFUSE output from the dataset in ref.(*32*). We thank Liyang Song and Jian Yang for answering questions about how to co-analyze multiple biological replicates using gsMap. We acknowledge the deceased and living individuals whose genetic data made this study possible.

## Author contributions

J.M.-L. and A.L.P. conceived the study. J.M.-L. performed all analyses except the ones conducted by other authors as indicated below, under the supervision of A.L.P. B.T. performed the scDRS analysis. G.K. performed the epigenomic partitioning analysis. Y.Z. performed the PHBC analysis. K.H. performed genome-wide plasma protein QTL mapping. A.P. performed forward-in-time population-genetic simulations. A.A. and D.R. prepared the ancient DNA time-transect dataset and made it available for development of these analyses prior to its public release. A.A. ran the LMM to obtain natural-selection statistics, as well as polygenic selection tests. J.M.-L., A.A., D.R. and A.L.P. interpreted results. A.A., D.R. and A.L.P. supervised different aspects of the study. J.M.-L. and A.L.P. wrote the initial draft. J.M.-L., A.A., D.R. and A.L.P. edited the draft. J.M.-L. prepared the **Supplementary Materials**.

## Data, code and materials availability

LMM selection statistics and selection fine-mapping results are available at https://zenodo.org/records/19563295. Complete numerical results from our analyses are available as **Supplementary Data**. GWAS summary statistics analyzed in this study not generated by the FinnGen consortium are available at https://zenodo.org/records/19563295 and as indicated in the **Materials and Methods** and **Data S1**. GWAS summary statistics generated by FinnGen are available at https://www.finngen.fi/en/access_results. LD Score Regression reference files are available at https://zenodo.org/records/7768714. GTEx cis-QTL summary statistics are available at https://gtexportal.org/home/downloads/adult-gtex/qtl. Immune-cell cis-QTL summary statistics are available at the eQTL catalogue https://www.ebi.ac.uk/eqtl/Data_access/. UK Biobank genome-wide plasma protein QTLs will be publicly released upon publication. Ancient and modern genomes included in the selection scan are available as indicated in ref(*6*). Other data are available as indicated in the **Materials and Methods** or captions of **Supplementary Figures**.

GEMMA v0.98.5 is available at https://github.com/genetics-statistics/GEMMA. SLiM 4 v4.0.1 is available at https://messerlab.org/slim coloc v5.2.3 is available at https://github.com/chr1swallace/coloc. LDSC (LD Score Regression) v1.0.1 is available at https://github.com/bulik/ldsc. LCV is available at https://github.com/lukejoconnor/LCV. PHBC is available at https://github.com/yjzhao1004/pleioh2g. PLINK v2.0 is available at https://www.cog-genomics.org/plink/2.0/. MAGMA v1.10 is available at https://cncr.nl/research/magma/. PoPS v0.2 is available at https://github.com/FinucaneLab/pops. CALDERA is available at https://github.com/kheilbron/caldera/tree/multi. CrossMap is available at https://crossmap.sourceforge.net/. bedtools is available at https://bedtools.readthedocs.io/en/latest/. gseapy v1.1.9 is available at https://github.com/zqfang/GSEApy. statsmodels v0.14.5 is available at https://www.statsmodels.org/stable/install.html. gsMap v1.73.5 is available at https://github.com/JianYang-Lab/gsMap. scDRS v1.0.0 is available at https://github.com/martinjzhang/scDRS. AnnData is available at https://anndata.readthedocs.io. Custom scripts used in our analyses are available at https://github.com/javiermaravall/aDNA_immune_natural-selection.

