## Supplementary Materials for "Ancient DNA reveals that natural selection has upregulated the immune system over the last 10,000 years"

**Supplementary Figures** **38**

**Supplementary Tables** **76**

**Supplementary Data** **84**

**References** **86**

### Materials and Methods

Methodological details for secondary analyses can be found in the captions of the corresponding Supplementary Figures.

Whenever analyses were implemented in custom scripts, we used `Python` v3.12.8(*71*) with libraries `numpy (72)` v2.2.6, `pandas(73, 74)` v2.3.1, `polars` v1.35.2(*75*), `SciPy(76)` v1.16.3, and `matplotlib(77)` v3.10.3(*78*) for plotting; or `R` v4.4.2(*79*) with the library `data.table` v1.17.8(*80*). We used `GPT-4.5(81)` to refine code plotting results or implementing custom analyses.

We often made used of jackknife re-sampling(*82–85*) to obtain standard errors on an estimate.

#### A generalized linear mixed model (GLMM) to detect signals of natural selection using ancient DNA time series data

To test for temporally consistent allele-frequency change while correcting for population structure, Akbari et al.(*86*) used a generalized linear mixed model (GLMM) (see original publication for full details). Briefly, individuals were grouped into ancestry–time clusters to make inference tractable; for cluster $i$ with $n_i$ diploid individuals, the observed allele count at variant $j$ is $y_{ij} \in \{0, \ldots, 2n_i\}$ and was modeled as $y_{ij} \sim Binomial\,(2n_i,\ p_{ij})$. The allele frequency $p_{ij}$ was related to sample age and a structure/drift term using a logit link,

$$logit(p_{ij}) \equiv log\left(\frac{p_{ij}}{1-p_{ij}}\right) = \alpha_j + s_j\, t_i + g_{ij},$$

where $\alpha_j$ is an intercept, $t_i$ is the (negative) sampling date of cluster $i$ expressed in units of twice the generation interval, and $s_j$ is the per-generation selection parameter (assumed constant over the analyzed period). Population structure and correlated drift across space and time were captured by a multivariate normal random effect,

$$\mathbf{g}_j = (g_{1j}, \ldots, g_{Tj})^\top \sim MVN\left(\mathbf{0},\ \sigma_j^2 \mathbf{K}\right),$$

with covariance matrix $\mathbf{K}$ derived from a genetic relationship matrix (GRM). For clusters $m$ and $n$, $\mathbf{K}$ was defined as the mean pairwise relatedness across individuals in the two clusters,

$$K_{mn} = \frac{1}{N_m N_n} \sum_{a \in c_m} \sum_{b \in c_n} A_{ab},$$

where $c_m$ is the set of individuals in cluster $m$, $N_m = |c_m|$, and $A_{ab}$ is the individual-level GRM entry (estimated genome-wide, using a leave-one-chromosome-out scheme to avoid proximal contamination). For each variant $j$, parameters $(\alpha_j, s_j, \sigma_j^2)$ were estimated from the count data; to reduce false positives due to variant-specific heterogeneity in drift/background selection, $\sigma_j^2$ was constrained to be no smaller than a genome-wide, minor-allele-frequency-conditional lower bound. Evidence for selection was assessed by testing $H_0 : s_j = 0$ via a Wald test using $\widehat{s}_j / SE(\widehat{s}_j)$), with calibration and downstream significance assessment as described in ref.(*86*).

#### A linear mixed model (LMM) to detect signals of natural selection using ancient DNA time series data

We determined that GLMM selection statistics were not suitable as input for several downstream methods used in this study because the null distribution of the GLMM test statistics is not normal, largely due to the non-linear link function (Akbari 2026), similar to GWAS mixed-model association for unbalanced binary traits(*87*). To obtain selection statistics that follow statistical assumptions of GWAS methods, we considered a linear mixed model (LMM) in which the response variable is sample date and genotype is a predictor (i.e., the regression direction is reversed relative to allele-count GLMMs). Let $y_i$ denote the standardized sample date sampling time for individual (or cluster) $i$, and let $x_{ij} \in \{0, 1, 2\}$ denote the allele count at variant $j$. We model

$$y_i = \mu + \beta_j x_{ij} + u_i + \varepsilon_i,$$

where $\mu$ is an intercept, $\beta_j$ quantifies association between genotype and sampling time, and $(u_1, \ldots, u_N)^\top$ captures population structure via the GRM:

$$\mathbf{u} \sim MVN\left(\mathbf{0},\ \sigma_g^2 \mathbf{A}\right), \qquad \varepsilon \sim MVN\left(\mathbf{0},\ \sigma_e^2 \mathbf{I}\right),$$

with $\mathbf{A}$ the LOCO GRM and $\mathbf{I}$ the identity matrix. Association is tested via a Wald test for $H_0 : \beta_j = 0$. Intuitively, this LMM uses the same core framework as mixed-model GWAS (a random effect proportional to the GRM to absorb population structure) but targets variants whose frequency has a significant (linear) association through time, rather than directly modeling allele counts through time.

To fit the LMM, we used GEMMA(*88*) (v0.98.5).

#### Assessing LMM selection statistics as input for fine-mapping

##### Simulations with known causal variants

We used SLiM 4(*89*) v4.0.1 for all simulations used in our fine-mapping experiments. Our simulation consisted of user-specified chromosome(s) which were permitted to evolve neutrally for a parametrized "burn-in" period. After the burn-in period, the simulation ran for a user-specified number of generations. During this post-burn-in period, the genomes of a parametrized count of randomly selected individuals were outputted at regular, user-specified increments.

We enabled the user to select a timepoint within the post-burn-in generations at which one or more mutations could have its selection coefficient switched from neutral to positive. If a mutation was chosen for positive selection, its new selection coefficient was sampled randomly from a vector of possible values.

We placed 3 restrictions on the mutations which could be chosen for positive selection. First, to prevent premature fixation of selected variants, the user specified a minimum derived allele frequency (DAF) below which a mutation could not be chosen. Second, the user specified a width, in base pairs, for a chunk of the chromosome to be chosen as a "positive mutation eligible" region. The purpose of this "positive mutation eligible" region was to test whether we could fine-map causal variants when multiple positively selected variants were present in a constrained region. Third, only mutations in "annotated" regions could be positively selected. To define these "annotated" regions, we adjusted a PubMed Genome Data Viewer file (`https://www.ncbi.nlm.nih.gov/gdv/browser/gene/?id=1312`). Prior to running our simulations, we downloaded the NCBI Genome Data Viewer GRCh37.p13 (GCF_000001405.25) files by clicking Download Track Data > NCBI Homo sapiens Annotation Release 105.20220307 for each of the 22 autosomes, then removed all annotations marked as 'pseudogene' and restricted the file to only the annotation names and the start/end base pair coordinates for the annotations. Within the "positive mutation eligible" region, only "annotated" regions were capable of experiencing positive selection, while unannotated regions were only capable of experiencing neutral mutations.

We ran 3 types of simulations: neutral simulations (one for each chromosome for construction of the Genetic Relatedness Matrix), 100 replicates where 1 mutation was chosen to be positively selected, and 100 replicates where 2 mutations were chosen for positive selection. In all our simulations, we used the hg19 recombination rate map (`https://alkesgroup.broadinstitute.org/Eagle/downloads/tables/`), mutation rate of $1.41 \times 10^{-8}$ mutations per base pair per generation(*90*), population size of 5000, 300 post-burn-in generations, and sample size 50 outputted every 10 post-burn-in generations. For all non-neutral replicates, we used a "positive mutation eligible" region size of 3 million base pairs, minimum DAF of 0.01, and the following set of possible positive selection coefficients: 0.001, 0.002, 0.005, 0.01, 0.02, and 0.05.

We ran our GLMM- and LMM-based selection scan on these simulated data. When computing GLMM summary statistics, we deflated $Z$-scores by dividing them by 1.25, to account for inflation in GLMM null selection statistics (see above). This deflation value was estimated using the GWAS enrichment procedure described in detail in SInformation Sections 2 and 3 of Akbari et al.(*86*).

For fine-mapping analyses, we sometimes needed to provide a sample-size value. Apart from the number of simulated genomes included in the scan ($N$), we also computed an "effective sample size" $N_e$ per variant using the GLMM or LMM marginal effect estimate (`beta`), its standard error (`se`), and the minor allele frequency (`MAF`). Assuming an additive genotype coded as $G \in \{0, 1, 2\}$ with allele frequency $p = MAF$ and Hardy-Weinberg equilibrium, the genotype variance is $Var(G) = 2p(1-p)$. Under a single-variant linear regression model with a standardized phenotype (i.e., $Var(Y) = 1$), $Y = \beta G + \varepsilon$, and the residual variance is $\sigma_\varepsilon^2 = Var(Y) - \beta^2 Var(G) = 1 - \beta^2 Var(G)$. The sampling variance of the OLS estimator satisfies $Var(\widehat{\beta}) \approx \frac{\sigma_\varepsilon^2}{N_e Var(G)}$, and we approximate $Var(\widehat{\beta})$ by the reported squared standard error, $se^2$. Solving for $N_e$ yields the per-variant effective sample size estimator

$$N_e = \frac{\frac{1}{Var(G)} - \beta^2}{se^2} = \frac{\frac{1}{2p(1-p)} - \beta^2}{se^2}.$$

We computed this quantity for each variant using either GLMM ($N_{e,\mathrm{GLMM}}$) or LMM ($N_{e,\mathrm{LMM}}$) selection statistics. We note that this derivation is valid for linear regression, such that it is not fully theoretically justified for (generalized) mixed-model association. However, we still evaluated downstream use of these quantities using simulated data.

##### Assessing calibration and power of single-causal-variant fine-mapping of selection loci in simulated data

To assess calibration and power of fine-mapping of selection loci, we performed fine-mapping in simulated data with known favored (causally-selected) variants (see above). Following previous work(*91*, *92*), we divided data for each simulation replicate (a single chromosome) into overlapping 3Mb windows with start points 1Mb apart. To perform fine-mapping on a given window, we required at least one variant in the central sub-region to exceed a genome-wide significance threshold of $P < 5 \times 10^{-8}$. For each retained window, we used an approximate Bayes factor (ABF) framework(*93*) to compute variant-level posterior inclusion probabilities (PIPs) from either (i) per-variant selection effect sizes $\beta$ and their variances $var(\beta) = se^2$, or (ii) minor allele frequency, sample size, and selection $Z-$scores; treating the phenotype as quantitative and setting $sd_Y = 1$ (since sample ages were standardized prior to running the selection scan). Fine-mapping was performed using the function `finemap.abf` in the `coloc(94)` v5.2.3 package, with package defaults. Within each window we recorded per-variant PIPs. Because windows can overlap, the same variant can be assigned PIPs in more than one window; to obtain a non-redundant variant-level output, following ref.(*92*), we retained a single representative record per variant, prioritizing assignments in which the variant lay inside at least one window's central 1Mb sub-region and, among those, selecting the maximum PIP.

For each simulation replicate, we tried several combinations of input parameters: (i) the $P-$value used for window discovery, either coming from the LMM $Z-$scores or the GLMM deflated $Z-$scores; and (ii) the variant-level estimates used for fine-mapping a window, either LMM or GLMM raw effect size estimates with their variances, or GLMM deflated $Z-$scores. When supplying GLMM deflated $Z-$scores, `coloc` required also supplying minor allele frequencies and a sample-size value. We tried 3 different values of sample size: $N$, the number of samples included in the selection scan; $N_{e,\text{GLMM}}$, the effective sample size estimated from GLMM selection statistics (see above); and $N_{e,\text{LMM}}$, the effective sample size estimated from LMM selection statistics (see above). Because LMM selection statistics are less well-powered than GLMM selection statistics (**SI Section 1**), we also tried setting the prior probability of association to $p_1 = 10^{-3}$ when running fine-mapping (larger than the default of $10^{-4}$).

We assessed calibration and power of our selection fine-mapping procedure. We stratified simulations by the maximum non-null selection-coefficient ($S$) value in the replicate and also reported results pooled across all simulations. Because the fine-mapping pipeline was designed to act only on loci passing a genome-wide significance filter, some simulations yielded no fine-mapping output; for each stratum we therefore recorded both the total number of simulations and the subset that produced fine-mapping results. Power metrics were computed using the latter subset as the denominator. For each simulation with fine-mapping results, we extracted the PIP assigned to the single true causal variant and computed power at multiple thresholds $t \in \{0.01, 0.5, 0.9, 0.95\}$ as

$$\widehat{Power}(t) = \frac{1}{M} \sum_{m=1}^{M} \mathbb{1}\left[PIP_{causal}^{(m)} > t\right],$$

where $M$ denotes the number of simulations in the stratum that produced fine-mapping output. Uncertainty was summarized using a leave-one-out jackknife across simulations. For each simulation, we also quantified credible set performance by testing whether the 95% credible set contained the true causal variant. Credible sets were constructed within each window by ranking variants by decreasing PIP and selecting the minimal variant set whose cumulative PIP reached $\geq 0.95$. Credible set power was then defined as the fraction of simulations (among those with fine-mapping output) in which the causal variant was included in its window-level 95% credible set, summarizing uncertainty using a leave-one-out jackknife across simulation. To summarize global error control, we pooled discoveries across all variants and all simulations within a stratum and computed the empirical false discovery rate (FDR) at each PIP threshold $t$ as the fraction of variants with PIP exceeding the threshold that had a null selection coefficient, i.e.:

$$\widehat{FDR}(t) = \frac{\#\{(m,i) : PIP_i^{(m)} > t \ \wedge \ S_i^{(m)} = 0\}}{\#\{(m,i) : PIP_i^{(m)} > t\}},$$

with jackknife standard errors computed by leaving out each simulation in turn. As an additional summary statistic, we computed the mean PIP across all variants and the mean PIP restricted to causal variants (one per simulation) and reported their ratio,

$$R = \frac{\mathbb{E}[PIP \mid S > 0]}{\mathbb{E}[PIP]},$$

again with uncertainty quantified by a leave-one-out jackknife across simulations.

We note that, because `coloc` uses the product of approximate Bayes factors as its main colocalization statistic(*94*), assessment of calibration for fine-mapping purposes extends to assessment of calibration for colocalization purposes.

##### Single-causal-variant fine-mapping of selection loci in real data using coloc

To identify likely causal variants for selection, we performed single-causal-variant fine-mapping of LMM selection association statistics. Following previous work(*91*, *92*), we divided each chromosome into overlapping 3Mb windows with start points 1Mb apart, choosing windows such that all variants belong to the central 1Mb of at least one window. To perform fine-mapping on a given window, we required at least one variant in the central sub-region to exceed a genome-wide significance threshold of $P_{\text{GLMM}} < 5 \times 10^{-8}$, where $P_{\text{GLMM}}$ is the deflated GLMM $P-$value, since the GLMM is better-powered than the LMM for discovery (see above). For each retained window, we used an approximate Bayes factor (ABF) framework(*93*) to compute variant-level posterior inclusion probabilities (PIPs) from per-variant LMM selection effect sizes (which are well-calibrated for fine-mapping according to our simulations; see above) $\beta$ and their variances $var(\beta) = se^2$, treating the phenotype as quantitative and setting $sd_Y = 1$ (since sample ages were standardized prior to running the LMM). Fine-mapping was performed using the function `finemap.abf` in the `coloc(94)` v5.2.3 package. To account for the fact that LMM selection statistics are underpowered, we set the prior probability of association set to $p_1 = 10^{-3}$; we verified this still produced well-calibrated output (**SFigure 6**). Remaining settings followed package defaults. Within each window we recorded per-variant PIPs, and we retained the package's explicit "null" component (no association) as a separate record labeled uniquely by window. Because windows can overlap, the same variant can be assigned PIPs in more than one window; to obtain a non-redundant variant-level output, following ref.(*92*), we retained a single representative record per variant: if the variant lay inside one window's central 1Mb sub-region, we assigned it the corresponding PIP; otherwise, we conservatively selected the smallest of its assigned PIPs (cite).

Because of (i) window start and end base pairs being predefined such that a selection peak may span more than one window, and (ii) window overlap, we constructed 95% credible sets from the fine-mapping PIPs using an iterative, genome-wide seeding procedure within each chromosome. First, we harmonized variant identifiers and genomic positions across window-level outputs; for the fine-mapping "null" records, we assigned the position of the highest-PIP non-null variant in the same window, to maintain a consistent positional representation. At each iteration, we selected a lead (seed) variant from the remaining variants as the one with the largest deflated GLMM $\chi^2$. Around the seed, we defined a $\pm 1.5$ Mb interval (3Mb total) and collected all remaining variants in that interval. Variants in this interval were sorted by decreasing PIP, and we formed a 95% credible set as the minimal set whose cumulative PIP reached at least 0.95. If the total PIP mass in the $\pm 1.5$ Mb interval was $< 0.95$ but at least 0.10, we retained the entire interval as a conservative credible set; if total mass was $< 0.10$, we discarded only the seed variant and continued. After defining a credible set, we removed all variants in its $\pm 1.5$ Mb interval from further consideration to produce non-overlapping credible sets along the chromosome.

We note that our fine-mapping (and colocalization) approaches assumed the existence of at most a single causal variant per locus, despite pervasive evidence of multiple causal variants in disease GWAS(*95*) and anecdotal evidence in this study (Figure S24b). Multiple causal variant methods require a reference LD matrix and are very sensitive to the input matrix(*91*), but it is unclear which LD matrix to use for our selection statistics, given that individuals contribute equally to the LD matrix but likely do not to the selection scan (for example, ref.(*86*) found that older individuals contribute higher statistical power for positive-selection detection). Thus, a standard, unweighted LD matrix is intuitively not the correct choice. Experiments at selected loci indeed suggested a high false positive rate of multiple-causal-variant fine-mapping methods when applied to our selection data (results not shown). Although single causal variant fine-mapping is prone to false negatives, it is not prone to false positives(*91*, *96*). Therefore, we settled on the single-causal-variant assumption as a conservative approach.

Assessing agreement of fine-mapped selected variants with putative favored mutations for well-characterized selective sweeps: To assess agreement of fine-mapping results in real data with previous conclusions from the literature, we collected putative favored mutations from well-characterized selective sweeps from ref.(*97*). For each putative favored mutation, we assessed its PIP and whether it belonged to one of the 95% credible sets produced by our fine-mapping procedure.

Functional enrichment of fine-mapped selected variants: We performed genome-wide functional enrichment analysis of fine-mapped selection variants using the autosomes (chromosomes 1–22), for binary functional annotations coming from (i) the established baselineLD v2.2 model(*98*, *99*) and (ii) a binarized version of the probabilistic consensus variant-to-function (cV2F) annotation(*100*) (with a threshold of 0.75, as recommended by the authors).

Selection variants were annotated with the convention `CHR_BP_REF_ALT` with respect to the GRCh37/hg19 human reference sequence build. To map them to rsIDs (used by our functional annotations), we used dbSNP(*101*) build 156 (GRCh37/hg19), accessed 12 Aug 2024 (`https://ftp.ncbi.nlm.nih.gov/snp/archive/b156/VCF/GCF_000001405.25.gz`), requesting an unambiguous match of all four fields. The variant universe $\mathcal{U}$ was defined as: selection variants that (i) had a single rsID match based on chromosome, base pair, reference and alternative alleles, and (ii) had a non-missing value for all target annotations. The baseline prevalence of category membership was computed as $p_2(c) = \frac{|\{i \in \mathcal{U} : Category_i = c\}|}{|\mathcal{U}|}$, for $c \in \{True, False\}$. Fine-mapping input was obtained as detailed above. Variants in $\mathcal{U}$ without a PIP were assigned a PIP of 0 (since we only performed fine-mapping at genome-wide significant loci for selection). Following ref.(*91*), we evaluated three PIP strata with fixed thresholds: PIP $\geq$ 0.95, 0.5 $\leq$ PIP $<$ 0.95, and 0.05 $\leq$ PIP $<$ 0.5. Within each stratum $r$, we collected the set $\mathcal{F}_r$ of unique annotated fine-mapped variant IDs meeting the PIP criterion, and the stratum-specific category fraction $f_r(c) = \frac{|\{i \in \mathcal{F}_r : Category_i = c\}|}{|\mathcal{F}_r|}$. Functional enrichment was defined as the ratio of the stratum fraction to the baseline prevalence, following previous work(*91*, *102*), $E_r(c) = \frac{f_r(c)}{p_2(c)}$, such that $E_r(True) > 1$ indicates enrichment of annotated variants among fine-mapped variants in stratum $r$ relative to baseline (and analogously for `False`). Uncertainty was estimated via delete-one-block jackknife using $B = 200$ approximately equally-sized genomic blocks of adjacent variants, similar to previous work(*103*). For each block $b$, all variants in that block were excluded by set subtraction from (i) the variant universe $\mathcal{U}$ when recomputing $p_2^{(-b)}(c)$ and (ii) each stratum set $\mathcal{F}_r$ when recomputing $f_r^{(-b)}(c)$, yielding a leave-one-block-out enrichment estimate $E_r^{(-b)}(c)$. The reported enrichment estimate for each $(r, c)$ was the mean of the leave-one-block-out estimates, $\widehat{E}_r(c) = \frac{1}{B}\sum_{b=1}^{B} E_r^{(-b)}(c)$, and the standard error followed the implemented scaling, $SE\left(\widehat{E}_r(c)\right) = SD\left(\left\{E_r^{(-b)}(c)\right\}_{b=1}^{B}\right)\sqrt{B-1}$.

#### Assessing LMM selection statistics as input for stratified LD score regression (S-LDSC)

##### Assessing linear scaling between LD scores and selection statistics

To evaluate whether selection statistics exhibited the approximate linear dependence on linkage disequilibrium (LD) expected by LD score regression (LDSC)(*103*), we assessed the scaling between LMM selection $\chi^2$s, deflated GLMM selection $\chi^2$s, and UK Biobank(*104*, *105*) height GWAS(*91*) $\chi^2$, respectively; and standard LD scores computed from European-ancestry individuals(*99*) from the 1000 Genomes Phase 3 project(*106*). For simplicity, we restricted computations to HapMap3(*107*) SNPs excluding the HLA region, following established practice(*98*, *102*, *103*, *108*).

To test whether the conditional mean relationship between $\chi^2$ and LD score deviated from linearity for any of the three sets of statistics, we compared a linear model against a cubic spline alternative. Let $x = L2$ and $y = \chi^2$. For numerical stability, we standardized LD score within each dataset: $x_s = (x - \mu_x)/(\sigma_x)$. We then specified a restricted cubic spline basis using $K = 10$ knots placed at equally-spaced quantiles between two boundary quantiles of the unstandardized $x$ distribution:

$$q = (0.05, 0.15, 0.25, 0.35, 0.45, 0.55, 0.65, 0.75, 0.85, 0.95), \qquad k_i = Q_x(q_i),$$

where $Q_x(\cdot)$ denotes the empirical quantile function. Knots were transformed to the standardized scale for basis evaluation, $k_{i,s} = (k_i - \mu_x)/\sigma_x$. We used the truncated-power representation of restricted cubic splines, yielding $K - 2$ basis functions corresponding to the $K - 2$ internal knots, and enforced the natural boundary constraints (linear tails beyond the boundary knots). The null (linear) model was

$$\mathbb{E}[y \mid x_s] = \beta_0 + \beta_1 x_s,$$

and the spline alternative was

$$\mathbb{E}[y \mid x_s] = \beta_0 + \beta_1 x_s + \sum_{j=1}^{K-2} \gamma_j \, s_j(x_s),$$

where $\{s_j(\cdot)\}_{j=1}^{K-2}$ are the restricted cubic spline basis functions(*109*). Thus, for $K = 10$ knots, the nonlinearity component has $K - 2 = 8$ degrees of freedom. To quantify uncertainty, we used a block jackknife over $B = 200$ approximately equally-size genomic blocks containing adjacent variants, similar to previous work(*103*). For a model parameter vector $\widehat{\theta}$ estimated on the full dataset, we recomputed the estimator leaving out each block $b \in \{1, \ldots, B\}$ to obtain leave-one-block-out estimates $\widehat{\theta}^{(-b)}$. We estimated the covariance of $\widehat{\theta}$ using the standard jackknife formula

$$\widehat{Cov}(\widehat{\theta}) = \frac{B-1}{B} \sum_{b=1}^{B} \left(\widehat{\theta}^{(-b)} - \bar{\theta}\right)\left(\widehat{\theta}^{(-b)} - \bar{\theta}\right)^{\top}, \qquad \bar{\theta} = \frac{1}{B} \sum_{b=1}^{B} \widehat{\theta}^{(-b)}.$$

We tested for nonlinearity using an omnibus Wald test of the joint null hypothesis

$$H_0 : \ \gamma_1 = \cdots = \gamma_{K-2} = 0,$$

with covariance estimated by the $B$-block jackknife described above. Let $\widehat{\gamma}$ denote the estimated spline coefficients and $\widehat{Cov}(\widehat{\gamma})$ the corresponding jackknife covariance submatrix. The Wald statistic $W = \widehat{\gamma}^{\top} \, \widehat{Cov}(\widehat{\gamma})^{-1} \, \widehat{\gamma}$

was compared to a $\chi^2$ distribution with $K-2$ degrees of freedom to obtain a two-sided $p$-value. We note that this procedure is agnostic to whether deviations from linearity occur preferentially at low or high LD score because the spline basis allows smooth departures across the full support of $x$.

For visualization purposes, we also generated normalized overlays (LMM vs height and GLMM vs height) of binned $\chi^2$–$L2$ relationships (10 $L2$ deciles vs. mean $\chi^2$ within each decile). To place curves on a common scale, we applied an intercept and slope normalization, where we fit the linear model $\chi^2 = \alpha + \beta\, L2$ by ordinary least squares (OLS) and transformed $\chi'^2 = (\chi^2 - \alpha)/\beta$.

###### Assessing concordance of LD scores computed using modern and ancient individuals

We evaluated the concordance between LD scores computed using ancient and modern reference individuals. We split individuals included in our selection scan into 3 sets: (i) those coming from UK Biobank(*104*, *105*) (moderns), (ii) those with a mean age (either from archaeological context or from radiocarbon dating) younger than the median ancient sample age (2175 years before the present), and (iii) those with a mean age older than the median ancient sample age. We computed LD scores for HapMap3(*107*) SNPs (excluding the HLA region)(*103*) for each of these 3 sample sets in turn with respect to variants included in our selection scan (see ref.(*86*) for inclusion criteria), using LDSC(*103*) software v1.0.1 with the `--l2` flag, with the standard genetic-window value of 1cM (`--ld-wind-cm 1`). We regressed the latter onto the former using OLS, and evaluated the Pearson correlation, slope, and intercept of the fitted model.

###### Assessing concordance of LDSC output computed using in-sample and reference LD scores

As an alternative way to assess whether the use of LD scores computed in a modern reference panel was appropriate for our selection statistics obtained from a selection scan including ancient individuals, we ran LDSC v1.0.1 using selection LMM and GLMM statistics, using 1000G European LD scores(*99*) and in-sample LD scores (computed using the union of the 3 sample sets, UKB, younger and older, above). This required weight LD scores apart from regression LD scores. Weight LD scores were computed as above, with the additional flag `--extract` pointing to a list of HapMap3(*107*) SNPs (excluding the HLA region), such that only those variants were included in the computation. We assessed the sensitivity of the 3 main estimates output by LDSC (SNP-heritability, intercept and ratio)(*103*), to LD-score panel choice.

###### Assessing robustness of functional-enrichment estimates to varying sample-size input values

S-LDSC estimates functional enrichment as a relative quantity (the proportion of heritability explained by an annotation normalized by the proportion of variants in that annotation), and is therefore expected to be invariant to the sample-size value supplied as input, provided that the supplied selection statistics obey the modeling assumptions of LD score regression(*102*). We reasoned that sensitivity of inferred enrichments to the sample-size input value likely reflects deviations of the input association statistics from the statistical properties assumed by S-LDSC. To quantify this sensitivity, we ran S-LDSC(*102*) v1.0.1 with a total of 105 functional annotations: those from the baselineLD(*98*) v2.2(*99*) model as well as the cV2F annotations(*100*) (including tissue-agnostic and tissue-specific probabilistic annotations and a binarized version of the tissue-agnostic probabilistic annotation, with a threshold of 0.75 as recommended by ref.(*100*)); for (i) GLMM selection statistics with sample-size input value $N$ and $N_{e,\mathrm{GLMM}}$, and (ii) LMM selection statistics with sample-size input value $N$ and $N_{e,\mathrm{LMM}}$ (see above).

For each pair of runs (fixed selection statistics, differing only in sample-size input value), we summarized the concordance of enrichment estimates across annotations using a scatterplot of enrichment estimates from the $N$-input run versus the $N_e$-input run. We fit an ordinary least-squares regression of $N_e$-based enrichment on $N$-based enrichment and reported the estimated slope (with its standard error) together with a two-sided $P$-value for the null hypothesis that the slope equals 1, which corresponds to perfect invariance of enrichment estimates to the choice of sample-size input. We note that, due to likely non-independence across annotations, OLS standard errors are likely anticonservative.

To test for systematic shifts in enrichment attributable to the sample-size input value, we conducted per-annotation comparisons using the genomic block-jackknife leave-one-out values for functional-enrichment estimates (we modified the software to output these values for each annotation). For each block and annotation, we computed the difference in enrichment estimates between runs, $\Delta_{b,i} = \widehat{Enrich}_{b,i}^{(N_e)} - \widehat{Enrich}_{b,i}^{(N)}$. For each annotation $i$, we estimated the mean difference $\widehat{\Delta}_i$ and its standard error using the standard delete-block jackknife variance estimator,

$$\widehat{Var}_{JK}(\widehat{\Delta}_i) = \frac{B-1}{B}\sum_{b=1}^{B}\left(\Delta_{b,i} - \bar{\Delta}_i\right)^2, \qquad SE_{JK}(\widehat{\Delta}_i) = \sqrt{\widehat{Var}_{JK}(\widehat{\Delta}_i)},$$

where $B$ denotes the number of jackknife blocks and $\bar{\Delta}_i = \frac{1}{B}\sum_b \Delta_{b,i}$. We report per-annotation differences $\widehat{\Delta}_i$, $SE_{JK}(\widehat{\Delta}_i)$, and corresponding nominal $Z$-statistics $Z_i = \widehat{\Delta}_i / SE_{JK}(\widehat{\Delta}_i)$ with two-sided nominal $P$-values computed from the standard normal distribution.

Finally, we implemented a directional omnibus test for a systematic tendency toward higher enrichment under the effective sample-size input, $N_e > N$. For each jackknife block $b$, we computed the cross-annotation mean difference $S_b = \frac{1}{K}\sum_{i=1}^{K}\Delta_{b,i}$ (with $K$ annotations), and used the delete-block jackknife to estimate $\widehat{S} = \frac{1}{B}\sum_b S_b$ and its standard error. A one-sided $t$-test with $B-1$ degrees of freedom was used to test $H_0 : S \leq 0$ against $H_1 : S > 0$. "Flanking" and MAF-bin annotations were excluded from the omnibus test, since they are included in S-LDSC's heritability model mostly to avoid model misspecification, and their enrichment estimates are not usually studied(*98*).

##### Assessing inflation in selection statistics using the LDSC intercept

As an alternative way to evaluate inflation in the GLMM and LMM selection statistics, we evaluated the LDSC intercept. An intercept significantly above 1 reflects inflation not explained by polygenicity(*103*).

##### Comparing S-LDSC—estimated functional enrichment for selection and GWAS

To compare the functional enrichment of selection and GWAS statistics, we ran S-LDSC(*102*) v1.0.1 with the same set of functional annotations indicated above, for each of 74 approximately genetically-independent (pairwise genetic correlation, $|\rho_g| < 0.5$) GWAS traits with publicly-available summary statistics.

We performed a random-effects meta-analysis of S-LDSC enrichment estimates across the 74 traits for the 54 binary and probabilistic annotations out of the 105 analyzed (enrichment estimates for continuous-valued annotations are not interpretable), similar to ref.(*102*). For each annotation, we combined enrichment estimates using a DerSimonian–Laird random-effects model(*110*). Trait-specific inverse-variance weights were defined as $w_i = 1/s_i^2$, where $s_i$ is the enrichment standard error estimated by S-LDSC. The fixed-effect estimate was $\widehat{\mu}_{FE} = \sum_i w_i y_i / \sum_i w_i$, where $y_i$ is the S-LDSC enrichment point estimate. Heterogeneity was quantified by Cochran's statistic $Q = \sum_i w_i (y_i - \widehat{\mu}_{FE})^2$ with degrees of freedom $k-1$ (for $k$ traits).

Between-trait variance was estimated as

$$\widehat{\tau}^2 = max\left(0, \frac{Q-(k-1)}{\sum_i w_i - \frac{\sum_i w_i^2}{\sum_i w_i}}\right),$$

random-effects weights were $w_i^* = 1/(s_i^2 + \widehat{\tau}^2)$, and the random-effects meta-analytic enrichment was $\widehat{\mu}_{RE} = \sum_i w_i^* y_i / \sum_i w_i^*$ with standard error $SE(\widehat{\mu}_{RE}) = \sqrt{1/\sum_i w_i^*}$. Two-sided significance was assessed using $z = \widehat{\mu}_{RE}/SE(\widehat{\mu}_{RE})$ and $P = 2 \times (1 - \Phi(|z|))$, where $\Phi$ is the standard normal cumulative distribution function. Heterogeneity was additionally summarized using $I^2 = max\,(0, (Q-(k-1))/Q) \times 100\%$ and the $\chi^2$ test for $Q$, $P_Q = 1 - F_{\chi^2_{k-1}}(Q)$.

We also ran S-LDSC with the same set of functional annotations, using LMM selection statistics as input.

We compared annotation-specific S-LDSC enrichment estimates obtained from the selection analysis to corresponding random-effects meta-analyzed GWAS enrichment estimates. For each annotation, we extracted the point estimates and standard errors for selection ($\widehat{E}_{sel}$, $SE_{sel}$) and GWAS meta-analysis ($\widehat{E}_{gwas}$, $SE_{gwas}$). We tested for differences between selection and GWAS enrichment estimates using a Wald test assuming independent estimation errors across analyses: for each annotation we computed the difference $\Delta = \widehat{E}_{sel} - \widehat{E}_{gwas}$, its standard error $SE_\Delta = \sqrt{SE_{sel}^2 + SE_{gwas}^2}$, the corresponding $Z$-statistic $z = \Delta/SE_\Delta$, and a two-sided $P$-value $P = 2 \times (1 - \Phi(|z|))$, where $\Phi$ is the standard normal cumulative distribution function; annotations with nominal $P < 0.05$ were highlighted.

To summarize concordance while accounting for measurement error in both axes, we additionally computed a de-attenuated correlation between the underlying (error-free) enrichment effects by correcting the observed cross-annotation covariance for attenuation due to independent estimation error: letting $x_i$ and $y_i$ denote the latent GWAS and selection enrichments, and $\widehat{x}_i = x_i + e_{x,i}$ and $\widehat{y}_i = y_i + e_{y,i}$ their observed estimates with $Var(e_{x,i}) = s_{x,i}^2$ and $Var(e_{y,i}) = s_{y,i}^2$, we estimated $Var(x)$ and $Var(y)$ as the sample variances of $\widehat{x}$ and $\widehat{y}$ minus the mean squared standard errors, and estimated $Cov(x,y)$ by the sample covariance of $(\widehat{x}, \widehat{y})$ (unbiased under independent errors). The de-attenuated correlation was then $\widehat{\rho} = \widehat{Cov}(x,y)/\sqrt{\widehat{Var}(x)\widehat{Var}(y)}$. We obtained a standard error for $\widehat{\rho}$ via a parametric bootstrap(*111*, *112*) ($B = 500$ replicates) by simulating latent pairs $(x_i, y_i)$ from a bivariate normal distribution with mean equal to the observed means and covariance equal to the estimated between-annotation covariance matrix, adding annotation-specific Gaussian noise with standard deviations $(s_{x,i}, s_{y,i})$, and re-estimating $\widehat{\rho}$ in each replicate. Some bootstrap draws produced too little true cross-annotation variation once measurement error was removed, making the de-attenuated correlation undefined. Those draws were dropped from the computation.

#### GWAS data

We collected publicly-available GWAS summary statistics (2,997 traits with summary statistics for HapMap3(*107*) SNPs, suitable for LDSC(*102*, *103*, *108*)-based analyses; 2,564 of which with dense genome-wide summary statistics, suitable for fine-mapping and colocalization analyses), as follows. First, 452 UK Biobank traits from the PanUKBB collection(*113*). Second, 18 UK Biobank traits mapped using BOLT-LMM(*114*, *115*). Third, 61 FinnGen traits related to infection(*116*). Fourth, 122 FinnGen traits corresponding to laboratory values(*116*). Fifth, 694 traits coming from FinnGen-UK Biobank meta-analysis(*116*). Sixth, 330 traits coming from FinnGen-UK Biobank-Million Veteran Program (MVP) meta-analysis(*116*). Seventh, 495 traits with publicly-available GWAS summary statistics curated by the Price laboratory (`https://github.com/TiffanyAmariuta/TCSC/tree/main/sumstats`). Eight, 211 traits from the MiBio-

Gen consortium(*117*), corresponding to microbial abundance at the phylum, order, genus, family and class level. Ninth, 1 trait from ref.(*118*), corresponding to an unsigned composite (cross-species) test for association with oral microbial abundance. Tenth, 106 traits from ref.(*119*), corresponding to allergy, infection, inflammation and serology phenotypes in UK Biobank. Eleventh, 1 GWAS for COVID-19 from ref.(*120*). Twelfth, 41 traits from ref.(*121*), corresponding to genome-wide associations with cytokine (immune-related signaling molecules) levels. Thirteenth, 12 traits from ref.(*122*), corresponding to blood load of several viruses in UK Biobank and All of Us samples. Fourteenth, 163 traits from ref.(*123*) corresponding to antibody reactivity to viral peptides. Fifteenth, 281 traits from ref.(*124*) corresponding to fine-grained parameters of human immune cells. Sixteenth, 2 traits from ref(*125*) corresponding to Tuberculosis (a multi-ancestry and a European-ancestry meta-analysis).

We grouped these traits into 17 non-overlapping categories: Anthropometry, Autoimmune diseases, Behavioral, Biomarkers, Blood-Immune-Inflammatory (mostly blood cell traits), Cardio-Metabolic, Diet and Activity, Immune response (fine-grained mechanisms of immune response, like antibody production), Infection, Inflammatory diseases, Life history-Reproduction, Mental-Psychiatric-Nervous, Microbiome, Pigmentation-UV, Respiratory, Tumors and Other. We tried to closely follow the categories defined by ref.(*86*), adding granularity to immune-related traits. We tried to assign each trait to the single category which best represents it; we acknowledge that some traits could plausibly be assigned to more than one category.

#### Estimating genetic correlation between LMM selection statistics and GWAS association statistics using cross-trait LD score regression (cross-trait LDSC)

We estimated genetic correlations ($\rho_g$) between selection and each GWAS trait using cross-trait LD score regression(*108*) (v1.0.1), which relies on the idea that SNPs with high LD will have higher product of z-scores for two genetically correlated traits than SNPs with low LD. We used the `--rg` option using as input LMM selection summary statistics, and GWAS summary statistics, restricted to HapMap3(*107*) SNPs in both cases. We used the `{{print-delete-vals` flag to output per-genomic-block leave-one-out jackknife estimates of the heritabilities of the two traits and their genetic covariance. LD scores and regression weights were taken from the 1000 Genomes Phase 3(*106*) European reference provided with LDSC (`https://zenodo.org/records/7768714`). We collected each genetic correlation estimate, excluding traits with a non-finite point estimate (likely due to noisy heritability estimates, whose square roots are used in the denominator of $\rho_g$). For each retained trait, we identified significantly non-zero $\rho_g$ by computing a two-sided Wald test statistic $Z = \rho_g/SE(\rho_g)$ and $P = 2 \times (1 - \Phi(|z|))$, where $\Phi$ is the standard normal cumulative distribution function. Multiple testing across traits was controlled using the Benjamini-Hochberg procedure(*126*), and we report the corresponding FDR $q$-values.

#### Meta-analysis of genetic correlations while controlling for covariance in estimation errors

To meta-analyzed estimates of genetic correlation with selection for multiple GWAS traits while accounting for correlated estimation error, we used delete-one-block outputs from cross-trait LDSC(*108*). Specifically, for each trait we used the per-block leave-one-out jackknife (LOO) estimates of the heritabilities of the two traits and their genetic covariance. For each LD block $b$, we reconstructed the LOO genetic correlation as $r_{g,b}^{(-)} = \widehat{gcov}_b^{(-)} / \sqrt{\widehat{h^2_{1,b}}^{(-)}\ \widehat{h^2_{2,b}}^{(-)}}$; blocks yielding non-finite values were excluded, and the remaining blocks were required to be paired across traits (equal numbers of jackknife blocks). Point estimates for each trait were taken as the mean of the corresponding LOO $r_{g,b}^{(-)}$ values. We estimated the cross-trait covariance in genetic-correlation estimation error using the paired-block jackknife covariance matrix of the LOO series: letting $\mathbf{r}_b^{(-)}$ denote the vector of LOO $r_g$ values across the $K$ traits for block $b$, and $\bar{\mathbf{r}}$ the mean across blocks, we computed

$$\widehat{\mathbf{V}} = \frac{B-1}{B} \sum_{b=1}^{B} \left( \mathbf{r}_b^{(-)} - \bar{\mathbf{r}} \right) \left( \mathbf{r}_b^{(-)} - \bar{\mathbf{r}} \right)^{\mathsf{T}},$$

where $B$ is the number of retained blocks.

We then computed a generalized least squares (GLS) meta-analytic genetic correlation using the covariance-aware variance matrix $\widehat{\mathbf{V}}$. The GLS estimate was $\widehat{r}_{g,GLS} = \frac{\mathbf{1}^{\mathsf{T}}\widehat{\mathbf{V}}^{-1}\mathbf{r}}{\mathbf{1}^{\mathsf{T}}\widehat{\mathbf{V}}^{-1}\mathbf{1}}$, with $\mathbf{r} = (r_g^1, \ldots, r_g^K)^{\mathsf{T}}$ denoting the trait-level point estimates and standard error $SE(\widehat{r}_{g,GLS}) = \sqrt{1/(\mathbf{1}^{\mathsf{T}}\widehat{\mathbf{V}}^{-1}\mathbf{1})}$.

This same procedure was used to:

(i) obtain a meta-analyzed estimate of $\rho_g$(Selection, Tuberculosis) using two independent Tuberculosis (TB) GWAS: the European cohort from ref.(*125*) (`Infectious_Tuberculosis_Schurz_et_al_2024_eLife_EUR`) and the best-powered FinnGen(*116*) TB endpoint (`FinnGen_infectious_HG38_TBC_RESP`, which had the largest SNP-heritability $Z-$score ($Z_{h^2}$) among FinnGen TB endpoints)

(ii) obtain a meta-analyzed estimate of $\rho_g$(Selection, Ulcerative colitis) using different cohorts

(iii) obtain a meta-analyzed estimate of $\rho_g$(Selection, Infectious disease), using the following 12 approximately genetically-independent (pairwise $|\rho_g| < 0.5$), well-powered ($Z_{h^2} > 6$) infectious-disease GWAS traits:

```
FinnGen_UKB_MVP_meta_HG38_AB1_VIRAL_HEPATITIS
FinnGen_UKB_MVP_meta_HG38_H7_CONJUNCTIVITIS
FinnGen_UKB_MVP_meta_HG38_H8_MED_SUPP
FinnGen_UKB_MVP_meta_HG38_J10_LARYNGITIS
FinnGen_UKB_MVP_meta_HG38_J10_PERITONSABSC
FinnGen_UKB_MVP_meta_HG38_J10_PNEUMONIA
FinnGen_UKB_MVP_meta_HG38_L12_INFECT_DERM
FinnGen_UKB_MVP_meta_HG38_M13_OSTEOMYELITIS
FinnGen_UKB_meta_HG38_APPENDACUT_NOCOMPLIC
FinnGen_UKB_meta_HG38_K11_GINGIVITIS_PERIODONTAL
Infectious_Kamitaki_et_al_virome_2025_medRxiv_HG38_blood_EBV_invnorm
Infectious_Tuberculosis_Schurz_et_al_2024_eLife_ALL-COHORTS
```

We obtained a $q-$value for the meta-analyzed $\rho_g$(Selection, TB) by obtaining a $Z-$test $P-$value for a significantly non-zero genetic correlation from the meta-analyzed point estimate and SE, and reapplying Benjamini-Hochberg(*126*) together with the previous genetic-correlation $P-$values.

#### A polygenic test to detect significant shits in average genetically-predicted trait values over time

To test whether alleles that are predictive of a trait in present-day European populations have tended, on average, to increase in frequency over time, we followed the established procedure from our previous study(*86*). Briefly, we construct a polygenic score (PGS) for each ancient individual using approximately independent GWAS SNPs. We used the sign-based version of the test developed in ref.(*86*), which only uses the direction of the GWAS effect and is thus less sensitive to misspecifications of the GWAS effect magnitudes. The SNP weight is

$$w_j \;=\; sign(\beta_j) \in \{-1, +1\},$$

where $\beta_j$ is the GWAS effect estimate at SNP $j$. The score for individual $i$ is then

$$PGS_i \;=\; \sum_{j=1}^{M} w_j \, G_{ij},$$

where $G_{ij}$ is the genotype of individual $i$ at SNP $j$ and $M$ is the number of SNPs included.

To estimate temporal change in this sign-based polygenic score while correcting for residual population structure, we fit a linear mixed model

$$y_i \;=\; \alpha + t_i\gamma + g_i + e_i,$$

where $y_i$ is the standardized polygenic score of individual $i$, $t_i$ is sampling time (scaled so that one unit corresponds to 10,000 years), $\alpha$ is an intercept, $g \sim MVN(0, \sigma_g^2 K)$ is a random effect with covariance given by a genetic relatedness matrix $K$, and $e \sim MVN(0, \sigma_e^2 I)$ is residual error. The coefficient $\gamma$ therefore measures the change in the standardized polygenic score over 10,000 years.

The statistic $\gamma_{sign}$ is the estimate of $\gamma$ from this model when the PGS is built with sign-only weights. A significantly nonzero $\gamma_{sign}$ indicates that trait-increasing alleles, defined by GWAS effect direction, show coordinated frequency change through time beyond what is expected from genetic drift and population structure alone. Residual inflation can be calibrated by random sign-flipping of SNP weights, (see ref.(*86*) for full implementation details).

To fit the model, we used GEMMA(*88*) (v0.98.5).

We note that cross-trait LDSC imposes a particularly stringent criterion for statistical significance (a genome-wide consistent pattern as assessed by a genomic-block jackknife), such that this polygenic test may be better powered to detect directional effects of natural selection on genetically-predicted disease risk under a scenario of signal sparsity (*127*) (although we expect cross-trait LDSC and $\gamma_{sign}$ results to be correlated).

#### Single-causal-variant fine-mapping of GWAS loci using `coloc`

We performed single-causal-variant fine-mapping of GWAS loci using `coloc`(*94*) v5.2.3, following the same procedure as indicated above for selection loci.

Fine-mapping windows were defined with respect to the GRCh37/hg19 or GRCh38/hg38 human reference builds depending on the reference build of the source GWAS data. Since selection fine-mapped windows were defined with respect to the hg19 build, we attempted to harmonize genomic window definitions between GRCh37/hg19 and GRCh38/hg38. We constructed a one-to-one correspondence between window identifiers defined in each reference build. For each hg38 window, we computed the midpoint of its central 1 Mbp interval (defined by the table's central-start and central-end coordinates) and represented it as a 1-bp genomic interval. For each hg19 window, we represented the full window interval as a BED feature and additionally computed its midpoint. We mapped hg19 window intervals to hg38 coordinates using UCSC chain files(*128*) and CrossMap(*129*) (`CrossMap bed`), obtaining lifted hg38 intervals for each hg19 window. We then intersected the set of hg38 midpoints (one per hg38 window) with the lifted hg19 window intervals using `bedtools(130) intersect`. This produced, for each hg38 midpoint, a set of candidate hg19 windows whose lifted hg38 interval contained that midpoint. To select a unique best hit when multiple lifted hg19 windows overlapped a given hg38 midpoint, we computed the absolute distance between the hg38 midpoint coordinate and the hg19 midpoint of each candidate window and selected the candidate with minimal midpoint distance. If no lifted hg19 window overlapped an hg38 midpoint, the hg38 window was marked as unmapped.

#### Assessing shared causal variants (colocalization) at loci with evidence of both selection and GWAS association using `coloc`

We tested for colocalization between selection signals and GWAS association signals using `coloc(94)` v5.2.3 within the same set of autosomal 3-Mbp windows we used for fine-mapping (see above). We used GRCh37/hg19 windows for GWAS data in this reference build; GRCh38/hg38 windows otherwise. To perform colocalization, we required both datasets to show evidence of association within the window's central 1Mb sub-region (to ensure the analysis targeted a locally supported signal in each dataset). Specifically, we required at least one variant in the central region with $P_{sel} < 5 \times 10^{-7}$ (where $P_{sel}$ is the deflated GLMM selection $p-$value) and at least one variant in the central region with $P_{GWAS} < 5 \times 10^{-7}$ (we did not require both variants to be the same).

Colocalization was performed using an approximate Bayes factor framework under a single causal variant assumption(*93*), as implemented in `coloc(94)` v5.2.3. We used its function `coloc.abf`. For selection, we used LMM statistics (per-variant LMM effect sizes and variances) as input, and set prior probabilities to `p1 = 1e-3` for association with selection and `p12 = 5e-5` for a shared causal variant to account for the LMM statistics being underpowered (we verified via simulations that this still produced well-calibrated fine-mapping posterior probabilities for selection, see **SFigure 6**); remaining prior settings followed package defaults. We set the selection phenotype standard deviation to 1, since sample ages were normalized before running the LMM (see above). In the case of GWAS input, when effect estimates and standard errors were available, we supplied $\beta$ and $var(\beta)$; otherwise, we supplied two-sided $P$-values. When required by the model parameterization, we additionally provided minor allele frequencies and sample size (taken as the median per-variant sample size within the window). For case–control GWAS, we provided the case fraction when available; for quantitative traits, we provided a phenotype standard deviation when available.

For each analyzed window, we recorded both posterior probabilities for the standard five hypotheses ($H_0$: no association; $H_1$: selection only; $H_2$: GWAS only; $H_3$: two distinct causal variants; and $H_4$: a shared causal variant/colocalization) and a variant-level posterior probability of being the shared causal one (PIP), (conditional on colocalization, such that the product of $H_4$ by the latter gives a per-variant marginal PIP). Within each window we identified the lead colocalizing variant as the variant with the largest conditional PIP. We note that our overlapping-window definition could lead to some variants being assigned a posterior probability by more than one variant (see above). When the same variant appeared in multiple windows, we retained a single representative record, if the variant lay inside one window's central 1Mb sub-region, we assigned it the corresponding PIP; otherwise, we conservatively selected the smallest of its assigned PIPs(*92*) (deduplication). Finally, to filter out redundant colocalizations, we retained a window only if the lead shared-causal variant was identical before and after this deduplication procedure. A colocalization assessment was considered positive whenever $PP_{H_4} > 0.8$, following the original `coloc` publication(*94*).

#### Quantitative-trait-loci (QTL) data

##### Cis-QTLs from the Genome to Tissue Expression (GTEx) project

We downloaded cis-quantitative trait loci (QTL) association summary statistics and fine-mapping results produced by the GTEx Consortium(*131*) from Google Cloud Storage (GCS) buckets. These downloads comprised the complete "all associations" results for GTEx v10 (50 tissues) cis-expression (e)QTL, cis-splicing (s)QTL and cis-alternative polyadenylation (apa)QTL analyses (i.e., genome-wide variant–phenotype association tables stratified by tissue), obtained from the following GCS prefixes:

```
gs://gtex-resources/GTEx_Analysis_v10_QTLs/GTEx_Analysis_v10_eQTL_all_associations/
gs://gtex-resources/GTEx_Analysis_v10_QTLs/GTEx_Analysis_v10_sQTL_all_associations/
gs://gtex-resources/GTEx_Analysis_v10_QTLs/GTEx_Analysis_v10_apaQTL_all_associations/
```

In addition, we downloaded GTEx v8 (49 tissues) RNA-editing (RNA-ed)QTL(*132*) "all associations" results from the public GCS bucket `adult-gtex` under:

```
gs://adult-gtex/bulk-qtl/v8/editing-qtl/all_associations/
```

##### Immune-cell cis-QTLs

We downloaded summary statistics and fine-mapping results for cisQTLs in 5 immune cell types from the eQTL Catalogue(*133*), for the following datasets(*134–142*):

```
Alasoo_2018
Nedelec_2016
Quach_2016
Schmiedel_2018
Randolph_2021
Nathan_2022
Cytoimmgen
Fairfax_2014
Kim-Hellmuth_2017
```

We only downloaded results for the following quantification methods:

- `ge` (gene expression): gene-level expression abundance, typically from RNA-seq. Used to map eQTLs.
- `microarray`: gene expression measured using microarray probe intensities rather than RNA-seq. Used to map eQTLs.
- `leafcutter`: splicing/intron excision phenotypes derived from RNA-seq using LeafCutter(*143*). Used to map sQTLs.
- `txrev`: transcript-structure usage phenotypes (e.g., alternative promoter usage, alternative splicing, alternative 3' end/polyA usage) produced by txrevise(*144*). Used to map transcript-usage QTLs.

We also downloaded fine-mapping results for single-cell expression and infection-response QTLs in leucocytes generated by ref.(*145*) from `https://dataset.owey.io/doi/10.48802/owey.e4qn-9190?version=1.0`.

##### Genome-wide (cis- and trans-) blood plasma QTLs from UK Biobank

Genome-wide pQTL mapping was performed in the UK Biobank Pharma Proteomics Project (UKB-PPP) cohort(*146*), restricted to 36K unrelated individuals of white British ancestry defined by genetic principal component analysis. Protein abundances were inverse-rank normal transformed prior to analysis. We performed association testing using PLINK(*147*) v2.0(*148*) under an additive genetic model, regressing normalized protein levels on genotypes while adjusting for age, sex, and the top 10 genetic principal components. Variants were restricted to a set of 7.7 million well-imputed autosomal SNPs(*149*). To improve cis-pQTL detection power, we regressed out the top 20 principal components computed from the full protein abundance matrix prior to association testing.

#### Assessing shared causal variants (colocalization) at loci with evidence of both selection and QTL association using coloc

We tested for colocalization between selection signals and QTL (either one of the 4 GTEx QTL types, one of the 3 immune-QTL types, or plasma pQTLs) association signals using `coloc`(*94*) v5.2. within the same set of autosomal 3-Mbp windows we used for fine-mapping (see above). We used GRCh37/hg19 windows for QTL data in this reference build; GRCh38/hg38 windows otherwise. Everything was carried out exactly as described for selection-GWAS colocalization assessment (see above), with the exception that GWAS input was substituted for QTL input (treated as a quantitative trait); in all cases, we supplied to `coloc.abf`(*94*) QTL effect sizes with variances, minor allele frequencies (MAF) and sample size (taken as the median per-variant sample size within the window).

#### Fine-mapping of selection genes using CALDERA

##### Using `MAGMA` to obtain gene-level selection $P-$values

We performed gene-based association testing for the selection scan using `MAGMA` (Multi-marker Analysis of GenoMic Annotation)(*150*) v1.10, which aggregates variant-level statistics for variants near each gene. First, we used the option `--annotate` to assign variants to genes using physical position by annotating each variant with overlapping gene coordinates from a reference gene-location file (`https://raw.githubusercontent.com/FinucaneLab/pops/refs/heads/master/example/data/utils/gene_annot_jun10.txt` provided by the `PoPS` GitHub repository, downloaded on July 22 2025). We did not use any window buffer around the genes, following the `PoPS` original publication(*151*). This produced a SNP-to-gene annotation mapping used as input for downstream gene-based tests.

We then used `MAGMA`(*150*) v1.10 to obtain gene-level selection $p-$values. Briefly, for each gene, MAGMA aggregates evidence across all annotated variants in the gene while accounting for linkage disequilibrium (LD) using a LD reference panel. We used flags: (i) `--bfile` to supply in-sample `plink` files for samples in the selection scan(*86*) (cite); (ii) `--pval` to supply variant-level selection $p-$values, as obtained from deflated GLMM results, since GLMM association is better-powered for discovery (see above); (iii) `N=222105`, the fitted effective sample size (see above) ; (iv) `--gene-model snp-wise=mean`, following the `PoPS` original publication(*151*).

###### Using PoPS to prioritize selected genes using polygenic enrichment of gene features

To prioritize selection genes, we used PoPS (Polygenic Priority Score)(*151*) v0.2, which leverages genome-wide patterns of enrichment with respect to gene-level features like bulk and single-cell expression, curated gene pathways, or protein-protein interaction networks; to provide a priority score for genes underlying GWAS loci (in our case, selection loci). We downloaded the PoPS GitHub repo on July 22 2025 (https://github.com/FinucaneLab/pops), as well as reference gene features using the provided link: https://www.dropbox.com/scl/fo/ne7xhxkt4dwhvd52a59ub/AFKkJu7ACaun1uuE99kmTkc/data/PoPS.features.txt.gz?rlkey=ltdbcld1enyr1zefg1lfqm61i. We munged features using the provided script pops/munge_feature_directory.py as indicated on GitHub. We then ran PoPS following options indicated on GitHub and using gene-level selection $P-$values computed by MAGMA(*150*) v1.10 (see above).

###### Using CALDERA to fine-map selected genes

We used CALDERA(*152*) to perform statistical fine-mapping of likely causal genes for selection. Briefly, CALDERA performs a logistic regression using 4 predictors: fine-mapped coding variants at a candidate locus, distance to genes, number of genes at the locus and PoPS(*151*) scores (see above); and 3 covariates encoding local genomic context(*152*). We downloaded the CALDERA GitHub repo on July 25 2025, using the branch indicated by the authors (https://github.com/kheilbron/caldera/tree/multi). We ran CALDERA on R(*79*) v4.4.2 using

```
source(file.path(caldera_path, "z_caldera.R"))
results = caldera(pops_file, cs_file, caldera_path),
```

where caldera_path was our local version of the CALDERA GitHub repo; pops_file was the output PoPS file (see above); and cs_file was a file encoding fine-mapped credible sets for selection (obtained as indicated above), formatted as indicated on the CALDERA GitHub.

#### Gene-set data

Gene-set collections were downloaded from the Human Molecular Signatures Database (MSigDB) database(*153*) v2025.1.Hs (https://www.gsea-msigdb.org/gsea/msigdb/human/collections.jsp), including: (i) 10,480 pathways from gene ontology (GO)(*154*) (c5.go); (ii) 4,023 canonical pathways(*153*) (CP) (c2.cp); (iii) 5,748 pathways from human phenotype ontology(*155*) (HPO) (c5.hpo); and (iv) 50 pathways from the Hallmark collection(*153*) (h).

We also constructed a gene-set collection representing 46 inborn errors of immunity (IEI) gene sets from the International Union of Immunological Societies (IUIS) IEI catalogue (July 2024, version 2(*156*), https://wp-iuis.s3.eu-west-1.amazonaws.com/app/uploads/2024/10/30094653/IUIS-IEI-list-for-web-si xlsx) by grouping the causal genes listed for each disorder. For each catalogue entry, we parsed the reported "Genetic defect" field to extract gene symbols. We then mapped these to Entrez gene identifiers using the NCBI *Homo sapiens* gene information table (http://ftp.ncbi.nlm.nih.gov/gene/DATA/gene_info.gz), prioritizing exact matches to official gene symbols and otherwise using symbol synonyms. Synonyms that mapped uniquely to a single Entrez Gene identifier were accepted; tokens mapping to multiple identifiers were treated as ambiguous and excluded from gene-set construction, and tokens with no match were recorded as unmapped. We grouped mapped genes under each combined “Major category”–“Subcategory” label. For each gene set, we retained the union of Entrez Gene identifiers observed across all contributing catalogue entries and wrote gene sets in MSigDB(*153*) format, with each line containing the set name, a constant descriptor field, and the list of member Entrez Gene identifiers(*157*).

To annotate gene sets as immune-related, we used the curated list from ImmPort(*158*) (`https://docs.immport.org/apidocumentation/genelist/overview/`), which contains gene sets from GO and CP. Analyses that compared immune vs. non-immune were thus restricted to the GO and CP collections.

#### Assessing gene set enrichment among prioritized genes for selection

To assess enrichment of specific gene sets among `CALDERA` selection genes, we assigned to each gene in the background universe (defined by the gene-location file `gene_annot_jun10_LOC.tsv` provided by `PoPS(`*151*`)`, see above) its maximum `CALDERA(`*152*`)` probability of being causal across loci; genes without a `CALDERA` probability were assigned a value of 0 (since selection fine-mapping was only performed at genome-wide significant loci). Gene identifiers in the location file were provided as Ensembl IDs with accompanying gene symbols; we mapped Ensembl IDs to Entrez Gene IDs by stripping any Ensembl version suffixes and using the NCBI correspondece table (`https://ftp.ncbi.nlm.nih.gov/gene/DATA/gene2ensembl.gz`, restricting to taxon 9606 for *Homo sapiens*). We retained only genes with a valid Ensembl→Entrez mapping and collapsed to one record per Entrez gene, keeping the first observed gene symbol as a label. We assessed overrepresentation of GO, CP, HPO, Hallmark and IEI gene sets among genes with CALDERA$> 0.1$, using the full set of genes with assigned CALDERA$\geq 0$ as background universe. Computations were performed using Enrichr, implemented in the python library `gseapy(`*159*`)` (v1.1.9), which uses hypergeometric tests, with arguments `gene_list` (list of genes to test), `gene_sets` (the input collection intersected with the background universe) and `background` (the background gene universe).

To assess enrichment of specific gene sets among colocalized genes, we selected the union of genes attaining $PP_{H_4} > 0.8$ for at least one of the GTEx e,s,apa and RNA-ed QTL colocalization tests. In the case of sQTls and apaQTLs, QTLs were mapped to genes using transcripts; in the case of RNA-edQTLs, QTLs were mapped to genes using the chromosomal location and base-pair position of the RNA editing site, together with a gene-location file listing transcribed regions (supplied by the `MAGMA` software, `https://vu.data.surfsara.nl/index.php/s/Pj2orwuF2JYyKxq`). We defined the GTEx QTL background gene set as the union of genes that were measurable in the corresponding molecular datasets (i.e., present in the corresponding GTEx release summary statistics files, see above) (mapped from Ensembl to Entrez as done for `CALDERA`). Over-representation analysis was performed using Enrichr, implemented in the python library `gseapy(`*159*`)` (v1.1.9), with the same arguments as done for `CALDERA`).

FDR $q-$values were obtained by applying the Benjamini-Hochberg procedure(*126*) to Enrichr $p-$values, across both sets of target genes (`CALDERA`$> 0.1$ and GTEx QTL colocalization) and across all gene-set libraries tested.

##### Assessing likelihood of immune pathways being overrepresented among enriched pathways

We assessed whether immune-related pathways (curated by ImmPort(*158*), see above) were significantly more likely to be significant in our overrepresentation analyses. We performed analyses separately for the collections `CALDERA>0.1` and `UNION_GTEx_QTLs`. For each gene set, we extracted its size from the denominator of the reported overlap statistic (pathway size) and used $log(size)$ as a continuous covariate. We defined significance using the Benjamini–Hochberg(*126*)-adjusted enrichment $q-$values and labeled a gene set as significant if $FDR < 0.10$. To account for gene-set size as a confounder when modeling significance, we fit a logistic regression with a binomial likelihood using `statsmodels(`*160*`)` (v0.14.5). Specifically, for pathway $i$ we modeled the probability of significance as

$$logit\{P(Y_i = 1)\} = \beta_0 + \beta_{immune}\, I_i + \beta_{\log s}\, log(s_i),$$

where $Y_i$ indicates whether the pathway is significant ($FDR_{global} < 0.10$), $I_i$ is an indicator for immune membership, and $s_i$ is the pathway size. We tested the directional hypothesis $\beta_{immune} > 0$ using a one-sided Wald test derived from the fitted coefficient and its standard error, and reported the corresponding odds ratio $exp(\beta_{immune})$. To provide an additional non-parametric assessment of immune enrichment conditional on pathway size that is robust to possible non-independence across gene sets due to overlap in gene content, we performed a permutation-based score test using $B = 10^6$ permutations. We fit a null logistic model without immune status, obtained fitted probabilities $\widehat{p}_i$ and residuals $r_i = Y_i - \widehat{p}_i$, and defined the score statistic as $S = \sum_i I_i r_i$. We computed a one-sided permutation $P$-value, $P_{perm} = (m+1)/(B+1)$, and $m = \#\{S_{perm} \geq S_{obs}\}$ with $S_{perm}$ obtained after randomly permuting the immune labels across pathways.

###### Assessing concordance of gene-set enrichment for GTEx QTL colocalization and CALDERA PIP>0.1 genes

To quantify pathway-level concordance between selection-based and GTEx QTL-based over-representation analyses (ORA), we compared enrichment effect sizes across collections by correlating gene-set odds ratios. Comparisons were made only for the same pathway definition within the same library, and we analyzed effect sizes on the log scale, $log(OR)$, to symmetrize multiplicative effects and stabilize variance. Let $x_i = log(OR_{i,CALDERA})$ and $y_i = log(OR_{i,QTL})$ for the $n$ gene sets tested. We summarized concordance by the Pearson correlation between $x$ and $y$ and reported its analytic $P-$value (as implemented in `scipy.stats.pearsonr(`*76*`)`), which assumes independent observations.

Because the independence assumption likely does not hold (for example, due to gene overlap and hierarchical library structure), we also computed a two-sided permutation $P-$value by breaking the pairing between CALDERA and QTL effects: we repeatedly permuted the ordering of $y$ relative to $x$ ($B = 10^6$ permutations) and recomputed the correlation $r_b$ each time. The permutation $P-$value was computed as

$$p_{perm} = \frac{1 + \sum_{b=1}^{B} \mathbb{1}\left(|r_b| \geq |r_{obs}|\right)}{B+1}.$$

#### Assessing directional impact of selection on pathway activity using genome-wide (cis- and trans-) pQTLs

We restricted analyses to gene sets verifying: (i) they were significant at FDR $<$ 0.10 in our gene-set enrichment analyses (see above); and (ii) they had the same direction of effect on a given biological process, i.e., they were GO(*154*) gene sets consisting of either genes that are all positive or all negative regulators, or Hallmark(*153*) gene sets consisting of either genes that are all upregulated or all downregulated under a particular condition (according to the MSigDB(*153*) definition, `https://www.gsea-msigdb.org/gsea/msigdb/human/genesets.jsp?collection=H`).

For each target gene set, we extracted the union of Entrez gene IDs, and mapped them to Ensembl IDs (see above) to identify them with UK Biobank assayed proteins(*146*) (this limited analyses to the 2794 proteins with genome-wide QTL data, see above). We prepared protein-specific summary statistics by restricting to HapMap3(*107*) SNPs. We estimated genetic covariance between Selection and each pQTL using cross-trait LDSC(*108*), with the same arguments as for GWAS (see above). Genetic correlations were not estimable in general due to noisy heritability estimates for many proteins. We note that very large *cis* effects will have little impact due to LDSC(*103*) automatically filtering out SNPs with extreme association $\chi^2$s. We parsed the reported `Total Observed scale gencov` and its standard error from each LDSC log and retained the accompanying block delete estimates from the `*.gencov.delete` files for jackknife aggregation. We repeated the same computation, but restricting the input SNPs to only those satisfying $|Z_{pQTL}| > T$ and $|Z_{Selection}| > T$, with $T = 3$.

For each pathway, we aggregated per-protein LDSC genetic covariance estimates ($\widehat{gencov}$) across the proteins (genes) belonging to that pathway. We computed (i) an unweighted mean across proteins and (ii) an inverse-variance weighted mean using weights $w_i = 1/SE_i^2$, where $SE_i$ is the LDSC-reported standard error for protein $i$. To propagate uncertainty while accounting for LD Score block structure, we used the LDSC delete-block outputs: for each protein we read the vector of delete-block estimates $\widehat{gencov}_{i,(-b)}$ across blocks $b = 1, \ldots, B$, harmonized proteins to a common block count by removing proteins having a block count different from the modal $B$, and formed pathway-level delete-block estimates by averaging across proteins (unweighted) or by weighted averaging across proteins (weighted). We then computed a standard block jackknife standard error for the pathway-level estimator from the delete-block series,

$$SE_{JK} = \sqrt{\frac{B-1}{B} \sum_{b=1}^{B} \left(\theta_{(-b)} - \bar{\theta}\right)^2},$$

and reported a signed $Z$-score as $Z = \theta / SE_{JK}$, where $\theta$ is the corresponding full-sample pathway-level mean and $\theta_{(-b)}$ is the pathway-level delete-block estimate. We performed this aggregation separately for the baseline LDSC outputs and the $|Z| > 3$ restricted outputs.

We treated the pathway-level weighted $Z-$score for $|Z| > 3$ input as our main test statistic to maximize; for these statistics, we converted pathway-level weighted jackknife $Z$-scores to two-sided $P$-values using the standard normal distribution and controlled the false discovery rate (FDR) at 0.10 across pathways using the Benjamini–Hochberg procedure(*126*). Among pathways passing $q < 0.10$, we additionally required consistent directionality across analyses by enforcing that the signs of the four pathway-level $Z$-scores (baseline unweighted, baseline weighted, $|Z| > 3$ unweighted, $|Z| > 3$ weighted) were all identical; pathways passing this filter as well as $FDR < 0.10$ were considered to be significant results.

##### Assessing likelihood of immune pathways being upregulated by selection

We tested whether pathways that we manually classified as immune-related were more likely than non-immune pathways to show a significant average covariance with selection indicating increased activation. Each significant pathway was encoded as a sign $s_i \in \{-1, +1\}$ based on the sign of its average genetic covariance with selection. We then defined an "increased-activity rule" $r_i \in \{-1, +1\}$ from the pathway identifier: for Gene Ontology(*154*) biological process terms beginning with `GOBP_POSITIVE_REGULATION`, we set $r_i = +1$ (increased activity corresponds to a positive sign), whereas for terms beginning with `GOBP_NEGATIVE_REGULATION` we set $r_i = -1$. For Hallmark(*153*) pathways (`HALLMARK_*`) we set $r_i = +1$ except for `HALLMARK_UV_RESPONSE_DN`, for which we set $r_i = -1$ to account for the "DN" ("downregulated") encoding. The observed indicator of directional agreement was defined as $I_i = \mathbb{1}[s_i = r_i]$, which takes the value 1 whenever selected alleles are predicted to increase the activity of the pathway, and 0 in the reverse case.

Immune pathways were identified by keyword matching on the pathway name using a manually-curated list of immune-related substrings (e.g., `IMMUNE`, `CYTOKINE`, `T_CELL`, etc., and selected Hallmark immune programs such as `HALLMARK_INTERFERON_GAMMA_RESPONSE`, etc.). Let $n_{imm}$ and $n_{non}$ denote the numbers of immune and non-immune pathways, and let $k_{imm} = \sum_{i \in imm} I_i$ and $k_{non} = \sum_{i \in non} I_i$ denote the numbers classified as increased within each group. To summarize immune enrichment for increased activity, we computed a log odds ratio using a Haldane-Anscombe correction(*161*, *162*) to handle the possibility of empty cells in the contingency table:

$$log(OR) = log\left(\frac{(a + 0.5)(d + 0.5)}{(b + 0.5)(c + 0.5)}\right),$$

where $a = k_{imm}$, $b = n_{imm} - k_{imm}$, $c = k_{non}$, and $d = n_{non} - k_{non}$.

As a diagnostic reference under a (likely unrealistic) pathway-independence assumption, we additionally computed one-sided binomial tests within immune and non-immune groups (testing $P(I_i = 1) > 0.5$) and a one-sided Fisher's exact test on the $2 \times 2$ table of increased vs. not increased by immune status.

To account for dependence induced by overlapping gene membership across pathways, we constructed dependence blocks using gene content. Let $G_i$ be the set of genes in pathway $i$. We quantified pairwise pathway similarity by Jaccard overlap, $J(i, j) = |G_i \cap G_j| / |G_i \cup G_j|$, (with $J(i, j) = 0$ whenever $|G_i \cup G_j| = 0$) and clustered pathways using complete-linkage hierarchical clustering on the distance $D(i, j) = 1 - J(i, j)$. Clusters were defined by cutting the dendrogram at distance $\leq 1 - \tau$ with $\tau = 0.10$ (i.e., pathways in the same block have gene-set Jaccard similarity at least 0.10 under the complete-linkage criterion). This yielded a collection of approximately-independent blocks of pathways used to constrain permutations.

We performed a block-wise sign-randomization procedure designed to preserve within-block coherence while randomizing directionality across blocks. Let $\mathbf{s} \in \{-1, +1\}^n$ denote the observed sign vector and let $\mathbf{r} \in \{-1, +1\}^n$ denote the increased-activity rule vector. For each permutation $b = 1, \ldots, B$ (with $B = 10^6$ ), we generated a permuted sign vector $\mathbf{s}^{(b)}$ by iterating over blocks and, independently for each block of size $> 1$, flipping the sign of all pathways in that block with probability 0.5 (i.e., multiplying the subvector by $-1$). For each permutation we recomputed $I_i^{(b)} = \mathbb{1}[s_i^{(b)} = r_i]$, the permuted group counts $k_{imm}^{(b)}$ and $k_{non}^{(b)}$, and the permuted log odds ratio $log(OR)^{(b)}$ using the same Haldane-Anscombe correction(*161*, *162*). We then computed one-sided permutation $P$-values for (i) immune bias toward increased activity,

$$p_{imm} = \frac{1 + \sum_{b=1}^{B} \mathbb{1}\left[k_{imm}^{(b)} \geq k_{imm}\right]}{B + 1},$$

(ii) non-immune bias toward increased activity,

$$p_{non} = \frac{1 + \sum_{b=1}^{B} \mathbb{1}\left[k_{non}^{(b)} \geq k_{non}\right]}{B + 1},$$

and (iii) immune enrichment relative to non-immune (a Fisher-like directional test),

$$p_{assoc} = \frac{1 + \sum_{b=1}^{B} \mathbb{1}\left[log(OR)^{(b)} \geq log(OR)\right]}{B + 1}.$$

**Assessing sign concordance of predicted directional effects of selection on pathway activity across paired positive/negative regulators of the same biological process**

Among pathways significant for our activation test, there were 6 gene-set pairs $(a_i, b_i)$, $i = 1, \ldots, 6$, where each pair comprised gene sets annotated as positive and negative regulators of the same (or very similar) underlying biological process. For each pathway $p$, we used a direction-of-effect label $s(p) \in \{-1, +1\}$, with $s(p) = +1$ denoting a positive genetic covariance with selection and $s(p) = -1$ denoting a negative covariance. We defined an indicator of concordant predicted activity direction as $K_i = \mathbb{1}[s(a_i) \neq s(b_i)]$. This definition reflects the paired construction: when one gene set represents positive regulation and the other represents negative regulation of the same process, opposite signs $s(a_i) \neq s(b_i)$ imply the same predicted direction of pathway activity (either increased activation or decreased activation), whereas equal signs imply discordant activity predictions. The observed test statistic was the number of concordant pairs, $T_{obs} = \sum_{i=1}^{m} K_i$, so that large values of $T_{obs}$ indicate stronger concordance of implied activity direction across the prespecified pairs.

To account for dependence among pathway-level sign assignments induced by overlapping gene membership, we constructed pathway blocks using pathway gene content. For each pathway $p$, we defined $G_p$ as the set of genes annotated to that pathway (parsed as a set of Entrez(*157*) identifiers). Using the Jaccard similarity $J(p, q)$ defined above, we formed an undirected graph on the pathways appearing in the $m$ pairs, adding an edge between pathways $p$ and $q$ if $J(p, q) \geq \tau$, where $\tau = 0.10$. Exchangeability blocks were defined as the connected components of this thresholded gene-overlap graph. By construction, singleton components are allowed and correspond to pathways that have no other pathway with gene-set overlap exceeding the threshold; under the conditional randomization scheme below, such singleton blocks remain fixed.

We performed a one-sided permutation test using $B = 10^6$ permutations with a fixed random seed. Under the null hypothesis of no association between pathway identity and sign label beyond that induced by gene overlap, we generated permuted sign assignments by shuffling the multiset of signs within each exchangeability block. Specifically, for each permutation $b = 1, \ldots, B$, we independently permuted the sign labels among pathways within each block, thereby preserving, for every block, the number of pathways labeled $+1$ and $-1$ while breaking any association between specific pathways and their sign labels within the block. Let $s^{(b)}(p)$ denote the permuted sign assignment, and define

$$T^{(b)} = \sum_{i=1}^{m} \mathbb{1}\left[s^{(b)}(a_i) \neq s^{(b)}(b_i)\right].$$

We computed a one-tailed permutation $P$-value for enrichment of concordant predicted activity direction across pairs (analogous to a one-tailed Binomial test) as

$$p = \frac{1 + \sum_{b=1}^{B} \mathbb{1}\left[T^{(b)} \geq T_{obs}\right]}{B + 1}.$$

#### Defining and annotating clusters of gene features selected by `PoPS` for adaptation

To biologically interpret the gene features selected by `PoPS` for adaptation (see above), we followed the same procedure as in the original publication(*151*) to obtain orthogonal feature clusters, which ref.(*151*) showed are informative about trait-specific biology: e.g., top clusters for LDL cholesterol highlighted lipid synthesis and liver expression; top clusters for schizophrenia highlighted synaptic assembly and calcium channels. For selected features, we constructed a gene-by-feature matrix and standardized each feature to zero mean and unit variance across genes, and we embedded features into a low-dimensional representation using truncated singular value decomposition (SVD) applied to the standardized matrix. Specifically, we fit truncated SVD with $K = 50$ components and represented each feature by its $K$-dimensional loading vector (i.e., the feature coordinates in SVD space). We then computed the pairwise Pearson correlation $r_{jk}$ between feature loading vectors and defined a redundancy-based distance $d_{jk} = 1 - r_{jk}^2$, so that perfectly (anti-)correlated features have $d_{jk} = 0$ and uncorrelated features have $d_{jk} \approx 1$. Features were clustered using complete-linkage hierarchical clustering on the distance matrix, and clusters were defined by cutting the dendrogram at distance threshold $d \leq 1 - \rho$, where $\rho = 0.01$ is an $r^2$ cutoff. This yields clusters such that, under the complete-linkage criterion, all features within a cluster have pairwise squared-correlation at least $\rho^2$ in the SVD embedding. The output of this step was a mapping of each selected feature to a cluster identifier. Values of $K$ and $\rho^2$ followed the original `PoPS` publication(*151*).

To quantify the contribution of each feature clusters to `PoPS` scores, we again followed the original `PoPS` publication(*151*). We first selected one prioritized gene per locus as the gene with the maximum `PoPS` score among genes assigned to that locus. We then reconstructed a standardized gene-by-feature matrix for the union of clustered features (using the same zero-mean, unit-variance scaling across genes) and extracted the `PoPS` marginal feature weights $\beta_f$ for the same feature set. For each feature cluster $c$, we computed a signed cluster contribution across prioritized genes as

$$C_c = \sum_{g \in \mathcal{G}} \sum_{f \in c} z_{gf} \, \beta_f,$$

where $\mathcal{G}$ denotes the set of prioritized genes, $z_{gf}$ is the standardized value of feature $f$ in gene $g$, and $\beta_f$ is the marginal `PoPS` weight for feature $f$. We ranked clusters by $C_c$ (largest to smallest) and summarized relative importance by the fractional contribution

$$F_c = \frac{C_c}{\sum_{c'} C_{c'}}.$$

For the top 25 ranked clusters, we manually examined features belonging to each cluster to produce a biologically meaningful label. Most clusters were unambiguous (e.g., all features referred to expression in the same tissue or cell type). 8 of the 25 clusters consisted fully of features annotated as `human_airway` (data coming from(*163*, *164*)), but without human-interpretable names. We failed to retrieve feature descriptions from the accompanying website (`https://www.genomique.eu/cellbrowser/HCA/`). For this case, Jacob C. Ulirsch (co-first author of the `PoPS` method(*151*)) advised using the pre-defined (`pre_def`) gene features as the basis for annotation. To do this, we downloaded a file of marker genes for human airway cell clusters (`https://www.genomique.eu/cellbrowser/HCA/HCA_airway_epithelium/markers.tsv`) (we restricted to `pvals_adj`<0.05). We performed weighted gene-set enrichment analysis of the `pre_def` features in each target cluster (4 of the 8 clusters contained some `pre_def` feature), with weights given by feature values after standardizing the feature to mean 0 and variance 1, against the marker gene sets. This was done to assess whether the `pre_def` gene sets were enriched for marker genes of specific airway cell types. Enrichment was evaluated using a pre-ranked Gene Set Enrichment Analysis(*165*, *166*) (GSEA) framework implemented in `GSEApy`(*159*) (`prerank`). GSEA was run with 1,000 permutations (`permutation_num`=1000). Gene set size thresholds were set to `min_size`=1 and `max_size`=100000 to retain small marker sets. For each (`PoPS` feature column, cluster) pair, we computed the background size as the number of genes with non-missing scores in that column and the overlap size as the number of marker genes for that cluster present in the background. In addition to the FDR reported by `GSEApy`, FDR

$q-$values were computed within each `PoPS` column by applying the Benjamini–Hochberg procedure(*126*) to the nominal $p-$values across all tested clusters (`statsmodels.stats.multitest.multipletests`, `method='fdr_bh'`). 3 of the 25 clusters were not homogeneous enough to be described by a single label, so were left unannotated. Full details on feature-cluster label choice are given in **SData 5**.

#### Identifying tissues enriched for signals of selection using polygenic enrichment of specifically-expressed genes (LDSC-SEG)

To identify critical tissues for adaptation, we tested whether signals of selection were enriched in regions near genes that are specifically expressed in diverse tissues using the S-LDSC(*102*) with European-ancestry tissue-specific annotations derived from gene expression (LDSC-SEG)(*167*) and the `--h2-cts` flag. Briefly, LDSC models the selection LMM statistic at each variant as a linear function of its LD scores with respect to a baseline set of functional annotations and an additional tissue-specific annotation, and tests whether the tissue-specific annotation adds significantly more predictive power conditional on the baseline model.

We used the precomputed multi-tissue gene expression annotation compendium (`Multi_tissue_gene_expr`) (`https://console.cloud.google.com/storage/browser/broad-alkesgroup-public-requester-pays/LDSCORE/LDSC_SEG_ldscores`), which contains per-tissue specifically-expressed-gene annotations derived from a combination of GTEx(*131*) (human) and Franke lab(*168*, *169*) (human, mouse, rat) expression datasets, together with a matched "all genes" control annotation for each tissue. For each tissue-specific model, the regression included functional annotations from the baseline heritability model(*102*) (not the baselineLD model(*98*), following best practices for critical-tissue discovery(*170*)); the reported tissue-specific enrichment statistic was the regression coefficient ($\tau$) for the specifically expressed gene annotation, with its standard error estimated by the LDSC block jackknife.

Statistical significance for each tissue was assessed using the standard LDSC-SEG one-sided $Z-$test for the tissue-specific regression coefficient being greater than zero, and we applied the Benjamini-Hochberg procedure(*126*) to estimate FDR across tissues. All regressions used European-ancestry 1000 Genomes Phase 3(*106*) reference LD scores and the recommended regression weights (`https://zenodo.org/records/7768714`).

#### Assessing spatial patterns of selection-signal enrichment in the human intestine using `gsMap`

To assess spatial patterns of enrichment of selection signals in the human intestine, we used `gsMap` (genetically informed spatial mapping of cells for complex traits)(*171*) v1.73.5, which integrates spatial transcriptomics with GWAS summary statistics (in our case, LMM selection statistics) to quantify trait relevance for each spatial spot/cell and to produce spatially resolved maps of trait-associated cells. Briefly, `gsMap` denoises spot expression using homogeneous neighbors (via a graph neural network) to compute spot-level gene specificity scores, maps these scores to nearby SNPs (with a $\pm$50 kb window or optionally using epigenomic SNP-to-gene links) to construct spot-specific SNP annotations, and tests heritability enrichment per spot using S-LDSC (conditional on baseline annotations), with optional cross-spot aggregation via a Cauchy combination test(*172*).

We used a single-cell transcriptomic and expression dataset across 8 anatomical regions of the human intestine(*173*). Because ref.(*173*) produced data that did not pair spatial and expression measurements within the same cell, we followed guidance from the authors to integrate both data modalities using the output of their software `MaxFuse` (*174*), a computational framework to perform this matching. The authors supplied `MaxFuse` output for the ref.(*173*) dataset. We constructed one `.h5ad` object per anatomical region by assigning an RNA expression profile to each CODEX spatial observation using the `MaxFuse` correspondence table. Briefly, we used three input files supplied by the authors: (i) a `MaxFuse` mapping table linking CODEX rows (spots) to RNA profile rows, (ii) a CODEX metadata table containing a region identifier and additional cell-level annotations, and (iii) an RNA count matrix (rows as RNA profiles, columns as genes). The mapping table provided integer indices `data2` (0-based CODEX row index) and `data1` (0-based RNA row index), allowing direct alignment between modalities. We retained only CODEX spots with an assigned RNA profile index `data1`. Next, for each unique anatomical region, we subset to CODEX spots assigned to that region, extracted their RNA indices `data1`, and sliced the global RNA matrix to obtain a spot-by-gene count matrix for that region. We created an `AnnData` object with `X` set to the sparse counts and `var` indexed by gene identifiers (columns of the RNA table). We filled in `obs` with spot-level annotations (e.g., tissue segments and cell type labels) and spatial coordinates. Missing annotation values were encoded as `NA`. Spatial coordinates were stored in `obsm["spatial"]`, and the raw matched expression counts were additionally stored in `layers["count"]`. Each region-specific `AnnData` object was written as a `.h5ad` file. This matches the data format required by `gsMap` (`https://yanglab.westlake.edu.cn/gps_data/website_docs/html/data_format.html`).

Because multiple samples were available for each of the 8 intestinal regions, we followed the `gsMap` recommendation of using a "slice mean" analysis to ensure more consistent results across samples (cite `https://yanglab.westlake.edu.cn/gps_data/website_docs/html/advanced_usage.html`). We first used `gsMap` to compute a `slice-mean` object for each anatomical region by aggregating the region-specific `AnnData` inputs across samples. For a given region, we collected all `.h5ad` files matching that region across samples and invoked `gsmap create_slice_mean` using the `count` layer as input. This produced a region-specific slice-mean output that served as the shared reference profile for subsequent per-sample `gsMap` analyses within that region.

We next ran `gsMap` in `quick_mode` separately for each sample-by-region `.h5ad` object and for each cell-level annotation (`obs`) of interest (tissue segment, cell type, etc.). For each run, we provided: the sample-specific `.h5ad` file, the corresponding region slice-mean Parquet file, the target annotation field in `obs`, and LMM selection statistics. We used the default "resource" directory supplied with the installation of `gsMap`.

Finally, we used `gsmap run_cauchy_combination` to aggregate results across samples, both for each anatomical region and across all anatomical regions. We did this for each cell-level annotation (from `obs`) of interest.

#### Identifying cell types enriched for signals of selection using **scDRS**

We performed polygenic disease enrichment analysis in snRNA-seq data using Single-cell Disease Relevance Score (scDRS)(*175*) v1.0.0 together with the snGTEx expression data(*176*). Briefly, scDRS computes cell-level disease (in our case, selection) enrichment scores by aggregating expression of putative disease (in our case, selection) genes derived from GWAS. To obtain putative selection genes, we used MAGMA as indicated above, except we selected a 10 kb mapping window around genes, and we selected the top 1000 genes by MAGMA $P$-value, as previously established(*175*). To calibrate these scores, scDRS generates 1,000 matched "control" gene sets with similar gene-level mean expression and variance, then normalizes the disease scores and derives empirical cell-level $P$ values from the pooled distribution of normalized control scores. scDRS also reports group-level enrichment (e.g., clusters, cell types, regions) using a unified Monte Carlo framework that compares a group's disease-score test statistic to the corresponding distribution from control scores within the same group.

**Investigating correlation between cell-level heterogeneity of selection-signal enrichment and other cell-level features**

Following the original scDRS(*175*) publication, we evaluated heterogeneity test using Geary's C(*177*), a spatial autocorrelation statistic that tests whether cell-level disease scores vary smoothly across neighboring cells in a cell–cell similarity graph. Geary's C significantly below 1 indicates structured within-group heterogeneity, meaning similar cells tend to have more similar disease scores than expected.

### Supplementary Text

#### Supplementary Text A Assessment of summary statistics of a linear mixed model (LMM) to detect signals of natural selection

##### A.1 Summary

We used a different set of genome-wide selection statistics than those computed by Akbari et al(*86*). In detail, Akbari et al.(*86*) computed genome-wide summary statistics for directional selection by applying a generalized linear mixed model (GLMM) to 15,836 West Eurasians who lived over the past 18,000 years and 6,438 contemporary Europeans (Materials and Methods). This GLMM modeled allele count as a function of a time-dependent fixed term and a random term designed to absorb drift and population structure (drawn from a multivariate normal distribution with covariance structure given by the empirical genetic covariance matrix across individuals); their main statistic captured the statistical significance for this time-dependent term being non-zero, thus identifying instances of a temporal trend in allele frequency change not fully explained by drift or structure. This method required the use of a non-linear function to model allele count.

We modified the method of Akbari et al.(*86*) to instead use a linear mixed model (LMM) that regresses sample date on genotypes for each variant while correcting for gene flow and drift (Materials and Methods). The reason for this was that the GLMM selection statistics were not suitable as input for methods developed for GWAS that we apply here, because their null distribution is not normal, mostly due to the use of a non-linear link function(*86*).

We performed 3 experiments to validate our use of LMM selection statistics in this study. First, we confirmed via realistic simulations that LMM selection statistics are normally distributed in the absence of directional selection, unlike those from the GLMM (Figure S1).

Second, we verified that probabilistic fine-mapping(*93*)*18)* (Materials and Methods) of LMM selection statistics produced well-calibrated output in simulations with known causal selected variants, unlike GLMM selection statistics (Figures S2-S6); accordingly, we verified that fine-mapped selected variants in real data largely agreed with putative favored mutations for well-characterized selective sweeps (Table S2), and were enriched for expected functional categories(*91*, *178*, *179*) (Figure S18a).

Third, we showed that LMM selection statistics had similar properties as GWAS association statistics when used as input to stratified LD Score Regression (S-LDSC)(*98*, *102*), unlike GLMM selection statistics (Figures S7,S8,S12-S17). In particular, application of S-LDSC to LMM selection statistics produced the expected functional enrichments (Materials and Methods, Figure S18b), with an estimated correlation (exclusive of noise) of $\hat{\rho}$=0.81 (SE 0.09) with functional enrichments from complex-trait GWAS; non-synonymous (missense) SNPs were more enriched for adaptation than for complex-trait GWAS (26.5x±8x vs. 6.8x±0.7x, $p < 0.02$ for difference), and background selection statistic enrichment closely matched that of complex-trait GWAS, consistent with our previous findings that background selection is not driving positive selection signals(*86*).

Full details are given in subsequent sections of Supplementary Text A.

##### A.2 Assessment of LMM calibration and power

This section evaluates the behavior of linear mixed model (LMM) selection statistics (Materials and Methods) using forward-in-time simulations and compares their performance to the generalized linear mixed model

(GLMM) introduced by (*86*).

Forward-in-time simulations were performed to model human evolution under a realistic European demographic history using the SLiM framework ((**SLIM4**)). The genome was partitioned into three compartments: coding, functional noncoding, and neutral regions. Multiple evolutionary models were simulated, including purifying selection and stabilizing selection, two major processes known to shape patterns of human genetic variation. Each simulation pair included an experimental condition with directional selection and a corresponding negative control that could include purifying selection, stabilizing selection, or both; the two conditions differed only by the presence or absence of directional selection. A detailed description of these simulations is provided in Supplementary Text 2 of (*86*).

In (*86*), a GLMM was applied to estimate selection statistics across the genome (see Materials and Methods for an overview of this methodology). Analyses of these simulations showed that the null distribution of GLMM test statistics is not normally distributed, largely due to the nonlinear link function used in the model (Supplementary Text 2: Non-normality of the null distribution in GLMM test statistics). Under standard assumptions, test statistics are expected to follow an approximately normal distribution under the null hypothesis, and inflation is typically corrected using genomic control ($\lambda_{\mathrm{GC}}$) when only a small fraction of the genome is affected. However, after applying $\lambda_{\mathrm{GC}}$ correction, quantile–quantile (QQ) plots show that the observed P-values become substantially overestimated, indicating clear misspecification of the null model. This behavior, illustrated in Figure S2.12 of (*86*), results in a severe loss of statistical power because the assumption of a normal null distribution does not hold in this GLMM framework.

To address this issue, (*86*) introduced a calibration approach based on enrichment of GWAS signals (Supplementary Information Section 3). Using this approach, the GLMM framework achieves substantially higher power than the linear mixed model (LMM) while maintaining a near-zero false positive rate across all simulated models. In contrast, applying genomic control to the GLMM statistics leads to a severe reduction in power, consistent with the misspecification of the null distribution. Supplementary Table S1 summarizes these results.

Although the LMM is less powerful than the GLMM, its null distribution more closely approximates a normal distribution (Figure S1), particularly for variants with minor allele frequency greater than 0.05 (the standard input for analyses such as polygenic enrichment tests, (*102*)). As a result, LMM-based selection statistics are more appropriate in applications that assume a normal distribution under the null, such as methods from the human statistical genetics literature.

##### A.3 Assessment of LMM selection statistics as input for statistical fine-mapping

We performed forward-in-time simulations with known causal selected variants to assess the calibration and power of fine-mapping of selection signals when using GLMM or LMM selection statistics as input (see Materials and Methods for details).

We evaluation calibration and power of fine-mapping of selection signals when using: (i) deflated GLMM statistics for locus discovery and raw GLMM statistics as input for the fine-mapping method (Figure S2); (ii) deflated GLMM statistics for locus discovery and deflated GLMM statistics as input for the fine-mapping method (Figure S3); (iii) raw LMM statistics for locus discovery and raw LMM statistics as input for the fine-mapping method (Figure S4); (iv) deflated GLMM statistics for locus discovery and raw LMM statistics as input for the fine-mapping method (Figure S5); and (v) deflated GLMM statistics for locus discovery and raw LMM statistics as input for the fine-mapping method, but setting $p_1 = 10^{-3}$ (Figure S6) (the prior probability of there being a causal selected variant at the locus), higher than the default $p_1 = 10^{-4}$, to account for the fact that LMM selection statistics are less well-powered than GLMM selection statistics (Supplementary Text A.2).

We concluded that:

1. Using GLMM selection statistics as input for fine-mapping resulted in miscalibrated posterior probabilities (unexpectedly high FDR for a given PIP threshold).

2. Using deflated GLMM selection statistics as input for fine-mapping partially, but not completely, rescued this miscalibration.

3. Using LMM selection statistics as input for fine-mapping resulted in well-calibrated posterior probabilities, even when using GLMM selection statistics for discovery.

4. Using LMM selection statistics as input for fine-mapping with $p_1 = 10^{-3}$ did not induce miscalibration, even when using GLMM selection statistics for discovery.

Given these observations and the fact that GLMM selection statistics are better powered for selection-locus discovery (Supplementary Text A.2), we decided to use GLMM statistics for discovery and LMM statistics as input for fine-mapping.

###### A.3.1 Assessing agreement between fine-mapped selected variants in real data and putative favored mutations for well-characterized selective sweeps

We performed fine-mapping of selection loci in real data, using GLMM statistics for discovery and LMM statistics as input for fine-mapping, following the conclusions from the previous section (see Materials and Methods for details). Putative favored mutations were collected from a published compendium ((*97*)), and we evaluated each favored mutation belonging to the 95% fine-mapped credible set at the locus.

We report results in Supplementary Table S2. We conclude that 8/11=72% of putative favored variants present in the (*86*) scan belong to the 95% CI at the locus.

##### A.4 Assessment of LMM selection statistics as input for stratified LD score regression (S-LDSC)

See Materials and Methods for details on analyses performed in this section.

###### A.4.1 Assessing scaling between LD scores and selection statistics

S-LDSC expects there to be an approximately linear relationship between LD scores and $\chi^2$ GWAS statistics ((*102*)). We studied the scaling between deflated GLMM selection $\chi^2$s (Figure S7), and raw LMM selection $\chi^2$s (Figure S8); and LD scores. We also compared them to that of a sample GWAS trait (height in the UK Biobank ((*104*, *105*))). We formally assessed a deviation from linearity.

We concluded that GLMM selection $\chi^2$s exhibited strong evidence for a deviation from the S-LDSC expectation, unlike LMM selection $\chi^2$s.

###### A.4.2 Assessing concordance of LD scores computed using modern and ancient individuals

To assess whether LD scores computed in contemporary European-ancestry individuals accurately reflected LD scores in ancient individuals, we compared LD scores computed using European-ancestry samples from the 1000 genomes projects (**Auton2015**, *99*) to LD scores computed using the individuals included in the selection scan performed by (*86*). We split the individuals included in the selection scan into 3 groups: 1) contemporary (from UK Biobank, (*104*, *105*)), 2) ancient individuals with a date more recent than the median sample date among ancient individuals present in the scan, and 3) ancient individuals with a date older than the median sample date among ancient individuals present in the scan; and performed these comparisons separately.

We concluded that LD scores in recent ancient individuals were very highly correlated with LD scores in 1000-genomes samples (Figure S10), with only marginally larger deviation than the one observed for UK Biobank and 1000-genomes individuals (Figure S9). LD scores in older ancient individuals exhibited a small drop in correlation to 1000-genome LD scores, but remained very highly concordant (Figure S11. Our regressions had a slope estimate different from 1, indicating that in-sample LD scores tended to be systematically larger than 1000-genome LD scores. This was not attributable to samples being non-contemporary, since it was also observed for UK Biobank individuals (Figure S9); instead, this suggests that the discrepancy is driven by having used slightly different genetic-variant universes to compute LD scores.

We conclude that pre-computed European-ancestry LD scores are an accurate proxy for ancient LD scores.

###### A.4.3 Assessing concordance of LDSC output computed using in-sample and reference LD scores

As a further evaluation of whether using reference LD estimated from contemporary Europeans was appropriate in the context of an ancient-DNA based selection scan, we studied three key quantities estimated by LDSC, when using LD scores computed using European-ancestry individuals from the 1000 genomes project ((**10002015global**, *99*)), or when using LD scores computed in sample, using the exact same set of individuals that were included in the selection scan performed by (*86*); for both deflated GLMM selection statistics (Figure S12) and raw LMM selection statistics (Figure S13).

We concluded that estimates appeared to be concordant for both sets of input LD scores, with LMM statistics suggestively showing higher agreement.

###### A.4.4 Assessing robustness of functional-enrichment estimates to varying sample-size input values

S-LDSC estimates heritability enrichment for a given set of functional categories, defined (for binary annotations) as the proportion of trait heritability contained in a given annotation, relative to the overall prevalence of that annotation ((*102*)) (we note that, in the context of selection statistics, the proportion of heritability can be interpreted as the proportion of variance in causal selection coefficients following theory that establishes an analogy between individual-level and variant-level definitions of this quantity, see (*102*, *180*)). Because enrichment is a relative quantity, the sample-size value given as input to S-LDSC should not alter functional-enrichment estimates. We thus reasoned that substantial differences in S-LDSC functional-enrichment estimates for selection statistics with different sample-size input values is evidence of the statistics violating the assumptions of the S-LDSC model.

We ran S-LDSC to model deflated GLMM or raw LMM selection statistics, using as input either $N$ (the actual number of individuals included in the selection scan performed by (*86*)) or $N_{e,\text{GLMM}}/N_{e,\text{LMM}}$ (an "effective" sample size estimated from deflated GLMM or raw LMM selection statistics, respectively; see Materials and Methods for details). We computed functional-enrichment estimates for binary or probabilistic annotations in the established baselineLD model ((*98*, *102*)). We found a pattern strongly suggestive of deviation for GLMM (Figure S14), but not LMM (Figure S15) selection statistics.

To formally assess the deviation pattern while controlling for covariance across annotations, we implemented an omnibus test. This test was significant for GLMM (Figure S16), but not LMM (Figure S17) selection statisticsl, with enrichment point estimates growing larger with larger sample-size input values.

We conclude that there is substantial evidence that GLMM selection statistics deviate from the generative model of S-LDSC; this is not the case for LMM selection statistics as assessed by the tests used in this section.

###### A.4.5 Assessing inflation in selection statistics using the LDSC intercept

As an alternative way to assess inflation in selection statistics not explained by the LDSC generative model, we examined the LDSC intercept; this quantity being significantly larger than 1 suggests that inflation in the modeled summary statistics is not entirely attributable to polygenicity ((*103*)).

###### A.4.6 Conclusions

Our results suggest that LMM selection statistics, unlike GLMM selection statistics, behave no differently than GWAS summary statistics with respect to the S-LDSC generative model, and are thus suitable input for polygenic enrichment tests.

#### Supplementary Text B Expanded discussion of selected immune-related genes fine-mapped with `CALDERA`

In Figure 3g, we highlight 10 immune-related genes fine-mapped with `CALDERA`, most exhibiting a positive selection-QTL colocalization, but no GWAS-QTL colocalization. Additional examples are reported in Table S5. We provide more details on 6 of these 10 genes:

(i) *DUOX2* ($\text{PIP}_{\text{CALDERA}}$=0.67, GTEx eQTL colocalization) is an epithelial oxidase that generates reactive oxygen species to support mucosal antimicrobial defense(*181*).

(ii) *MUC4* ($\text{PIP}_{\text{CALDERA}}$=0.55, GTEx eQTL colocalization) is a membrane-bound mucin that contributes to the mucosal glycocalyx barrier and impacts epithelial responses at the host–pathogen interface(*181*).

(iii) *CFTR* (not involving the major cystic fibrosis allele deltaF508(*86*)) ($\text{PIP}_{\text{CALDERA}}$=0.70, GTEx eQTL colocalization) is an ion channel that maintains airway/gut surface hydration and mucus properties critical for mucociliary clearance and antimicrobial protection(*181*).

(iv) *CDHR3* ($\text{PIP}_{\text{CALDERA}}$=0.52, GTEx sQTL colocalization) is an airway epithelial surface protein that serves as a rhinovirus C entry receptor(*181*).

(v) *IFNGR1* codes for the IFN$-\gamma$ (interferon gamma) receptor component 1 ($\text{PIP}_{\text{CALDERA}}$=0.61; monocyte eQTL, GTEx sQTL, and UK Biobank pQTL colocalization). The interferon gamma signaling axis sits at the heart of host immune response to mycobacterial pathogens(*182*, *183*) (the genus containing *M. tuberculosis*) and other intramacrophagic pathogens: IFN$-\gamma$ binding to its receptor on macrophages triggers these cells to clear mycobacteria(*184*). We also identified several selection$-$trans-QTL colocalizations for *IFNGR1*, as well as for *IFNG* itself and *IFNGR2*.

(vi) *EBF1* ($PIP_{CALDERA}$=1.00) is a transcription factor required for B-cell lineage specification and maturation, thereby supporting humoral immune competence(*181*).

Full numerical results are given in Data S3.

#### Supplementary Text C Expanded discussion of `gsMap` results for a human-gut spatial transcriptomics dataset

We applied `gsMap` ((*171*)) to assess patterns of enrichment in a single-cell, spatial transcriptomics dataset across 8 regions of the human intestine ((*173*)). The method computes cell-level enrichment $p$-values, as well as annotation-level enrichment $p$-values via a Cauchy aggregation test (*172*). Because the Cauchy aggregation test is robust to $p$-value dependence at the price of being sensitive to individual outliers within an annotation, it is recommended to only view as significant those annotations that show both small Cauchy and small cross-cell median $p$-values ((*171*)) (see Materials and Methods for details). We highlight several conclusions:

(i) As assessed by this metric, all individual gut regions highlighted the mucosal tissue layer specifically (Figure S29).

(ii) As assessed by this metric, all individual gut regions highlighted immune cells specifically, with the single exception of the Transverse region, where immune cells still had the smallest median $p$ (Figure S30).

(iii) When pooling data across gut regions and stratifying cells by finer spatial neighborhoods, the only neighborhoods exhibiting a robust pattern of significant Cauchy and median enrichment were immune in nature: the follicle, and those with overrepresentation of CD8+ T lymphocytes, adaptive immune cells, and plasma cells (Figure S31).

(iv) When pooling data across gut regions and stratifying cells by cell type (cluster term), the only cell types exhibiting a robust pattern of significant Cauchy and median enrichment were immune cell types: CD4+ and CD8+ T lymphocytes, as well as B and plasma cells (Figure S32).

##### C.1 Consistency between `gsMap` results and other observations

Our analyses using `gsMap` indicated a specific connection between adaptation and immune cells in the gut mucosa. We note that this finding is consistent with several previous observations:

(i) Selection-implicated genes (Figure 3g), disease/traits (Figure 2a,3c) and pathways (Figure 4b) related to gastrointestinal infections and gut mucosal immunity (in particular, ulcerative colitis).

(ii) QTL colocalization and CALDERA implicating core gut-immune genes (Table S5)—consistent with their lumen-facing roles, enrichment strengthens toward the lumen: more apical cells show lower $p-$values (Figures S27c,d).

(iii) Selection-GWAS colocalizations with antibody reactivity to enteric viruses, diseases/traits related to gut mucosal inflammation (e.g. colorectal cancer), and calprotectin in feces (a biomarker of intestinal inflammation(*185*) (Data S3).

(iv) LCV results suggestively linking *H. pylori*, a gastric pathogen, and adaptation (Figure S22b).

(v) Genetic correlation (Figure 2a) and LCV (Figure S22b) results implicating *Akkermansia*, suggesting that selection strengthening the gut mucus layer has led to increased abundance of this commensal, consistent with positively-selected alleles being associated with increased expression of MUC2 (Data S3), a mucin protein that is the main component of this mucus layer.

(vi) Ancient pathogen DNA evidence of an increased burden of gut-invading pathogens, from the Neolithic period onwards(*186*).

#### Supplementary Text D Decomposing the selection–ulcerative colitis association with respect to protection against intestinal infections

##### D.1 Hypothesis

We found positively-selective alleles to be positively correlated with risk for ulcerative colitis (a form of intestinal inflammatory disease) (UC) and negatively-correlated with susceptibility to intestinal infections (Figure 1). This raises the possibility of an evolutionary trade-off through antagonistic pleiotropy.

We assessed whether alleles favored by selection tend to increase UC risk specifically through their effects on protection against intestinal infections.

##### D.2 Analyses performed

Let $S$ denote the LMM selection $Z$-score ($Z_s$; $S > 0$ indicates A1 favored), $U$ the UC GWAS $Z$-score ($U > 0$ indicates higher UC risk), and $Z_{\text{inf}}$ the intestinal infections GWAS $Z$-score (negative indicates protection). Let us recode $T \equiv -Z_{\text{inf}}$ (so $T > 0$ corresponds to infection protection). Let us align all alleles consistently across the three, using the effect allele for selection as the reference.

Using HapMap3 SNPs ((*107*)) with LD-based weights ($w = 1/\max(\text{LD-score}, 1)$) and a block jackknife over LD-ordered SNPs (100 blocks) ((*103*)), we estimated:

(i) weighted genome-wide correlations $\text{cor}(S, U)$ and $\text{cor}(S, Z_{\text{inf}})$

(ii) the weighted regression $U = \alpha + bT + \varepsilon$ (so $b > 0$ implies infection-protective alleles are UC-risk alleles)

(iii) a covariance decomposition of the selection–UC association into an "infection-linked" component and a residual:

$$\text{Cov}(S, U) = \underbrace{\text{Cov}(S, U_T)}_{\text{infection-linked}} + \underbrace{\text{Cov}(S, U_{\text{res}})}_{\text{residual}},$$

where $b = \text{Cov}(U, T)/\text{Var}(T)$, $U_T \equiv b\,(T - \mathbb{E}[T])$, and $U_{\text{res}} \equiv (U - \mathbb{E}[U]) - U_T$. We additionally report the fraction of $\text{Cov}(S, U)$ attributable to the infection-linked component:

$$f_T \equiv \frac{\text{Cov}(S, U_T)}{\text{Cov}(S, U)}.$$

Numerical results are reported in Supplementary Table S8.

##### D.3 Conclusions

Our results indicate that:

1. Genome-wide, selection is positively correlated with UC risk ($\mathrm{cor}(S, U) > 0$), and selection is aligned with infection protection ($\mathrm{cor}(S, Z_{\mathrm{inf}}) < 0$, equivalently $\mathrm{cor}(S, T) > 0$) (as we had previously observed using cross-trait LDSC ((*108*)), see (Figure 1)).

2. Infection protection itself is UC-protective ($\mathrm{cor}(U, T) < 0$ and $b < 0$).

3. The infection-linked component $\mathrm{Cov}(S, U_T) = b\,\mathrm{Cov}(S, T)$ is negative: the part of selection that tracks intestinal infection protection predicts *lower*, not higher, UC risk. At the same time, the residual component $\mathrm{Cov}(S, U_{\mathrm{res}})$ is positive and dominates the net positive selection–UC association. This is the opposite pattern of what an evolutionary-tradeoff scenario would predict.

Thus, the positive genome-wide selection–UC correlation does not arise from selection acting through intestinal infection protection; instead, selection contains at least two components: an infection-aligned, UC-protective component and a larger residual component that increases UC risk.

Our gene, pathway and tissue results have repeatedly implicated selection on stronger mucosal barrier immunity, in particular in the gut, which would be expected to lower, not increase UC risk (intestinal-barrier breakdown is a common trigger of intestinal inflammation, see (*187*)). On the other hand, we have also predicted adaptation leading to increased activity of inflammatory processes. Therefore, it is plausible that the first hypothesized component, protective for both intestinal infections and UC, is related to mucosal barrier immunity; whereas the second hypothesized components, which dominates the genome-wide relationship between selection and UC, is related to the pro-inflammatory effect of adaptation.

### Supplementary Figures

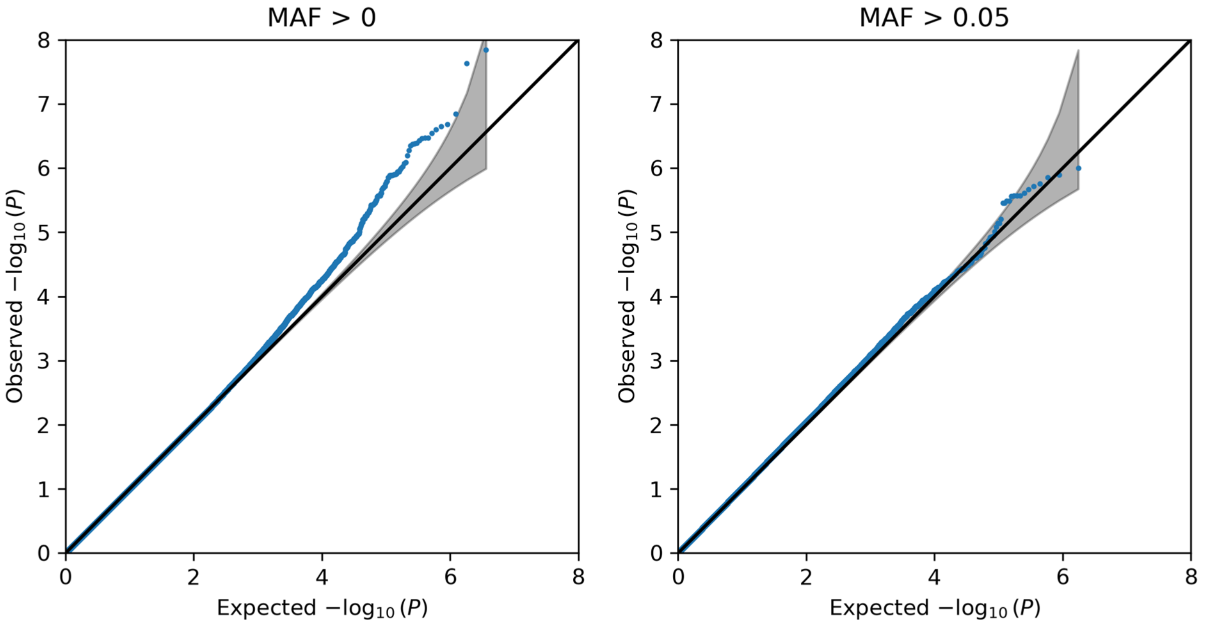

**Figure S1:** Quantile–quantile (QQ) plots comparing observed and expected P-values from LMM test statistics under the null model. Expected values assume test statistics follow a normal distribution under the null. The shaded region shows the 95% confidence band. Null simulations follow Model 2.1 from (*86*) and were performed in SLiM on a random 100 kbp genomic region, producing approximately 3.6 million SNPs from 20,000 replicates without directional selection. The left panel includes all variants (MAF > 0), and the right panel includes variants with MAF > 0.05.

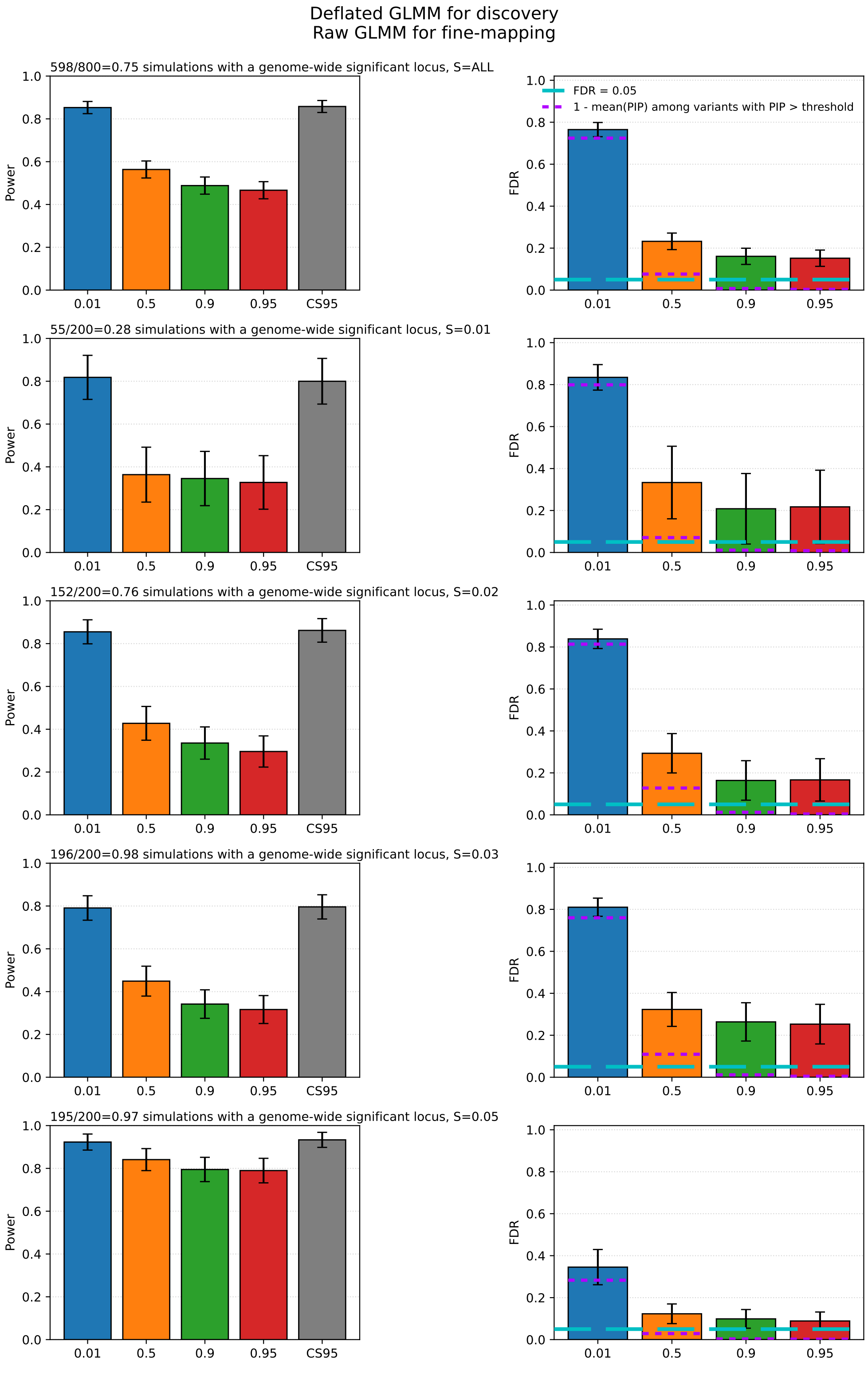

**Figure S2:** Calibration and power of fine-mapping of selection loci, using deflated GLMM selection statistics for discovery (i.e., to decide whether to run fine-mapping at a given locus) and raw GLMM selection statistics as input for the fine-mapping pipeline. We show power (including for belonging to the 95% credible set, CS95) and FDR, stratifying by value of $s$ (selection coefficient, different rows) and for variants with PIP above a given threshold $t$ ($X-$axis ticks); we plot the value across all simulations (plot estimate) and 1.96 $\times$ SE, computing standard errors (SEs) via a leave-one-simulation-out jackknife. For FDR, we indicate both 5% and 1-[average PIP for variants with PIP $>$ $t$], which should approximate the true FDR under well-calibrated posterior probabilities. See Materials and Methods for full details.

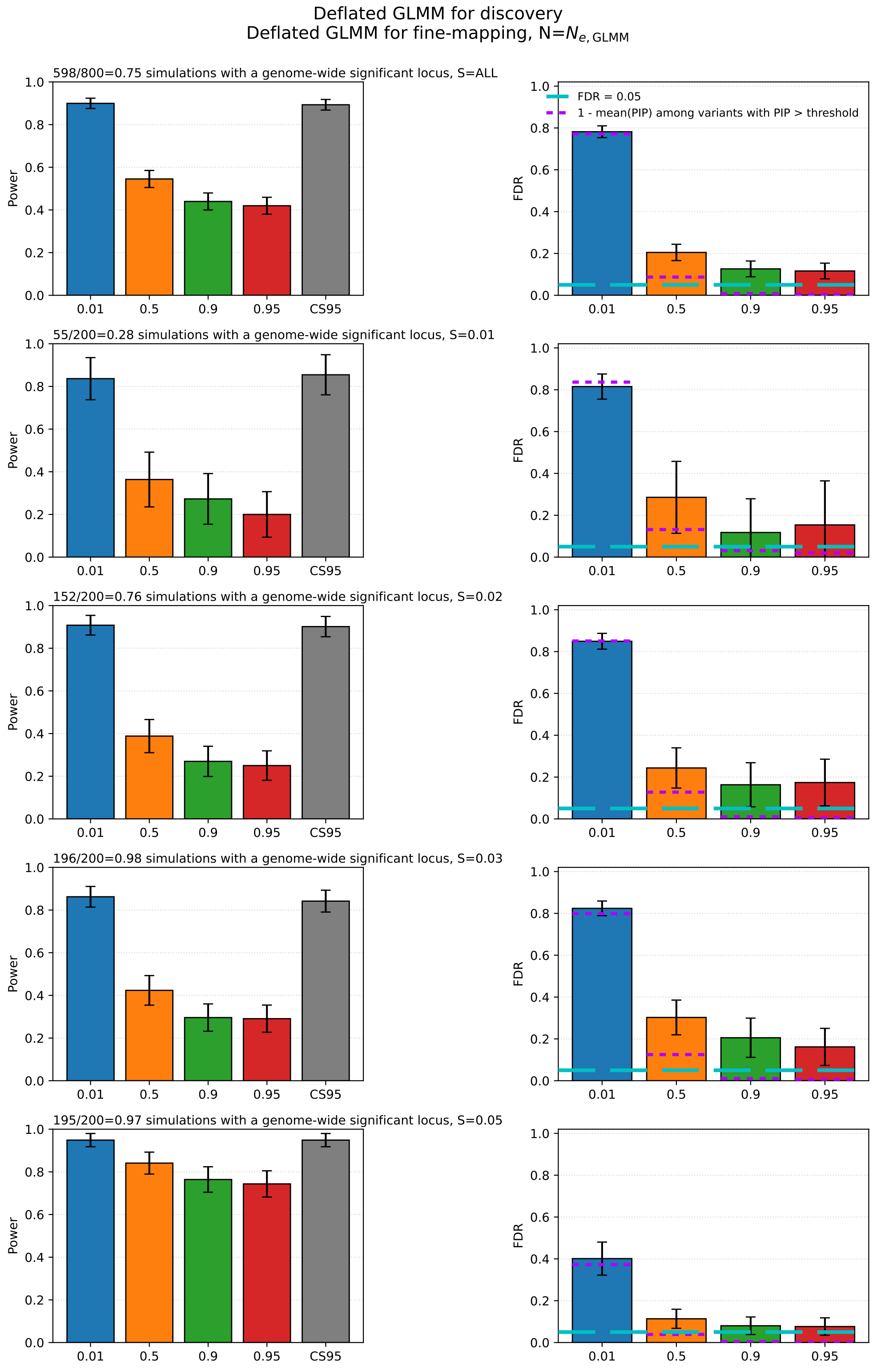

**Figure S3:** Calibration and power of fine-mapping of selection loci, using deflated GLMM selection statistics for discovery (i.e., to decide whether to run fine-mapping at a given locus) and deflated GLMM selection statistics as input for the fine-mapping pipeline. We show power (including for belonging to the 95% credible set, CS95) and FDR, stratifying by value of $s$ (selection coefficient, different rows) and for variants with PIP above a given threshold $t$ ($X-$axis ticks); we plot the value across all simulations (plot estimate) and 1.96 $\times$ SE, computing standard errors (SEs) via a leave-one-simulation-out jackknife. For FDR, we indicate both 5% and 1-[average PIP for variants with PIP $>$ $t$], which should approximate the true FDR under well-calibrated posterior probabilities. See Materials and Methods for full details.

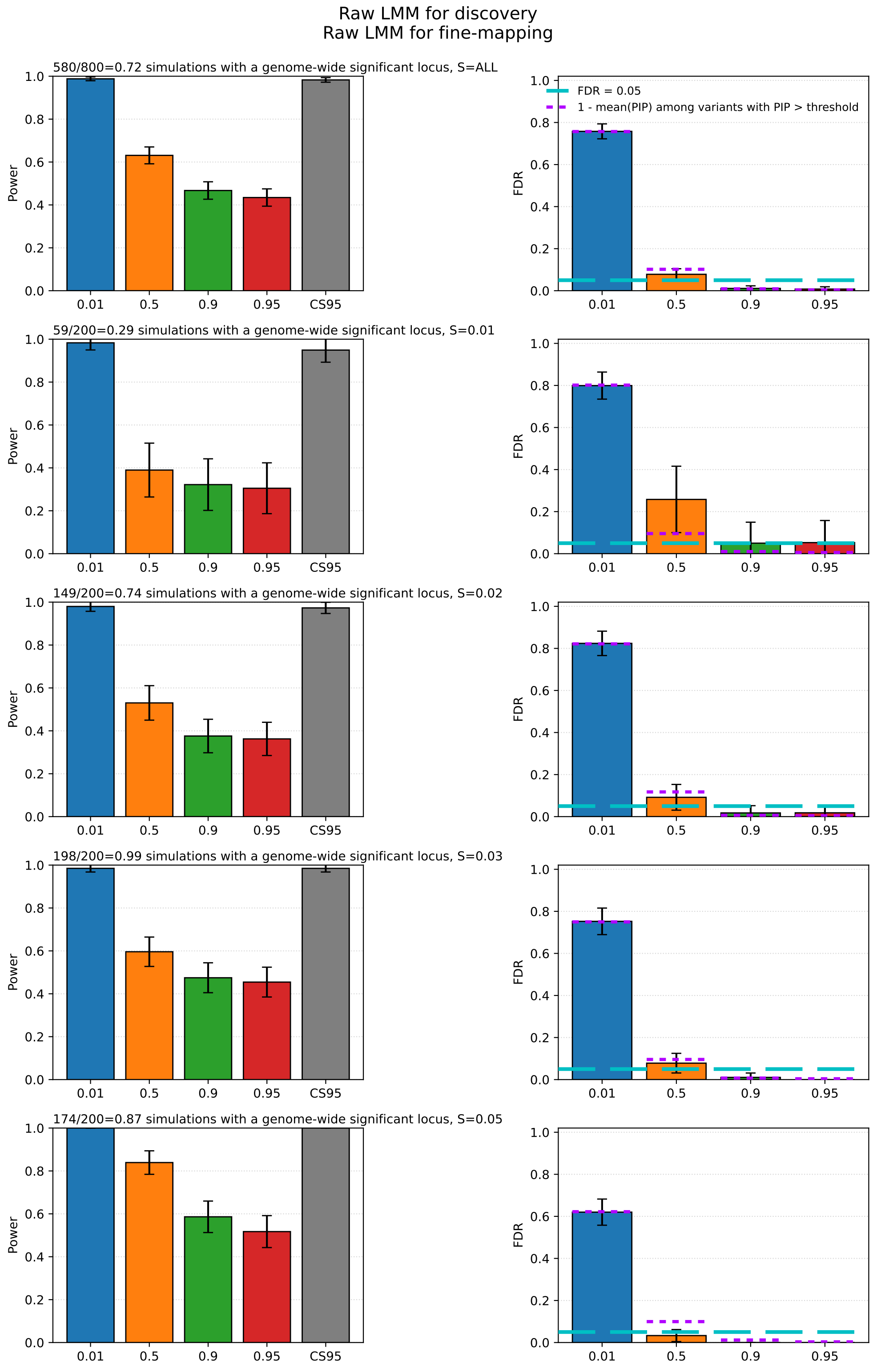

**Figure S4:** Calibration and power of fine-mapping of selection loci, using raw LMM selection statistics for discovery (i.e., to decide whether to run fine-mapping at a given locus) and raw LMM selection statistics as input for the fine-mapping pipeline. We show power (including for belonging to the 95% credible set, CS95) and FDR, stratifying by value of $s$ (selection coefficient, different rows) and for variants with PIP above a given threshold $t$ ($X-$axis ticks); we plot the value across all simulations (plot estimate) and 1.96 $\times$ SE, computing standard errors (SEs) via a leave-one-simulation-out jackknife. For FDR, we indicate both 5% and 1-[average PIP for variants with PIP $> t$], which should approximate the true FDR under well-calibrated posterior probabilities. See Materials and Methods for full details.

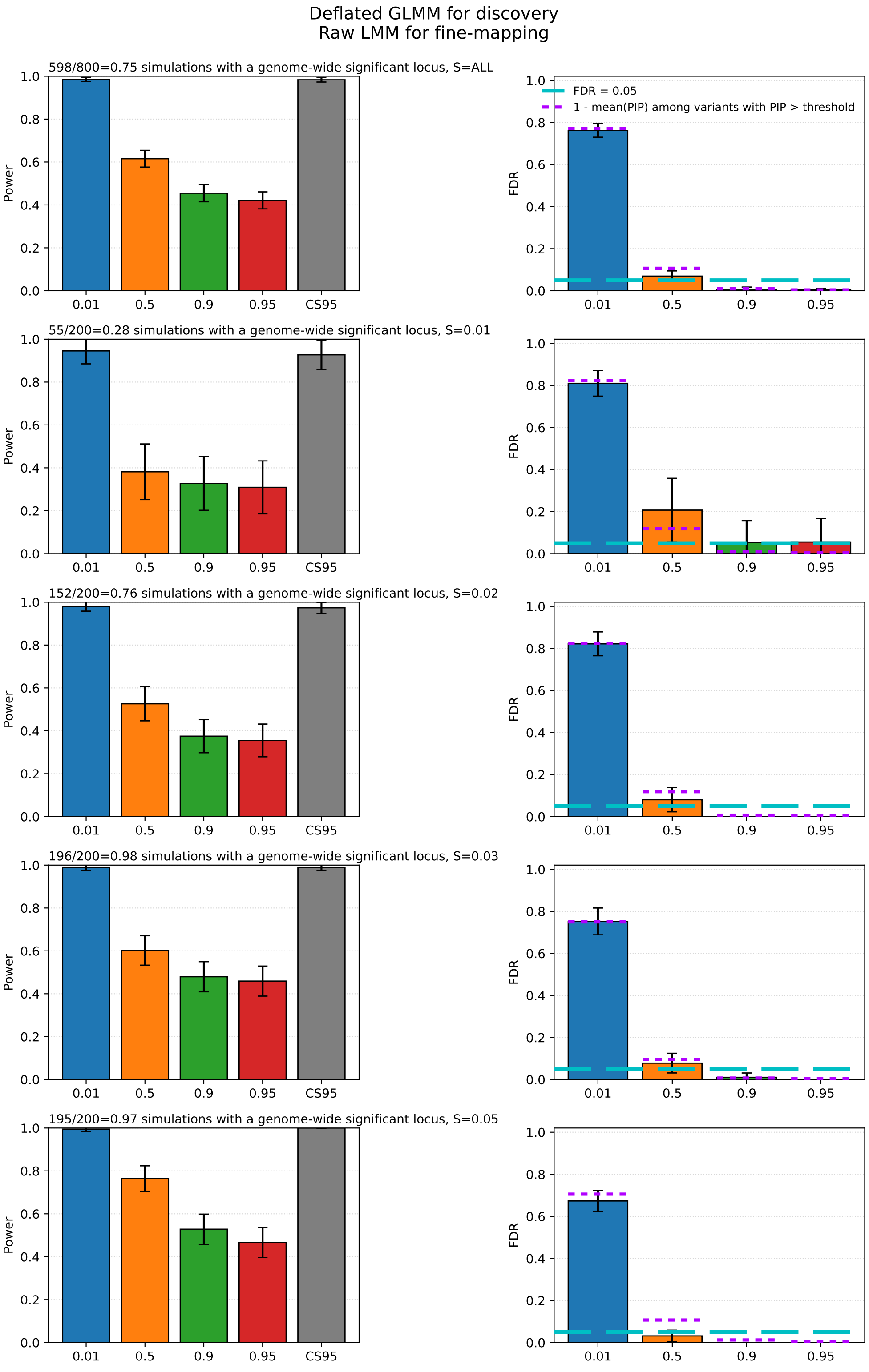

**Figure S5:** Calibration and power of fine-mapping of selection loci, using deflated GLMM selection statistics for discovery (i.e., to decide whether to run fine-mapping at a given locus) and raw LMM selection statistics as input for the fine-mapping pipeline. We show power (including for belonging to the 95% credible set, CS95) and FDR, stratifying by value of $s$ (selection coefficient, different rows) and for variants with PIP above a given threshold $t$ ($X-$axis ticks); we plot the value across all simulations (plot estimate) and 1.96 $\times$ SE, computing standard errors (SEs) via a leave-one-simulation-out jackknife. For FDR, we indicate both 5% and 1-[average PIP for variants with PIP $> t$], which should approximate the true FDR under well-calibrated posterior probabilities. See Materials and Methods for full details.

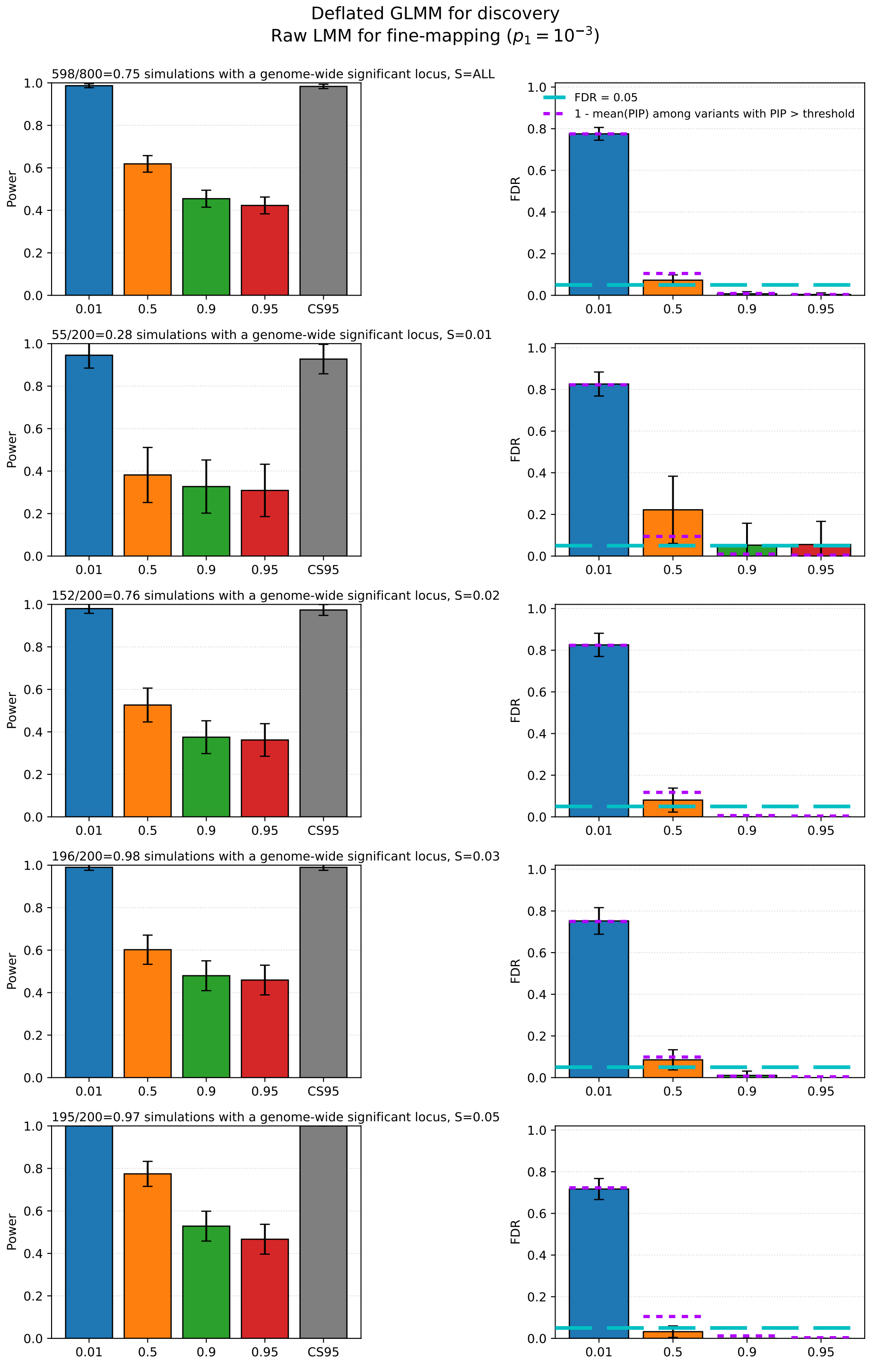

**Figure S6:** Calibration and power of fine-mapping of selection loci, using deflated GLMM selection statistics for discovery (i.e., to decide whether to run fine-mapping at a given locus) and raw LMM selection statistics as input for the fine-mapping pipeline. To account for the fact that LMM selection statistics are less well-powered than GLMM selection statistics (Supplementary Text A.2), we set $p_1 = 10^{-3}$ (the prior of there being a non-zero effect for selection at the locus), higher than the default $p_1 = 10^{-4}$. We show power (including for belonging to the 95% credible set, CS95) and FDR, stratifying by value of $s$ (selection coefficient, different rows) and for variants with PIP above a given threshold $t$ ($X-$axis ticks); we plot the value across all simulations (plot estimate) and 1.96 $\times$ SE, computing standard errors (SEs) via a leave-one-simulation-out jackknife. For FDR, we indicate both 5% and 1-[average PIP for variants with PIP $> t$], which should approximate the true FDR under well-calibrated posterior probabilities. See Materials and Methods for full details.

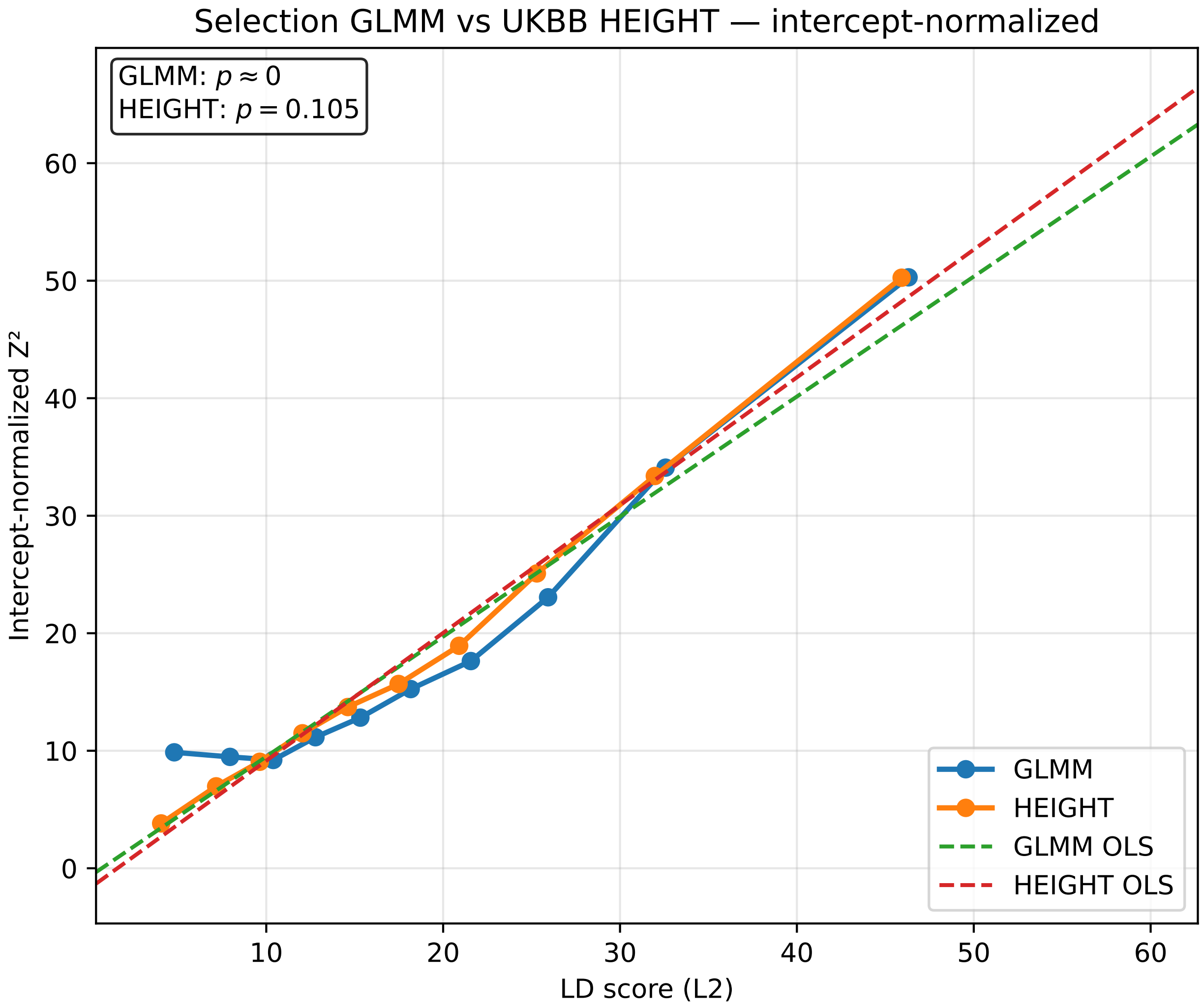

**Figure S7:** Scaling of average deflated GLMM selection or average GWAS $\chi^2$ ($Z^2$) with LD score (*103*), in 10 LD-score deciles. The curves have been normalized to be visualized overlaid on each other. The $p-$values are for a deviation from linearity, using a spline test with a genomic block jackknife across all (unbinned) HapMap3 ((*107*)) SNPs, excluding the HLA (*103*). As a representative GWAS trait, we show height, mapped in the UK Biobank (*104*, *105*) using boltLMM (*114*, *115*). See Materials and Methods for full details.

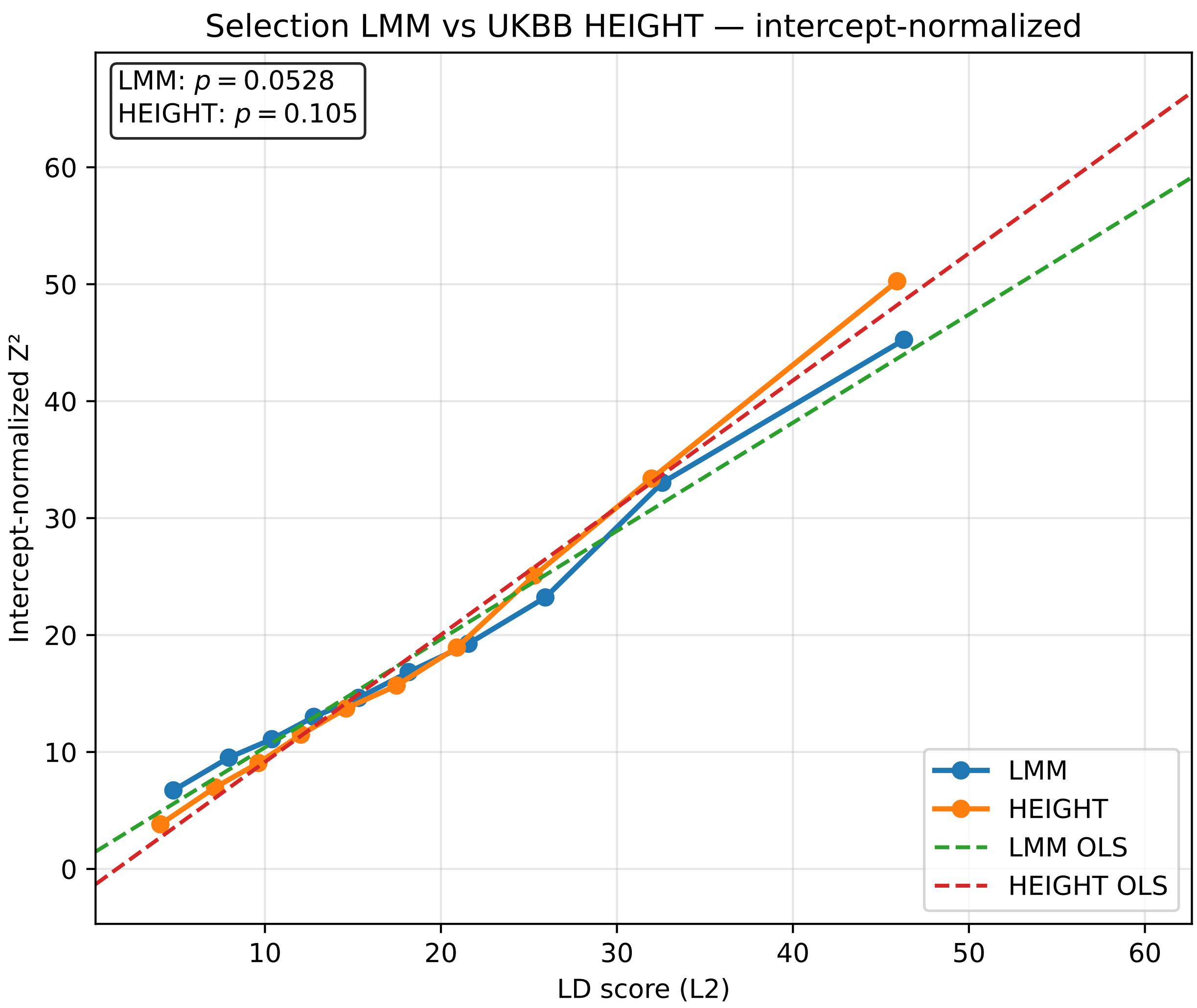

**Figure S8:** Scaling of average raw LMM selection or average GWAS $\chi^2$ ($Z^2$) with LD score ((*103*)), in 10 LD-score deciles. The curves have been normalized to be visualized overlaid on each other. The $p-$values are for a deviation from linearity, using a spline test with a genomic block jackknife across all (unbinned) HapMap3 ((*107*)) SNPs, excluding the HLA (*103*). As a representative GWAS trait, we show height, mapped in the UK Biobank (*104*, *105*) using boltLMM (*114*, *115*). See Materials and Methods for full details.

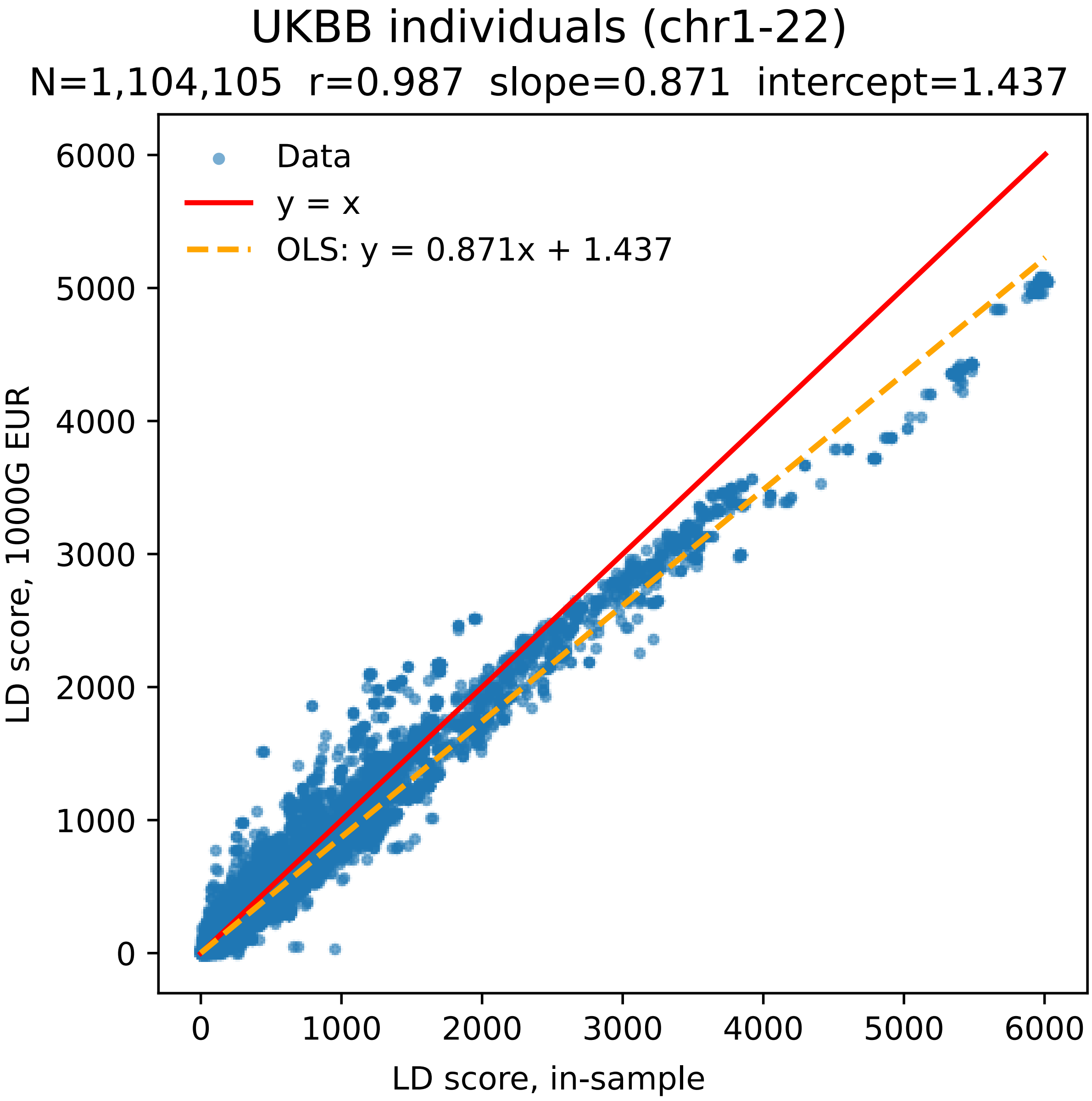

**Figure S9:** We show LD scores for HapMap3 SNPs (*107*), computed either in European-ancestry 1000 genomes individuals (**Auton2015**, *99*) or in UK Biobank (*104*, *105*) (contemporary) individuals included in the selection scan performed by (*86*). We report correlation and ordinary-least-squares results. See Materials and Methods for full details.

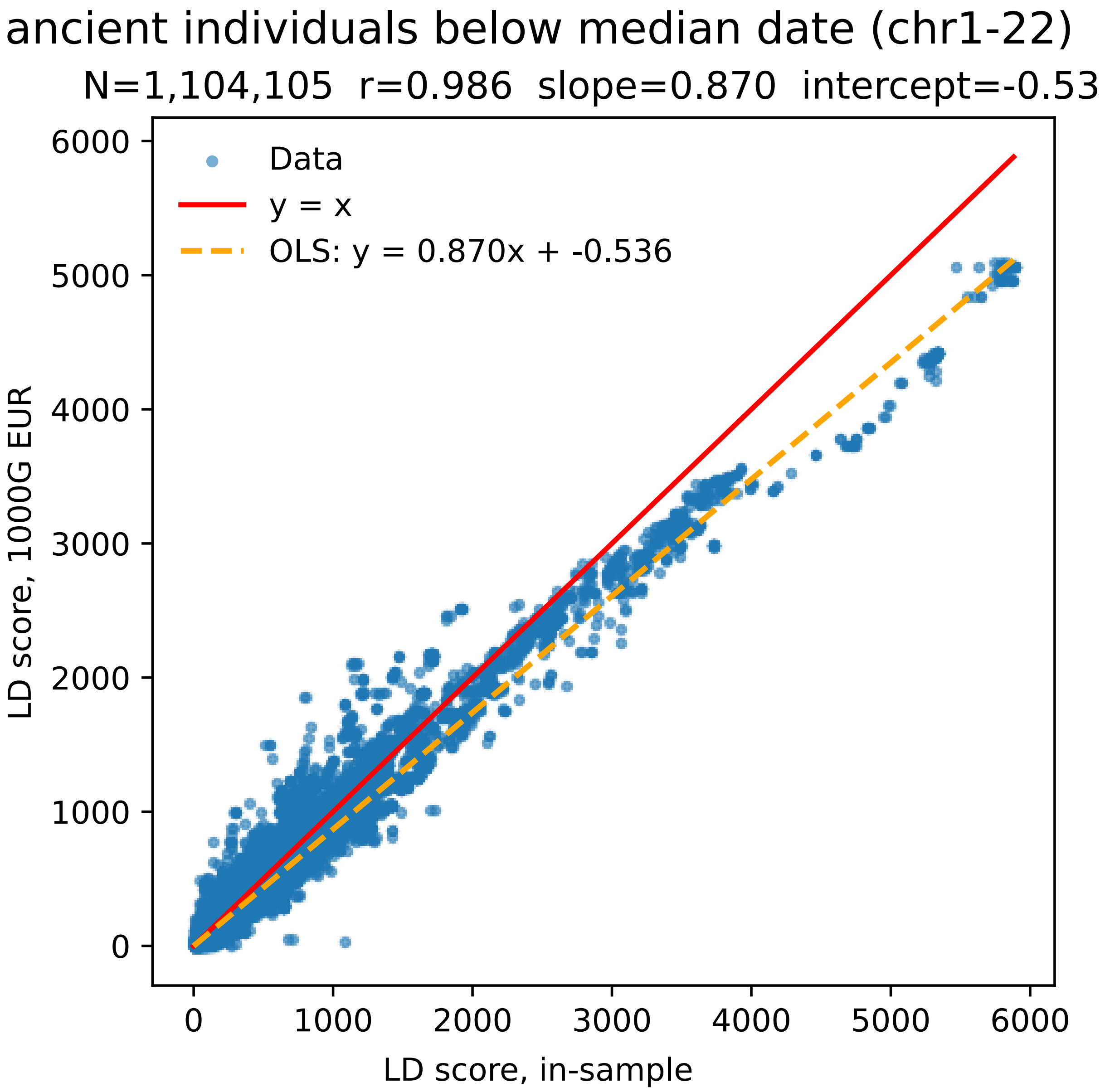

**Figure S10:** We show LD scores for HapMap3 SNPs (*107*), computed either in European-ancestry 1000 genomes individuals (**Auton2015**, *99*) or in ancient individuals included in the selection scan performed by (*86*) whose data is more recent than the median sample date for ancient individuals. We report correlation and ordinary-least-squares results. See Materials and Methods for full details.

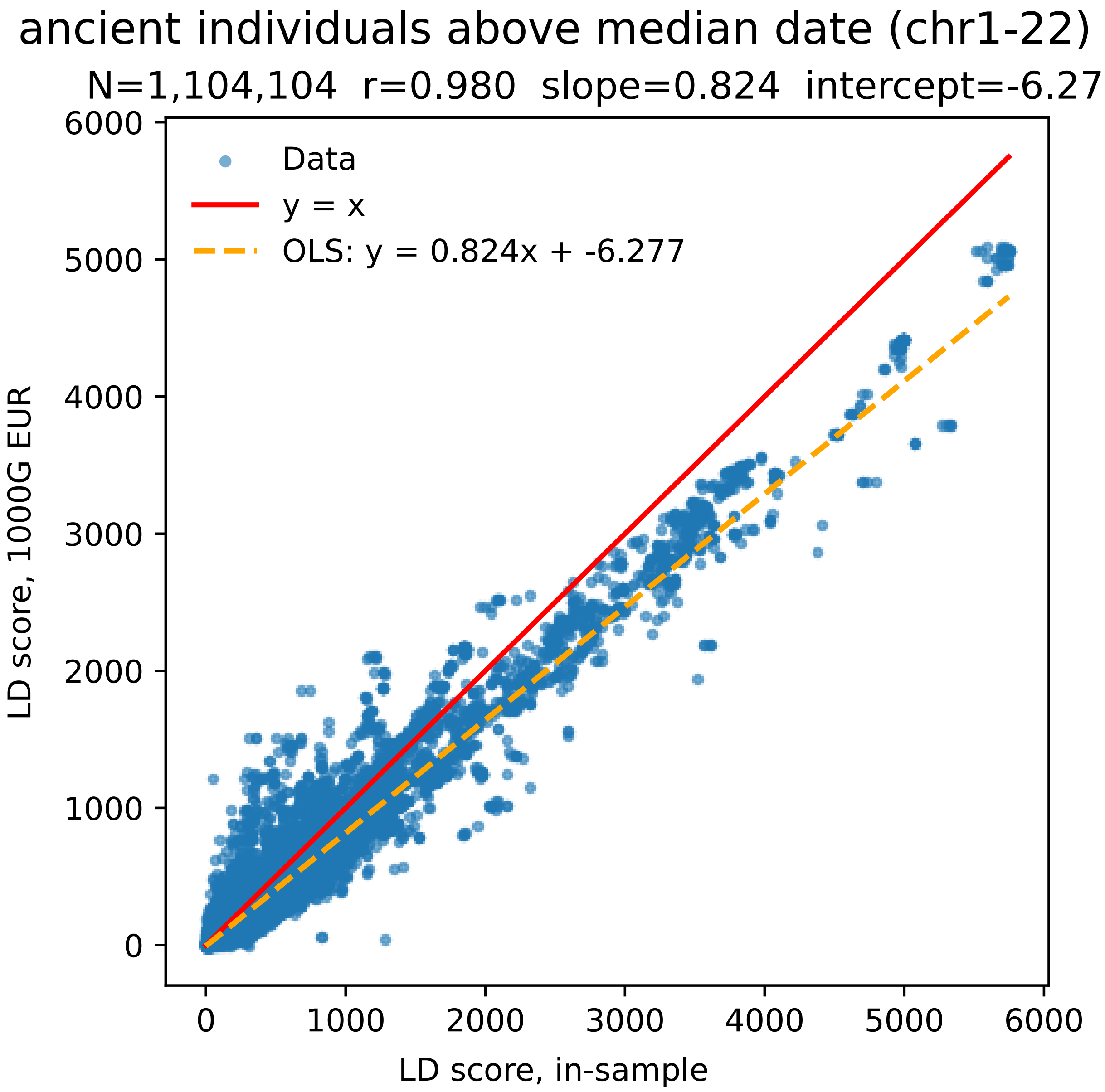

**Figure S11:** We show LD scores for HapMap3 SNPs (*107*), computed either in European-ancestry 1000 genomes individuals (**Auton2015**, *99*) or in ancient individuals included in the selection scan performed by (*86*) whose data is older than the median sample date for ancient individuals. We report correlation and ordinary-least-squares results. See Materials and Methods for full details.

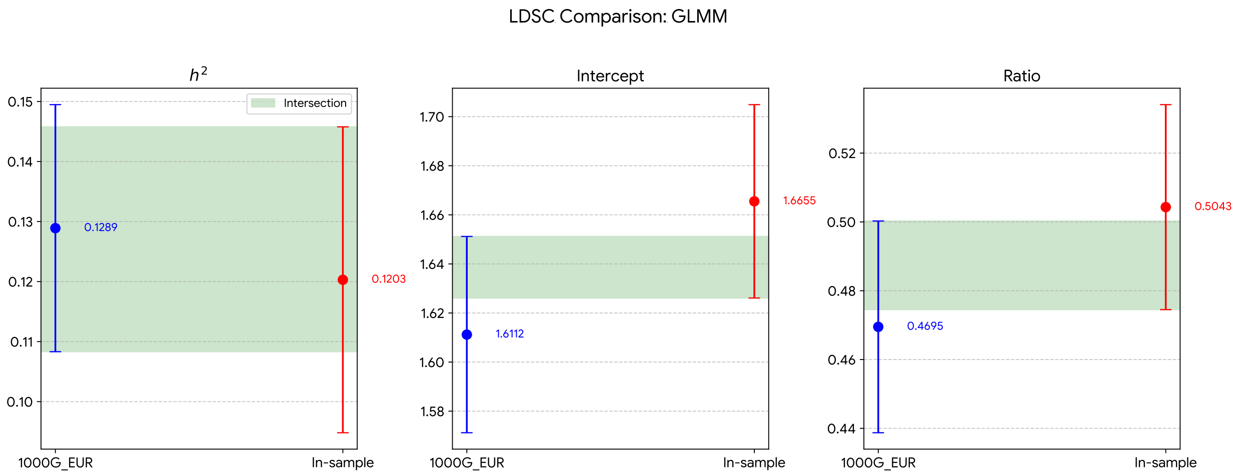

**Figure S12:** We show three key quantities estimated by LDSC for deflated GLMM selection statistics; when using LD scores (*103*) computed using European-ancestry individuals from the 1000 genomes project (**10002015global**, *99*), or when using LD scores computed in sample, using the exact same set of individuals that were included in the selection scan performed by (*86*). We show LDSC point estimates with 1.96 $\times$ SEs. We caution that the showed intersection does not take into account covariance in estimation errors. See Materials and Methods for full details.

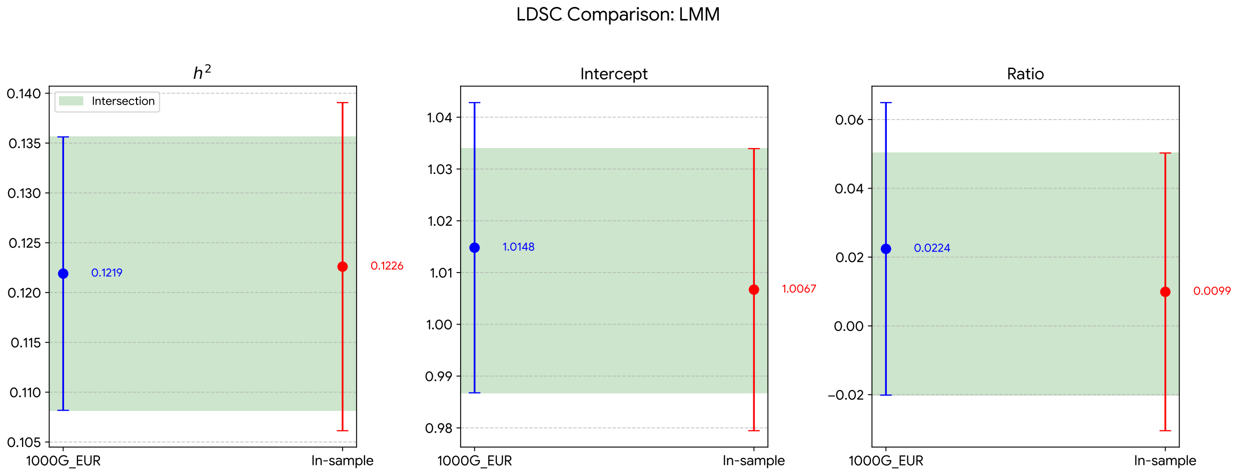

**Figure S13:** We show three key quantities estimated by LDSC for raw LMM selection statistics; when using LD scores (*103*) computed using European-ancestry individuals from the 1000 genomes project (**10002015global**, *99*), or when using LD scores computed in sample, using the exact same set of individuals that were included in the selection scan performed by (*86*). We show LDSC point estimates with 1.96 $\times$ SEs. We caution that the showed intersection does not take into account covariance in estimation errors. See Materials and Methods for full details.

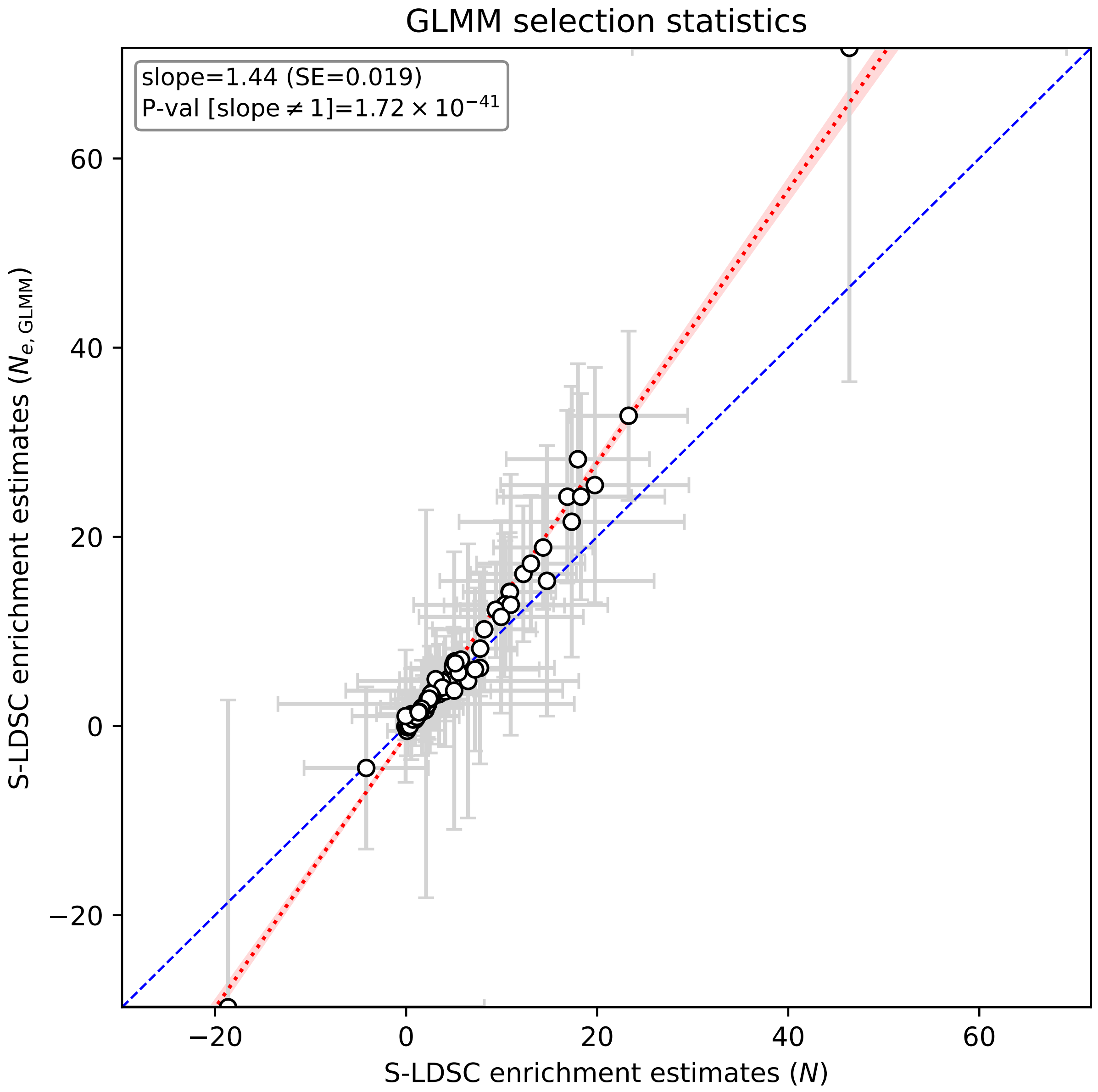

**Figure S14:** We show S-LDSC functional-enrichment estimates for deflated GLMM selection statistics, when using either $N$ or $N_{e,\mathrm{GLMM}}$ (Materials and Methods) as input sample-size value. We regressed the latter onto the former, and assessed the significance of a slope significantly different than 1 using ordinary least-squares. We caution that the $p-$ value is likely anticonservative due to assuming independence across functional annotations; for a test that takes into account cross-annotation covariance, see Figure S16. See Materials and Methods for full details.

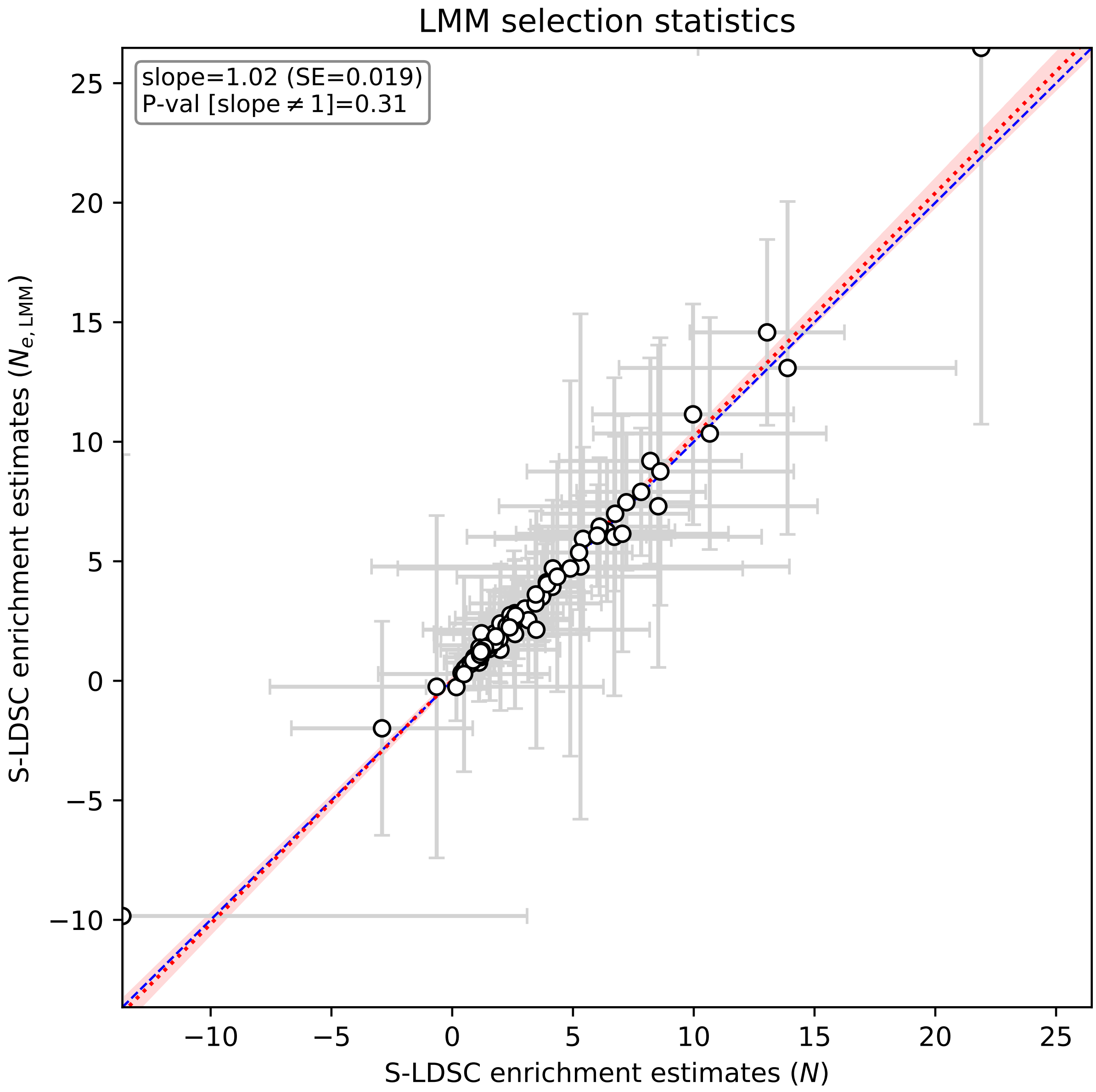

**Figure S15:** We show S-LDSC functional-enrichment estimates for raw LMM selection statistics, when using either $N$ or $N_{e,\text{LMM}}$ (Materials and Methods) as input sample-size value. We regressed the latter onto the former, and assessed the significance of a slope significantly different than 1 using ordinary least-squares. We caution that the $p-$ value is likely anticonservative due to assuming independence across functional annotations; for a test that takes into account cross-annotation covariance, see Figure S17. See Materials and Methods for full details.

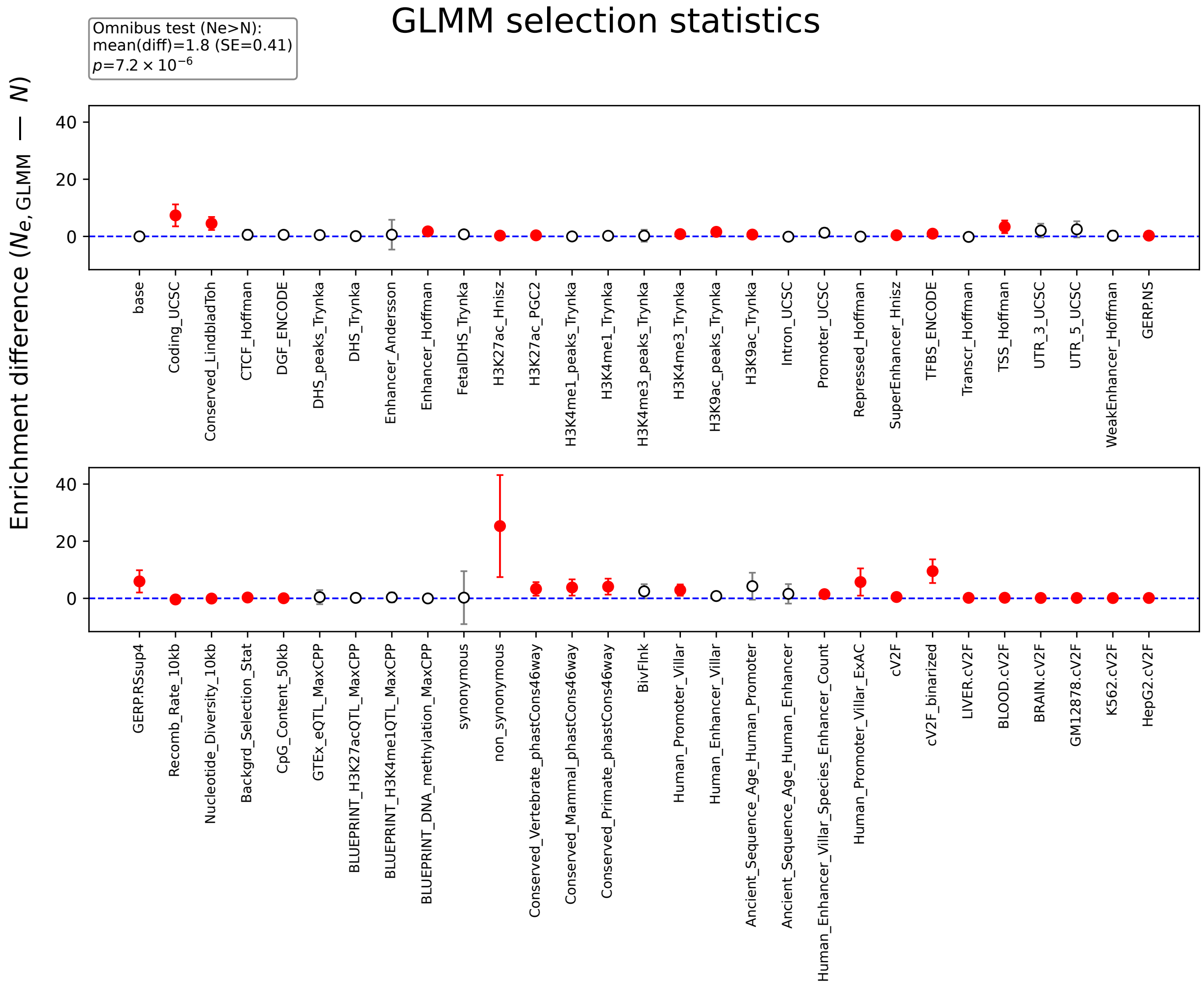

**Figure S16:** For selected functional annotations and deflated GLMM selection statistics, we show the difference between enrichment estimate for $N$ and enrichment estimate for $N_{e,\mathrm{GLMM}}$ as assessed by jackknifing this difference. Bars denote 1.96 $\times$ SEs. Differences nominally significantly larger than 0 are colored in red. The omnibus tests assessed the significance of a non-zero average difference across annotations, jackknifed across annotations to take into account covariance among them. See Materials and Methods for full details.

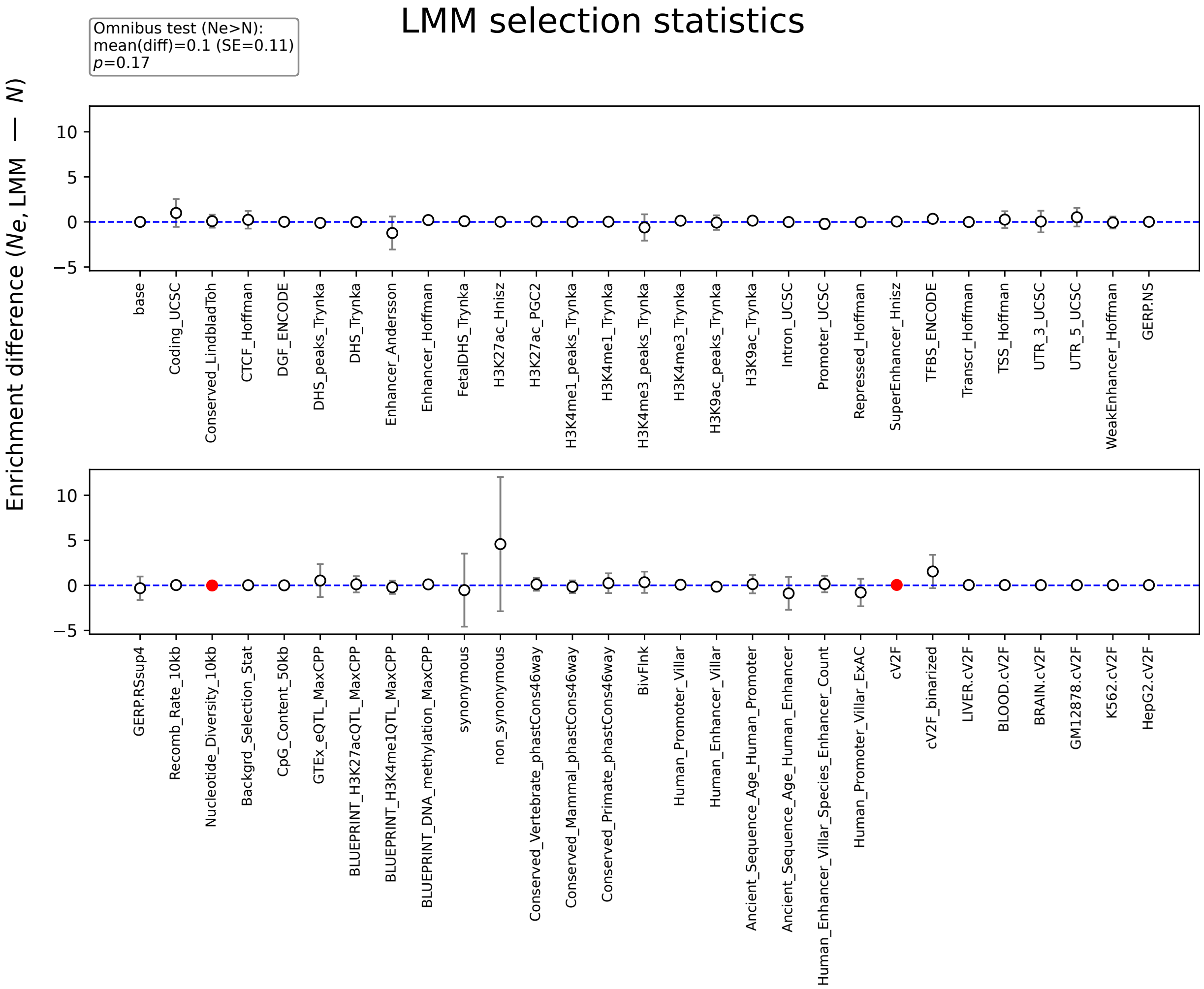

**Figure S17:** For selected functional annotations and raw LMM selection statistics, we show the difference between enrichment estimate for $N$ and enrichment estimate for $N_{e,\mathrm{LMM}}$ as assessed by jackknifing this difference. Bars denote $1.96 \times$ SEs. Differences nominally significantly larger than 0 are colored in red. The omnibus tests assessed the significance of a non-zero average difference across annotations, jackknifed across annotations to take into account covariance among them. See Materials and Methods for full details.

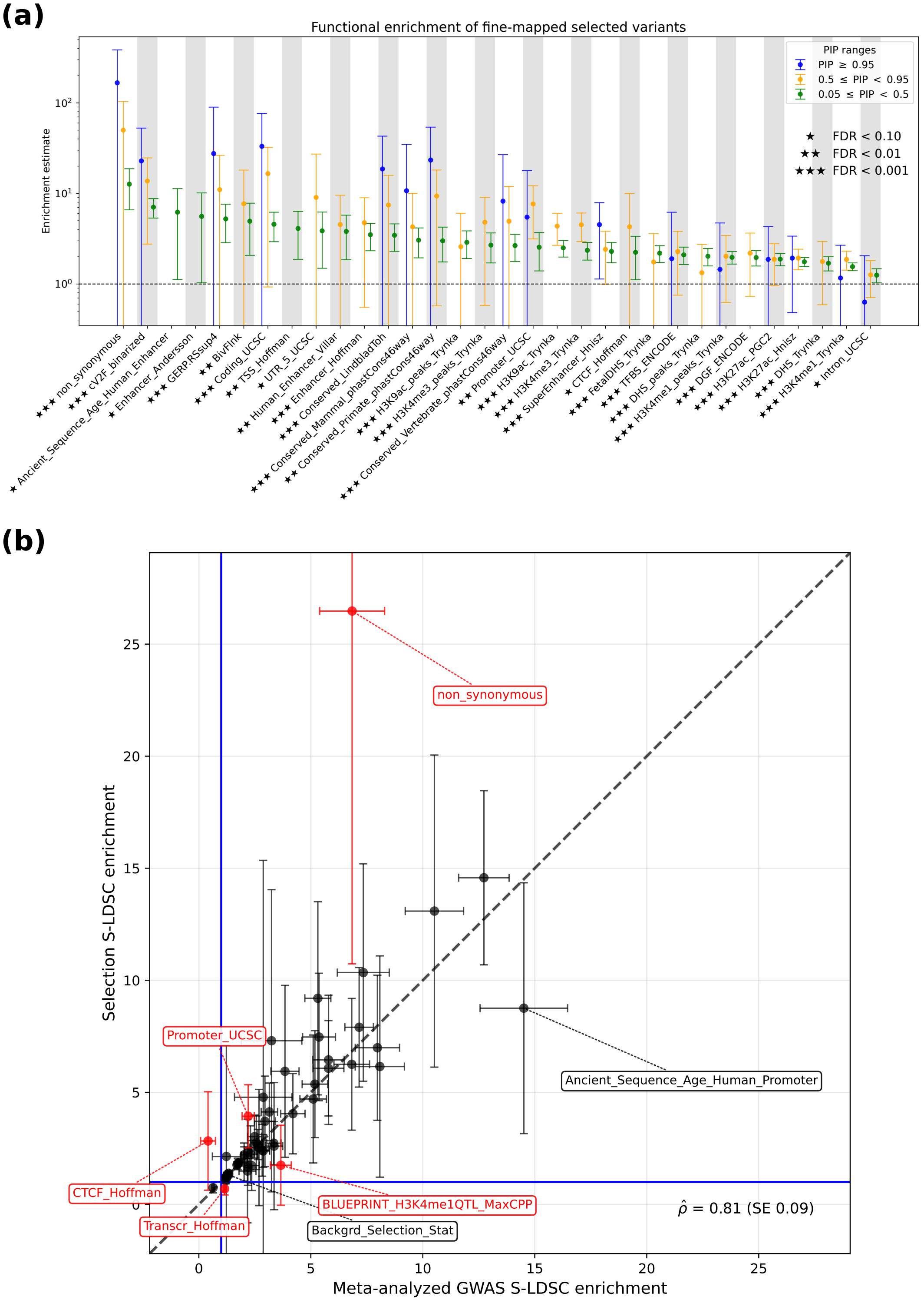

**Figure S18: Functional architecture of adaptation**. **(a)** Functional enrichment of fine-mapped selected variants. We define enrichment for a given binary annotation at a given posterior inclusion probability (PIP) threshold as the ratio of the proportion of variants that have PIP larger than the threshold, relative to the overall proportion of variants belonging to that annotation. We show 1.96 standard errors, estimated via a genomic-block jackknife with 200 blocks. **(b)** Comparison of S-LDSC functional enrichments for meta-analyzed GWAS traits and adaptation. Functional enrichment estimated by S-LDSC is similarly defined as the proportion of causal variance of selection signals contained in a given annotation, relative to the overall prevalence of that annotation. We show 1.96 standard errors, estimated via a genomic-block jackknife with 200 blocks. Full numerical results are reported in Data S1.

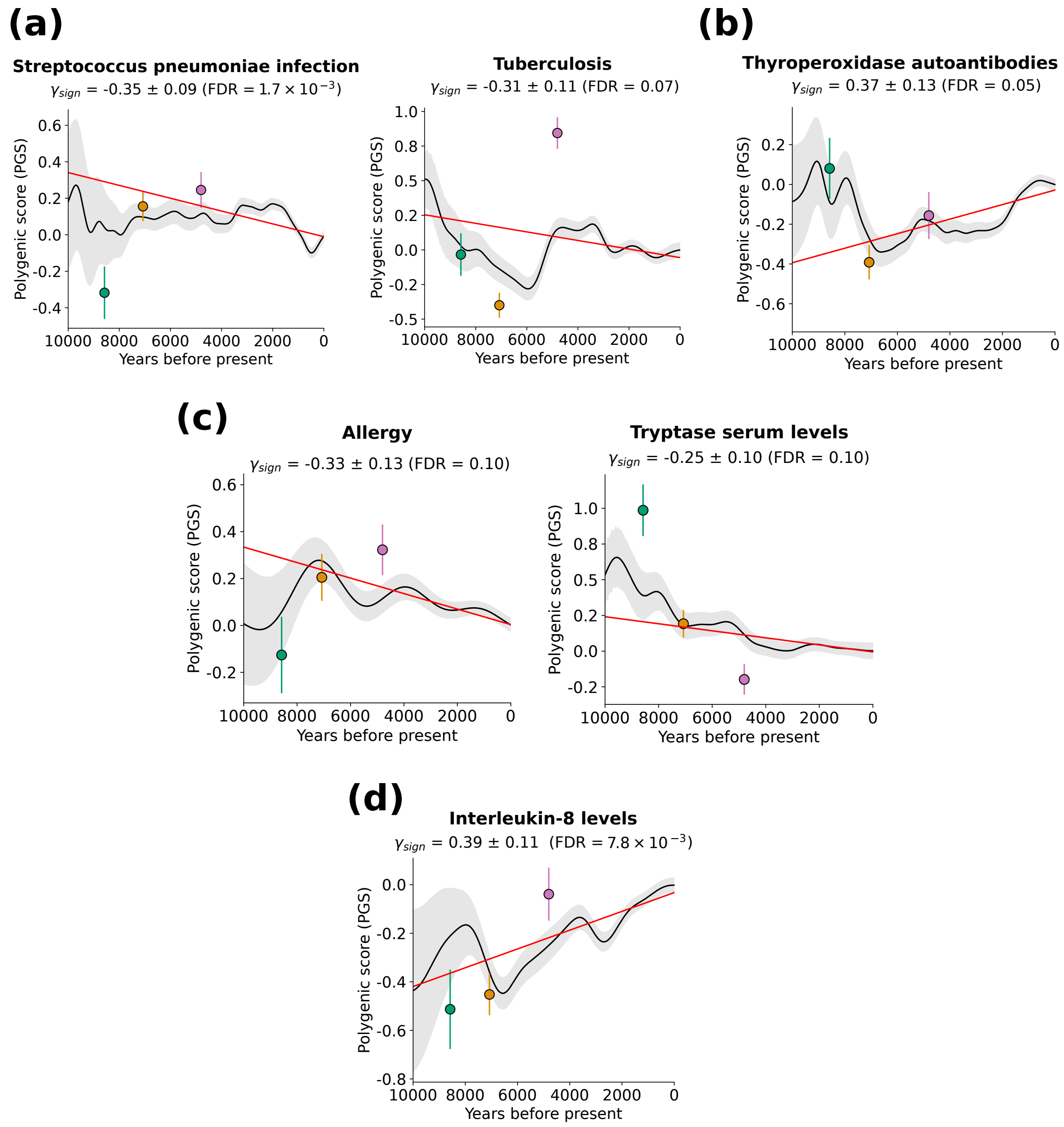

**Figure S19: Selected additional polygenic-test results.** **(a)** Additional significant decreases in genetically-predicted infection risk. *Streptococcus pneumoniae* is a major cause of pneumonia in humans(*184*). **(b)** Increase in genetically-predicted thyroperoxidase autoantibodies. Thyroperoxidase autoantibodies, which are directed against the body's own thyroid peroxidase enzyme, are a common biomarker of autoimmune thyroid disease(*184*). **(c)** Decreases in genetically-predicted risk for additional allergy-related phenotypes. Tryptase is a biomarker of allergic reactions (an enzyme mainly released by mast cells, immune cells heavily involved in such reactions)(*184*). **(d)** Increase in genetically-predicted interleukin-8 levels. Interleukin-8 is a major mediator of inflammatory response*(41)*. In **(a)**-**(d)**, the solid black line shows, for selected GWAS traits, the mean polygenic score (PGS) (an individual's effect size-weighted dosage of risk-increasing alleles that is used as a genetic predictor of a trait, trained in modern European individuals) of western Eurasian individuals over the past 10,000 years, with the 95% confidence band in grey. The red line indicates a polygenic test that assesses whether this average genetic trait predictor has shifted in magnitude beyond the neutral expectation (via a linear mixed model regression adjusted for population structure, with slope $\gamma_{\text{sign}}$, see Materials and Methods for details). Circles denote the mean PGS for Western hunter–gatherers (green; n = 131), early European farmers (orange; n = 452), and steppe pastoralists (pink; n = 293), proxies for the three ancestral populations to modern Europeans*(65)*. Error bars represent 95% confidence intervals. Nominal statistical significance for a difference from 0 was assessed via a two-tailed Z test; FDR refers to Benjamini-Hochberg(66) (BH) q-values. Full numerical results are reported in Data S1.

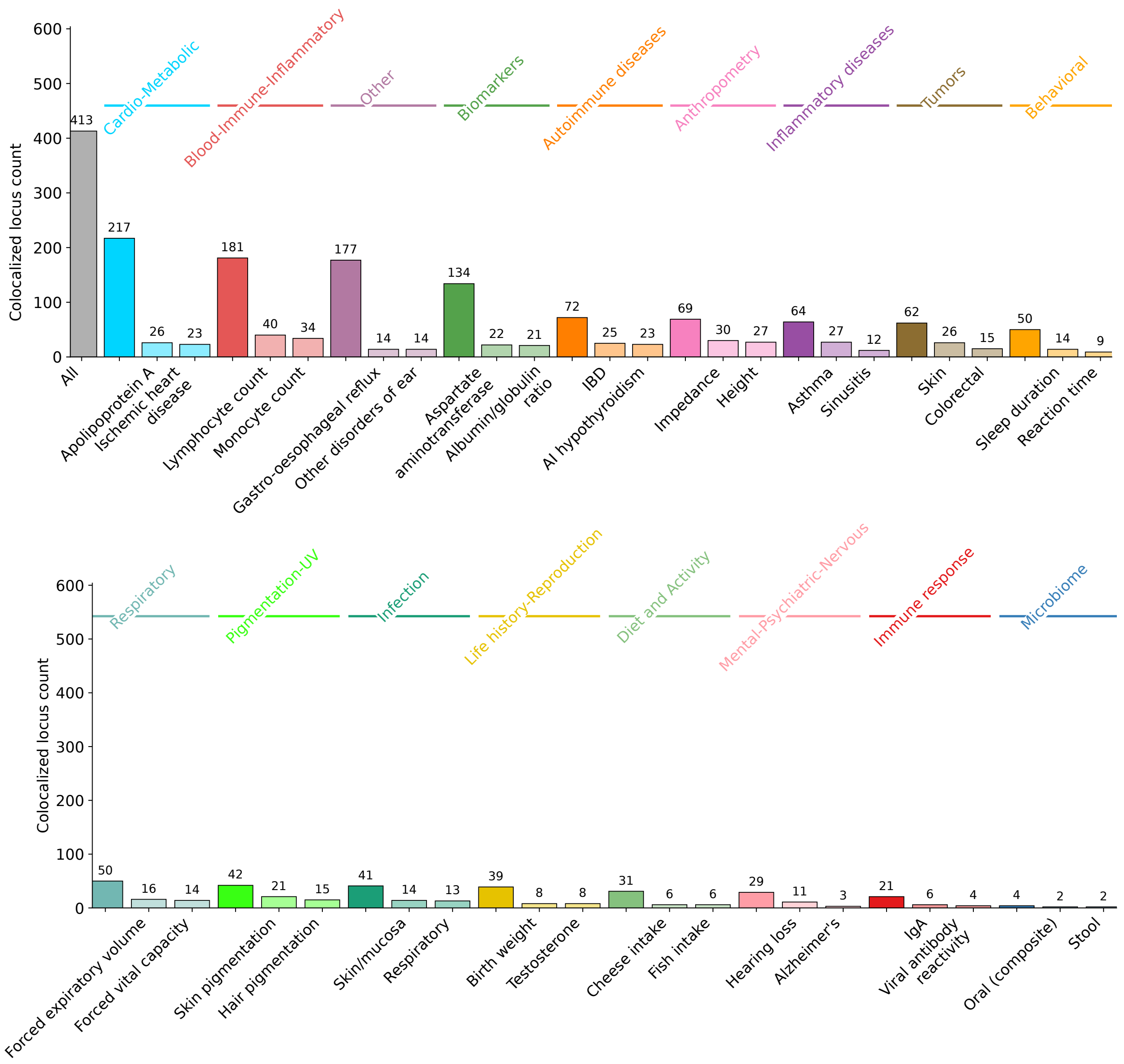

**Figure S20: Colocalizations between selection and GWAS traits partitioned into 17 trait categories**. The two traits with the highest number of colocalizations are shown for each category. See Figure S21 for finer-grained results for immune-related categories. Full numerical results for all trait-selection comparisons are reported in Data S3.

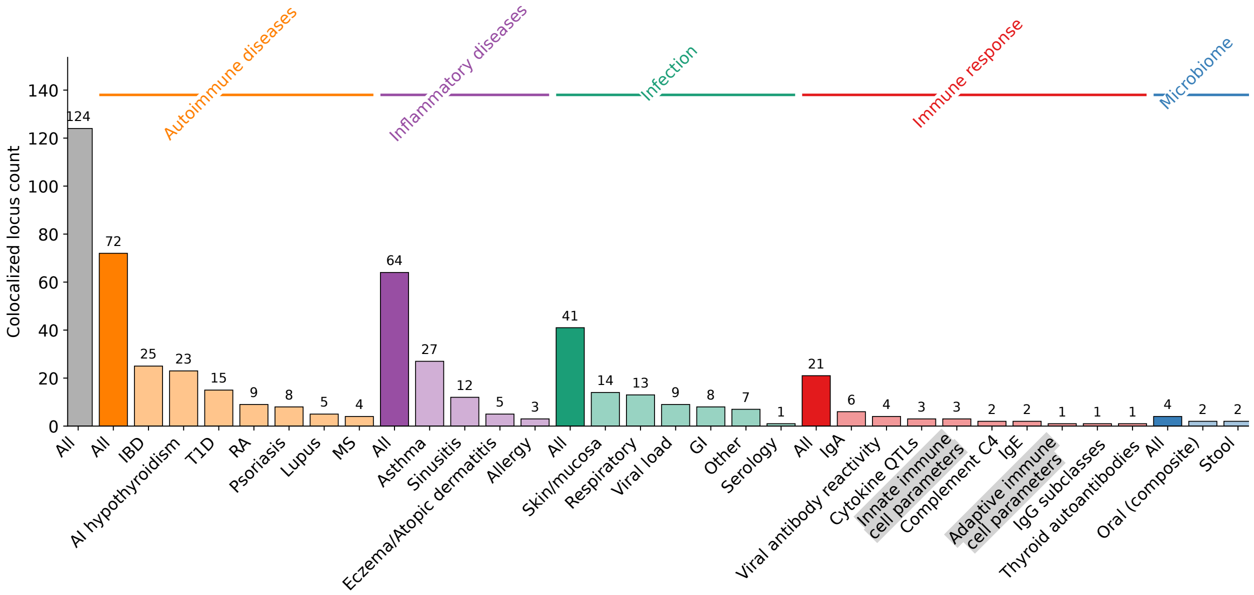

**Figure S21:** We show colocalizations between selection and fine-grained immune-related GWAS traits, as assessed by `coloc` (*94*). See Materials and Methods for full details; see also Figure S20. Full numerical results for all trait-selection comparisons are reported in Data S3.

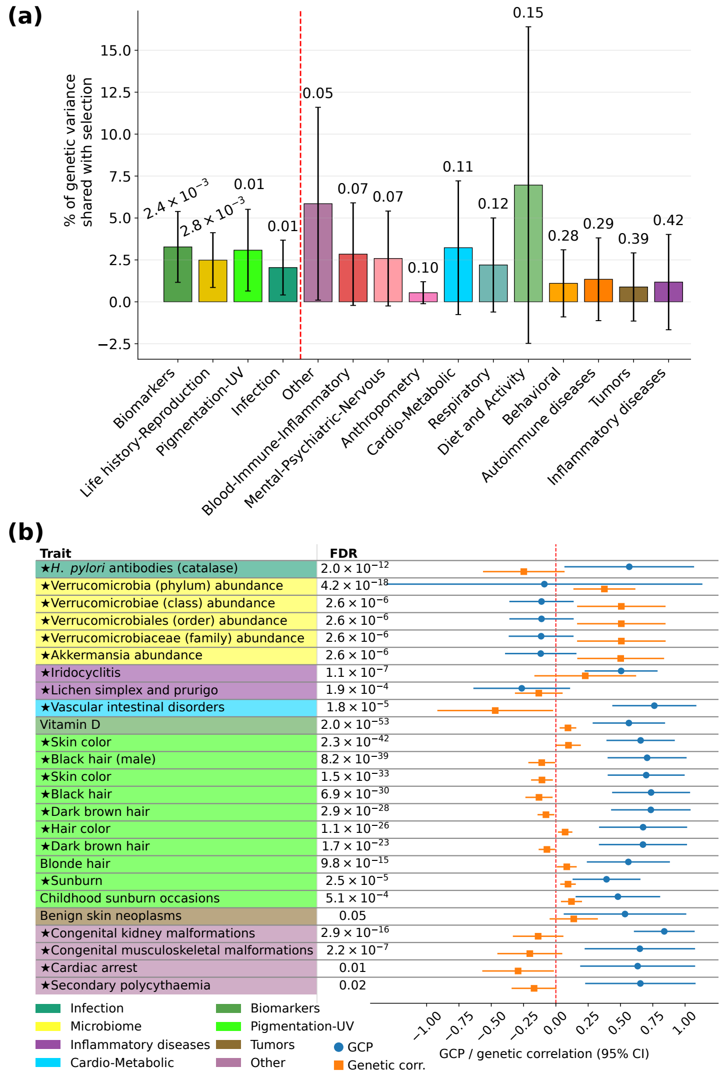

**Figure S22: Secondary analyses relating selection to GWAS traits**. Full numerical results are reported in Data S1. **(a)** PHBC results assessing trait genetic variance shared with adaptation. Pleiotropic shared Heritability with Bias Correction (PHBC) estimates the fraction of trait genetic variance shared with adaptation across a given set of auxiliary traits, using pairwise genetic correlations estimated by cross-trait LDSC (generalizing the squared genetic correlation with a single trait)(*180*) while applying a bias correction to correct for upward bias arising from finite sample size. We only used auxiliary traits with a heritability $Z$-score above 6. To avoid an unstable estimate in the presence of collinearity among auxiliary traits, we performed an initial selection on auxiliary traits for each trait category as follows: for each pair of auxiliary traits with $r_g^2 > 0.5$, we remove the trait with lower $h^2$ Z-score. For each of the 15 trait categories (spanning 268 well-powered traits), we obtained a category-specific estimate using post-pruned auxiliary diseases belonging to that category. We show 1.96 standard errors, estimated via a genomic-block jackknife with 200 blocks. Nominal $p$-values for a significantly non-zero shared genetic variance are shown above each category; those to the left of the red line were significant (FDR<10%), including pigmentation and infection. **(b)** LCV results for causal relationships between adaptation and GWAS traits. We show FDR for a significant LCV(*188*) result, assessing evidence of a causal relationship between adaptation and GWAS traits. LCV also estimates the genetic causal proportion (GCP), the proportion of trait A that is causal for trait B. A positive GCP indicates the trait being causal for adaptation, a negative GCP indicates adaptation causally impacting the trait. We show the GCP with 1.96 standard errors, but we caution that this normality assumption is probably not appropriate in the context of the LCV model(*188*). To aid interpretation of directional relationships, we also show cross-trait LDSC genetic correlations between selection and the GWAS traits. Traits with a heritability Z-score below 4 were excluded from analysis due to LCV results being robust only under a scenario of polygenicity(*188*). Traits flagged with a star have a heritability $Z$-score greater than 4 but smaller than 7, potentially leading to false positives(*188*).

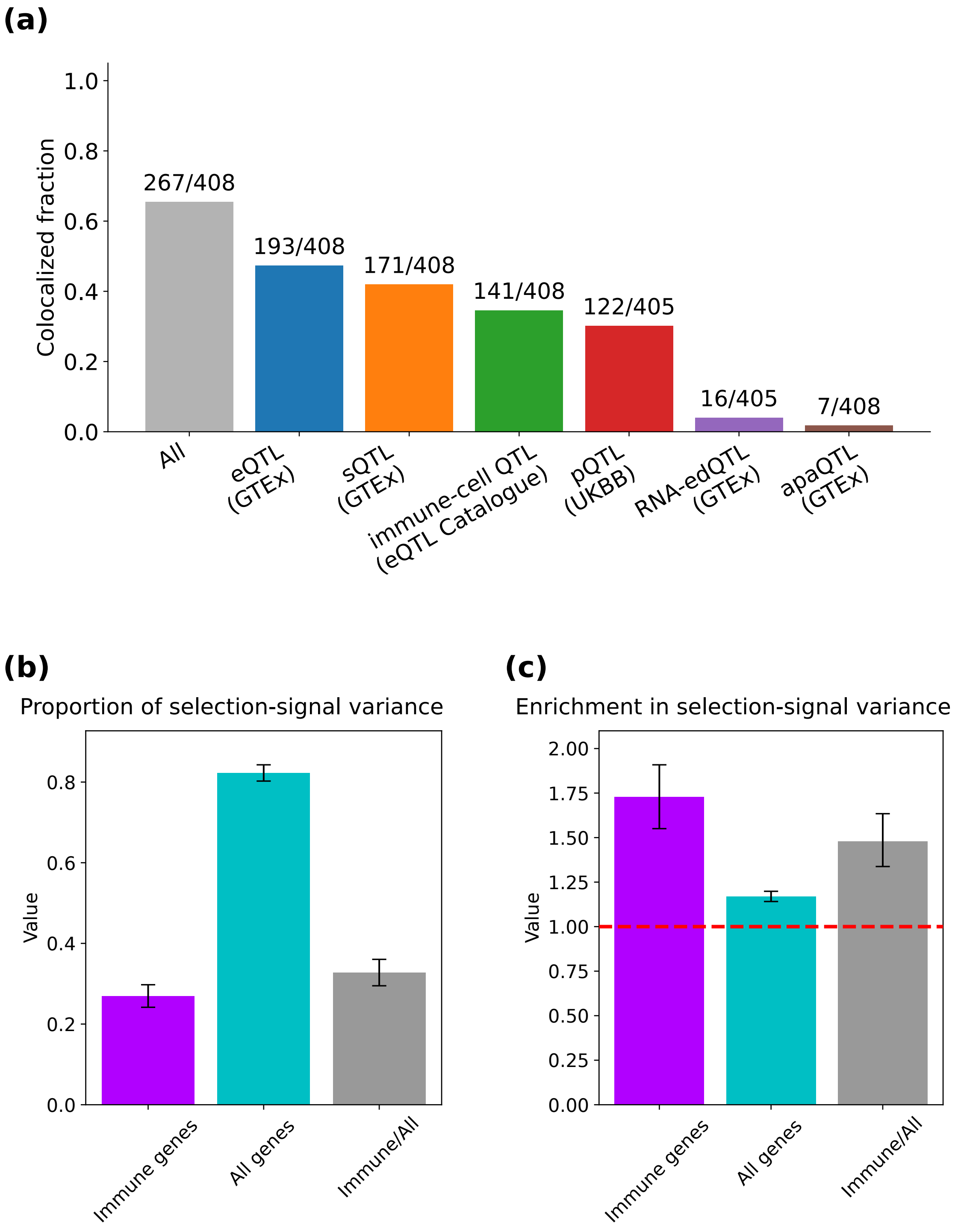

**Figure S23: Secondary analyses relating selection to genes**. **(a)** Colocalization rate between selection and different classes of QTLs. For the QTL classes we analyzed, we show the fraction of genome-wide significant loci for selection that colocalized with at least one QTL within that class. **(b)** Comparison between variance in signals of selection explained by variants near immune and all genes as estimated by S-LDSC. We annotated variants as falling within 100kb of an immune gene, following ref.(*158*) (Materials and Methods). We also included an annotation for variants falling within 100kb of any gene. We used S-LDSC to estimate the fraction of variance in selection signals explained by each annotation, jackknifing the ratio between the two. We show 1.96 standard errors, estimated via a genomic-block jackknife with 200 blocks. Full numerical results are reported in Data S3. **(c)** Comparison between enrichment in selection signals for variants near immune and all genes as estimated by S-LDSC. Similar to **(b)**, we used S-LDSC to estimate enrichment in selection-signal variance for each annotation, as well as jackknifing the ratio between enrichments for each of the annotations. Full numerical results are reported in Data S3.

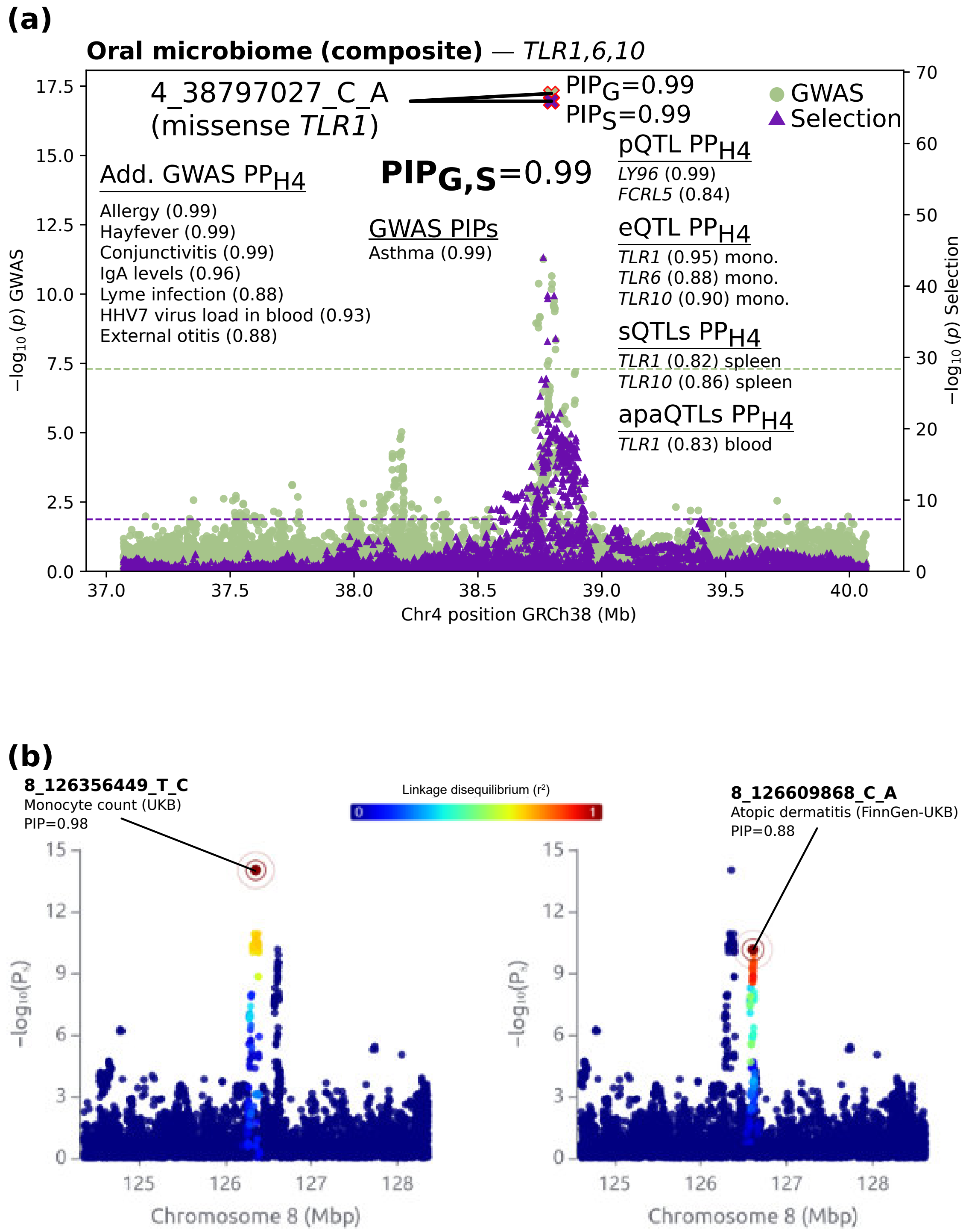

**Figure S24: Additional highlighted selection loci colocalizing with GWAS traits.** **(a)** A selection-oral microbiome colocalization at the well-known *TLR1*(*189*) selection locus. *TLR1* is a pattern-recognition receptor that detects bacterial lipoproteins and triggers inflammatory responses(*41*). This missense variant had $\text{PIP}_{\text{Sel}} > 0.99$, $\text{PIP}_{\text{GWAS}} > 0.99$ for oral microbiome composition, and joint $\text{PIP}_{\text{Sel,GWAS}} > 0.99$. (the locus also colocalized with 4 infection traits including conjunctivitis and Lyme disease). Additional colocalizations are indicated, together with the corresponding $\text{PP}_{\text{H4}}$ (the posterior probability of a shared causal variant, computed using `coloc`(*94*)). **(b)** GWAS fine-mapping identifies a plausible example of multiple independent causal variants at a selection locus. We highlight an instance of two seemingly independent selection signals within the same locus, as suggested by two lead variants in low LD to each other and being fine-mapped with high probability for two different GWAS traits. Full numerical results are reported in Data S3.

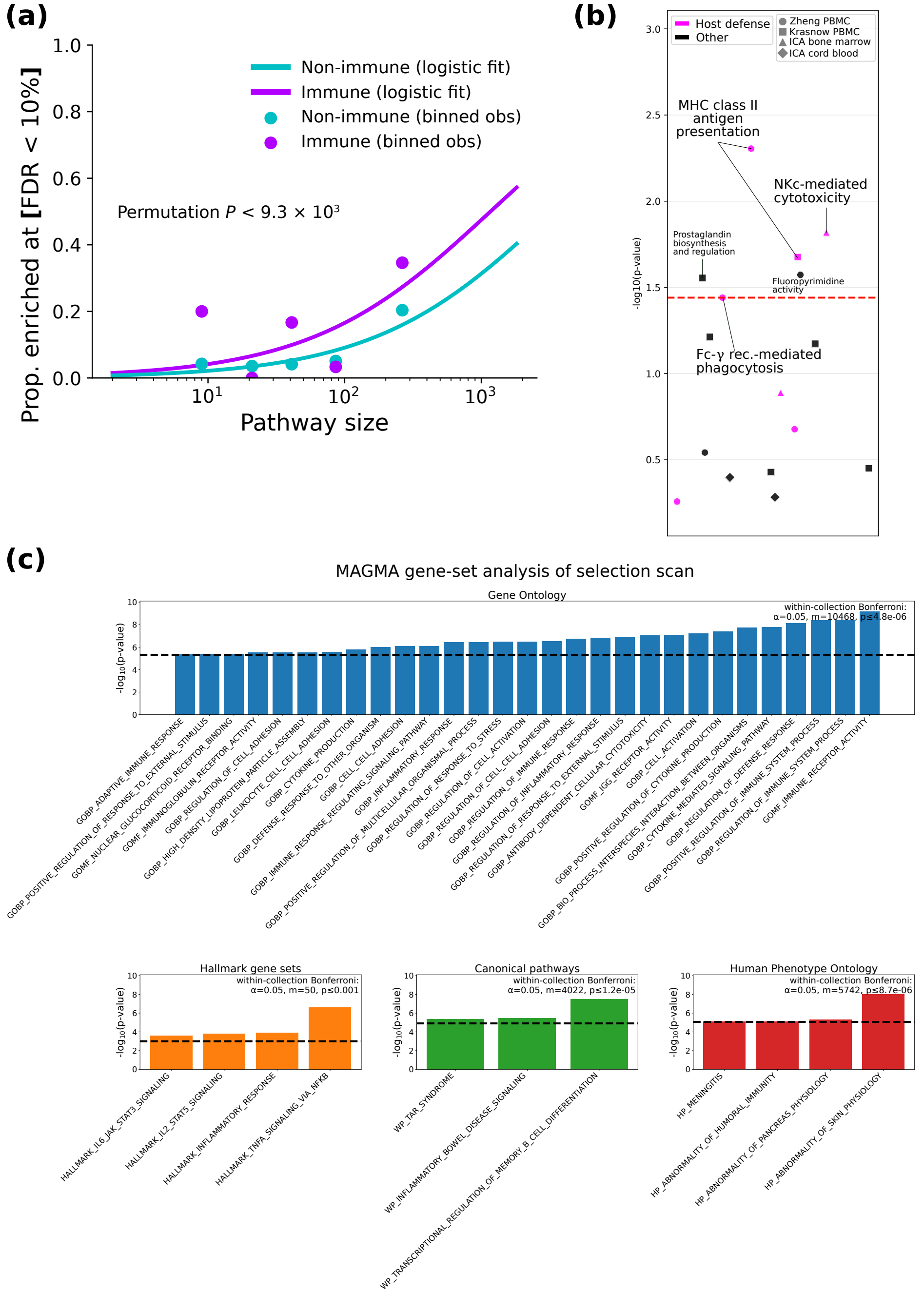

**Figure S25: Additional pathway analyses**. **(a)** Immune-related pathways are significantly more likely to be significantly enriched among genes with CALDERA>0.1 for adaptation. Analogue of Figure 4a for CALDERA>0.1 genes instead of GTEx-QTL colocalized genes. Curves represent a logistic regression fit adjusting for pathway size. *P*-values are computed via permutations to take into account redundancy in gene content across pathways (Materials and Methods). **(b)** Enrichment of signals of selection for blood-related biological processes as inferred by S-LDSC. We used S-LDSC to assess whether specific biological processes inferred from gene-expression data in blood-related tissues(*190*) are enriched in LMM selection statistic variance. The red line indicates FDR<10%. There was a pattern of host-defense processes being overrepresented among significant processes; however, this was not significant, likely due to the small sample size. **(c)** Enrichment of signals of selection for pathways across different gene-set collections as inferred by MAGMA. As an alternative to test for gene-set enrichment in selection signals, we used MAGMA, which aggregates selection evidence across variants within genes to obtain gene-level selection statistics; it then uses those statistics to look for pathways with a significant excess of signals of selection after adjusting for covariates like gene size or frequencies of within-gene variants(*150*). Similar to alternative methods, MAGMA highlighted pathways related to immunity and host defense. Full numerical results are reported in Data S4.

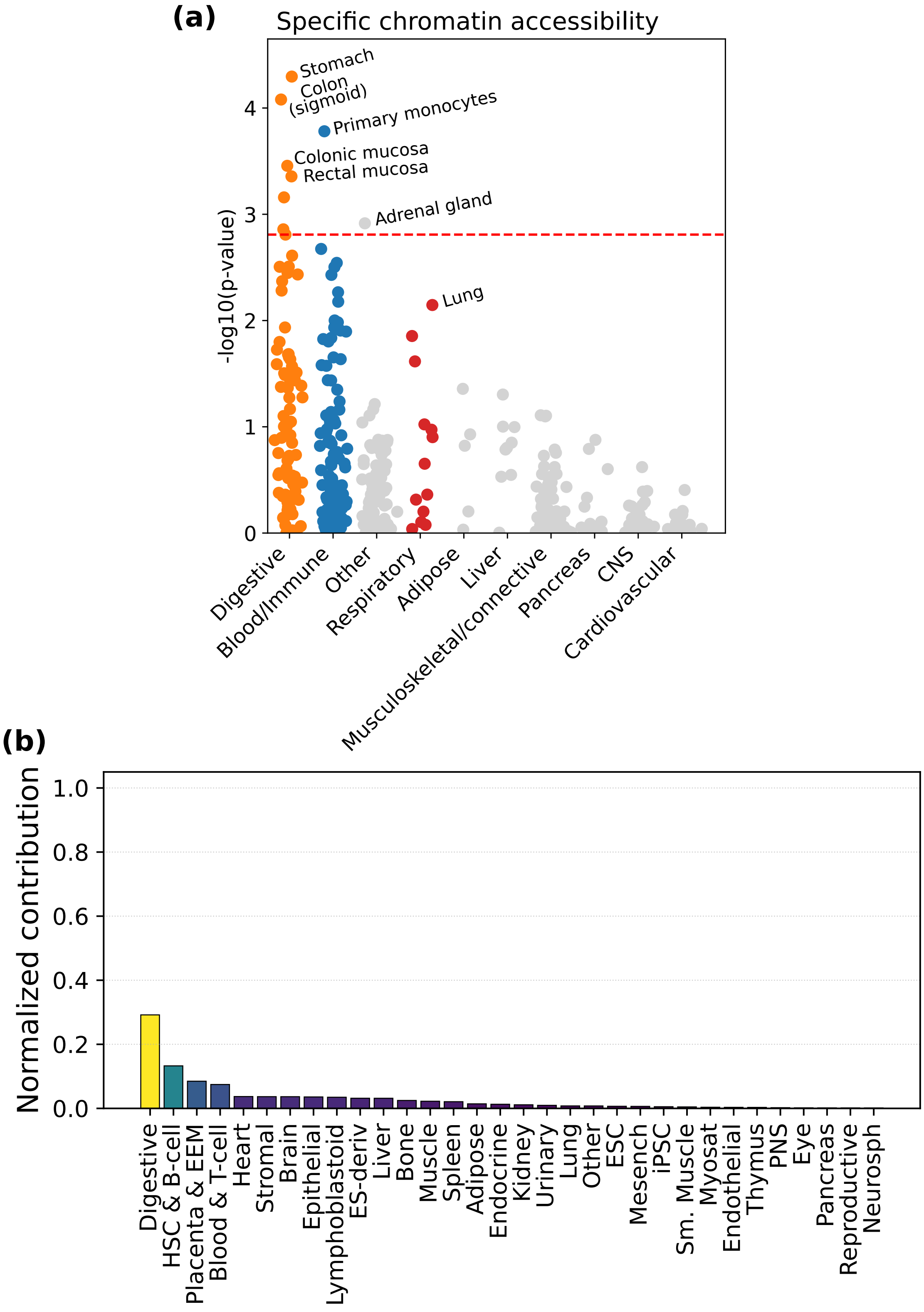

**Figure S26: Additional tissue analyses.** **(a)** Tissues with enrichment for signals of selection as inferred from tissue-specific chromatin accessibility. Analogue of Figure 5b using tissue-specific chromatin accessibility instead of tissue-specific gene expression. **(b)** Epigenomic partitioning of fine-mapped selected variants. We used J-PEP(*191*) to perform epigenomic partitioning of fine-mapped selected variants. Briefly, J-PEP uses tissue-specific chromatin data to partition fine-mapped variants into clusters of variants specifically associated with a given tissue, inferring variant-to-cluster and cluster-to-tissue weights. In the case of selection, a single cluster was identified by this method, and we color tissues by relative contribution to this cluster. Full numerical results are reported in Data S5.

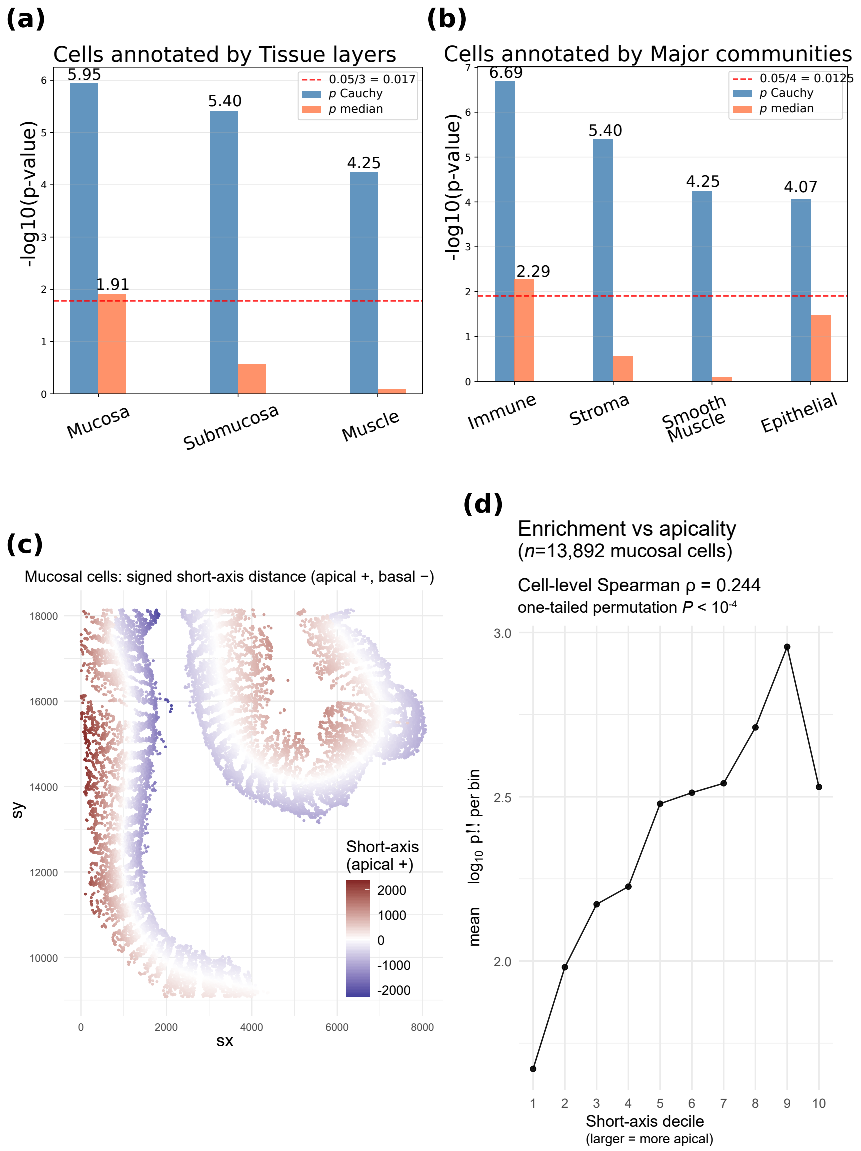

**Figure S27: Formal assessment of category-level enrichment using `gsMap` and relationship between cell apicality and `gsMap` cell-level enrichment in selection signals**. We show category-level enrichment $p$-values computed by `gsMap`, when stratifying cells by **(a)** tissue layer or **(b)** major community. These results are computed by aggregating across all 8 intestinal regions assayed by ref.(*173*), but results were consistent for individual intestinal regions and when stratifying cells by finer spatial neighborhoods or finer cell type (Supplementary Text C). Because `gsMap` computes category-level $p$-values via a Cauchy aggregation test(*171*, *172*) that is robust to $p$-value dependence at the price of being sensitive to individual outliers within an annotation, it is recommended to only view as significant those annotations that show both small Cauchy and small cross-cell median $p$-values(*171*). Separately, because sample B012 from ref.(*173*) had a visual trend of more apical mucosal cells (closer to the inside space of the gut) having more significant individual enrichment $p$-values (Figure 5c), we **(c)** quantified per-cell apicality by fitting a principal curve to the 2D coordinates and assigning mucosal cells to the nearest curve point, and defining apicality for each cell as its perpendicular distance to the assigned curve, with the sign oriented by a basal normal field: at each curve vertex, vectors to the $k$ nearest Submucosa/Muscle reference cells are averaged, smoothed along the curve, and normalized to define a consistent basal normal; cells projecting in the direction of this normal are labeled as basal (negative) and cells projecting opposite are apical (positive). We show **(d)** the association between per-cell apicality and per-cell enrichment $p$-value (restricted to mucosal cells) by computing the Spearman correlation between our apicality values and $-\log_{10}(p\text{-value})$ (from `gsMap`(*171*)). The significance of this association is assessed via a permutation test with $10^4$ permutations, to account for the possibility of non-independence across cells. We report the results computed individually across all cells (Figure SS37) but, for visualization purposes, show the trend in apicality deciles. Full numerical results are reported in Data S5.

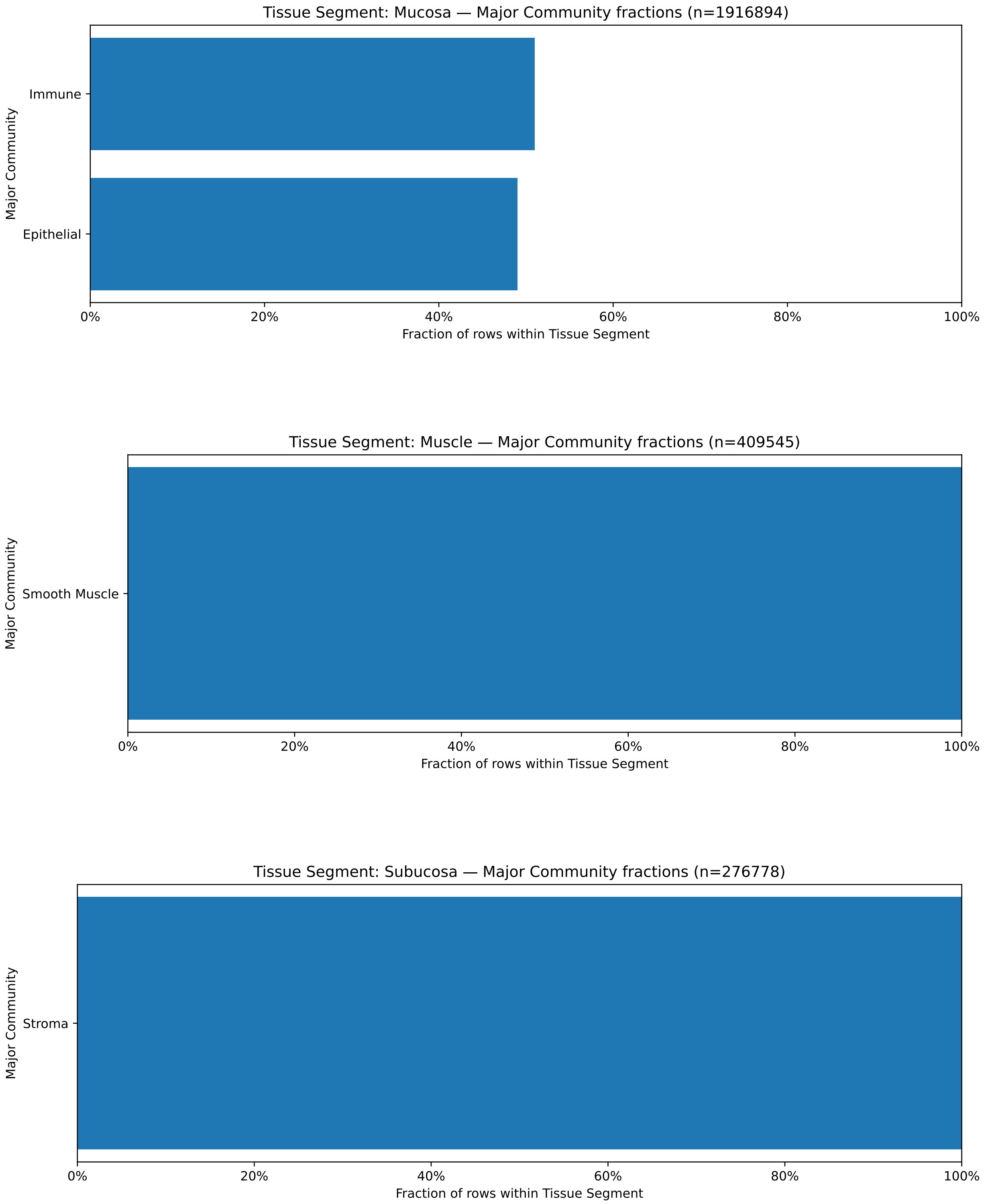

**Figure S28:** We show major-community fractions for cells from (*173*), stratified by tissue layers.

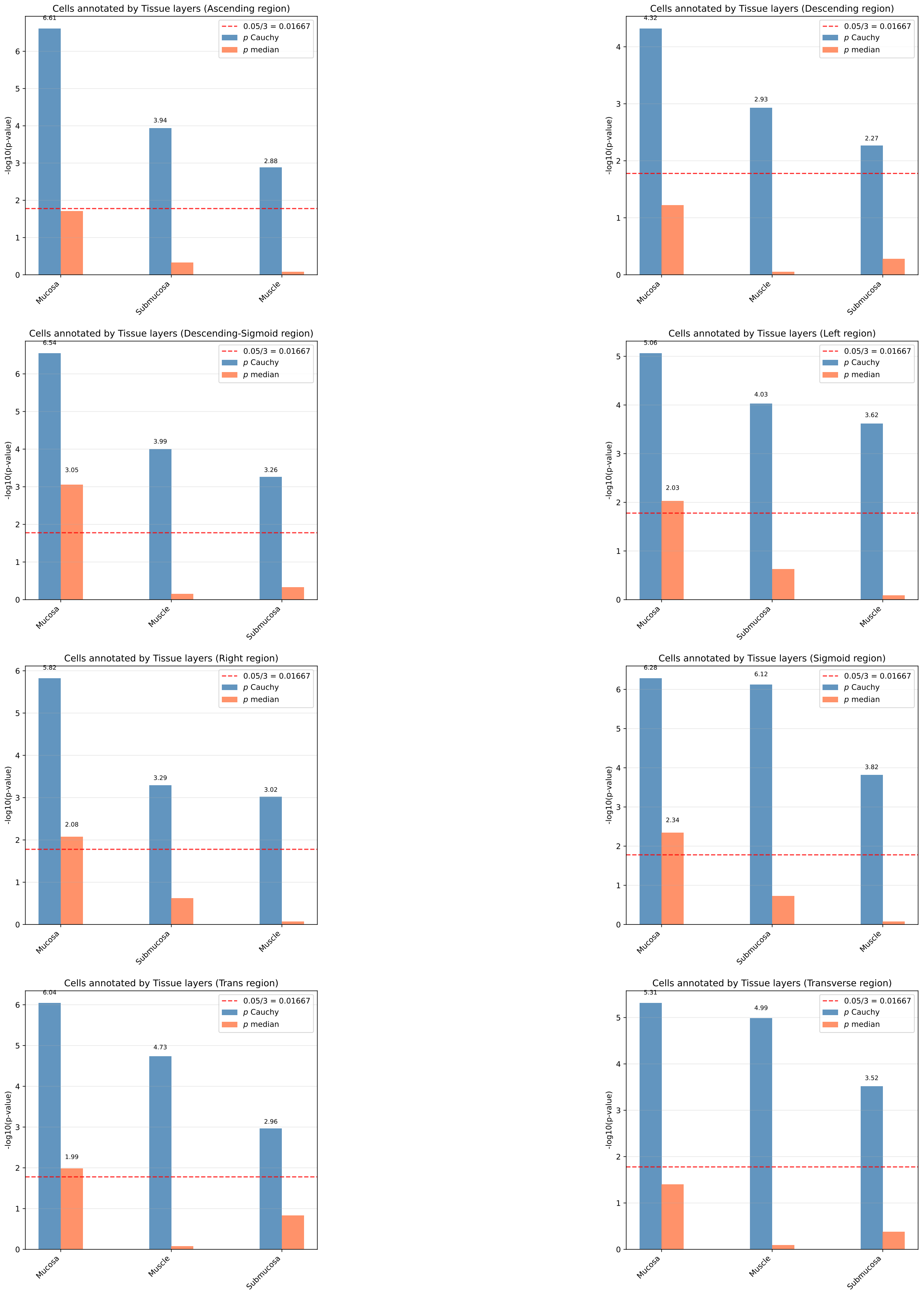

**Figure S29:** We show results of `gsMap` (*171*) for enrichments of selection signals in an intestinal single-cell, spatial transcriptomics dataset (*173*) across 8 intestinal regions of the human gut, when annotating cells by tissue layer. See Supplementary Text C for details.

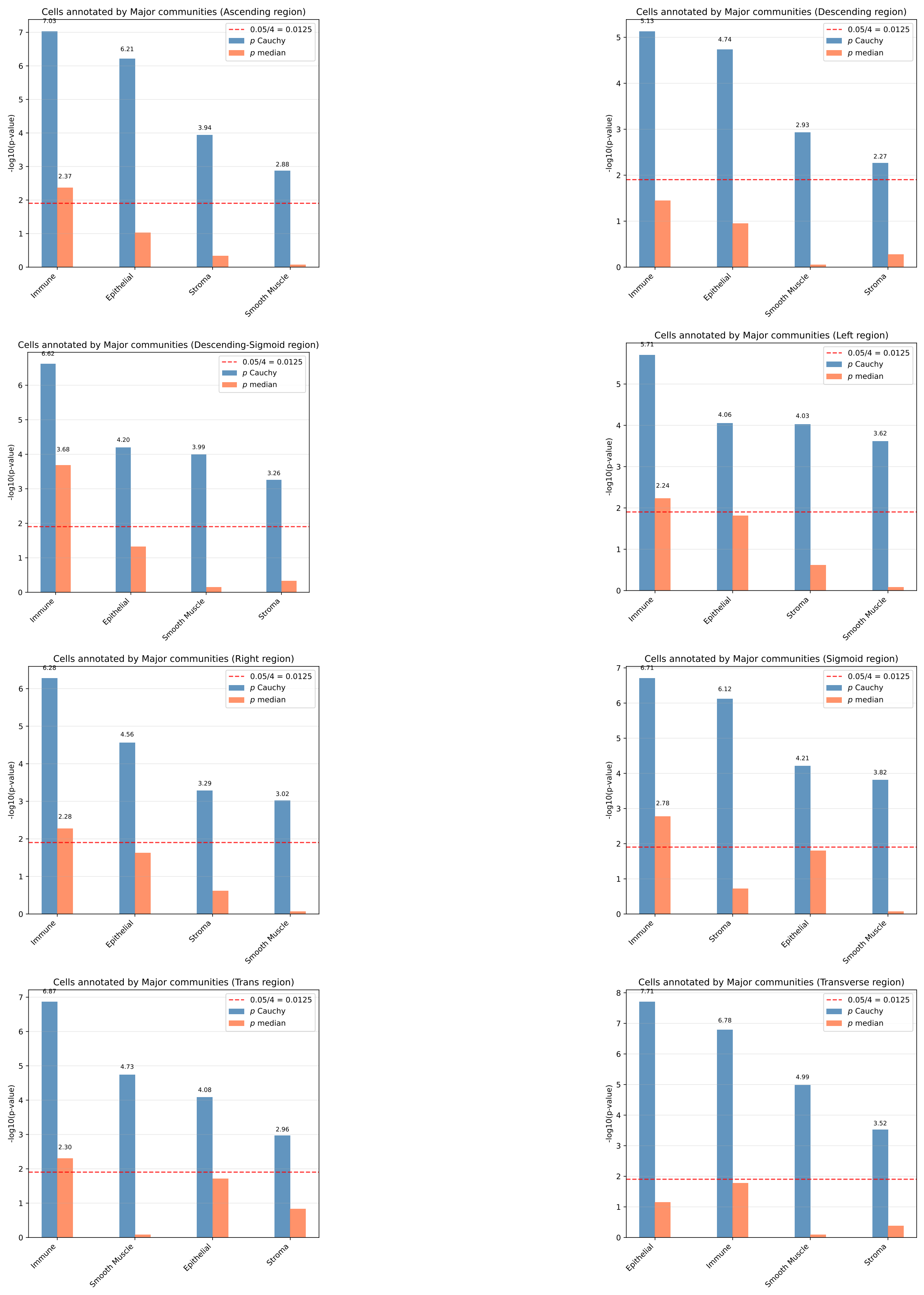

**Figure S30:** We show results of gsMap ((*171*)) for enrichments of selection signals in an intestinal single-cell, spatial transcriptomics dataset ((*173*)) across 8 intestinal regions of the human gut, when annotating cells by major community. See Supplementary Text C for details.

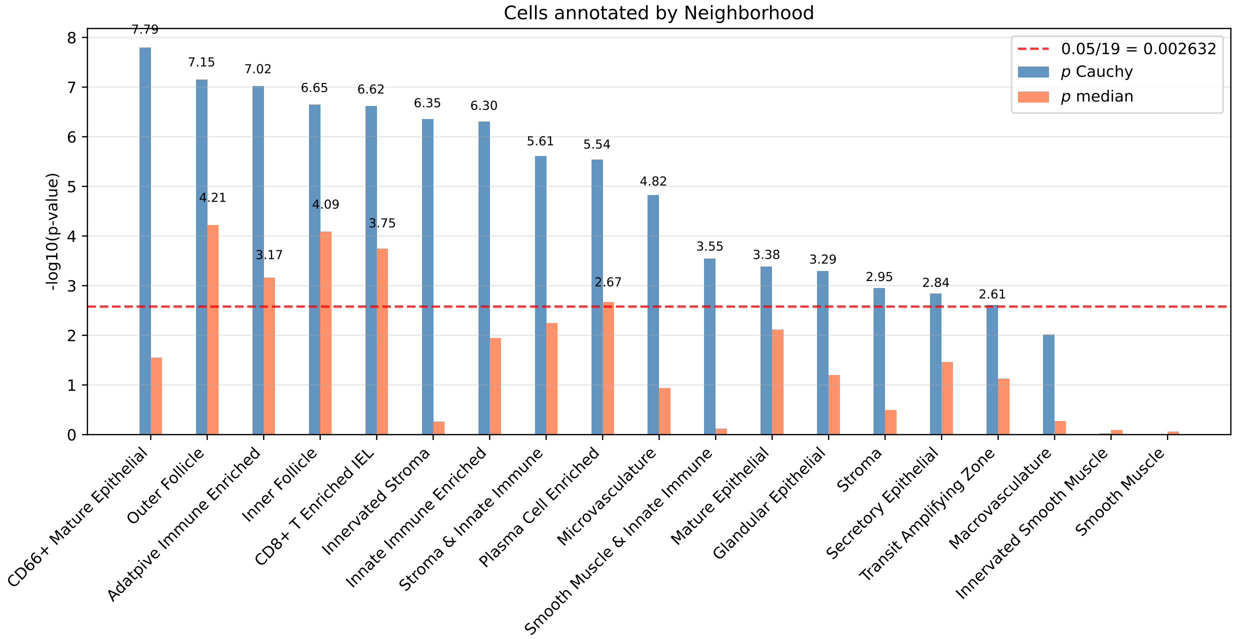

**Figure S31:** We show results of `gsMap` (*171*))for enrichments of selection signals in an intestinal single-cell, spatial transcriptomics dataset (*173*), when pooling data across 8 intestinal regions of the human gut and annotating cells by spatial neighborhood. See Supplementary Text C for details.

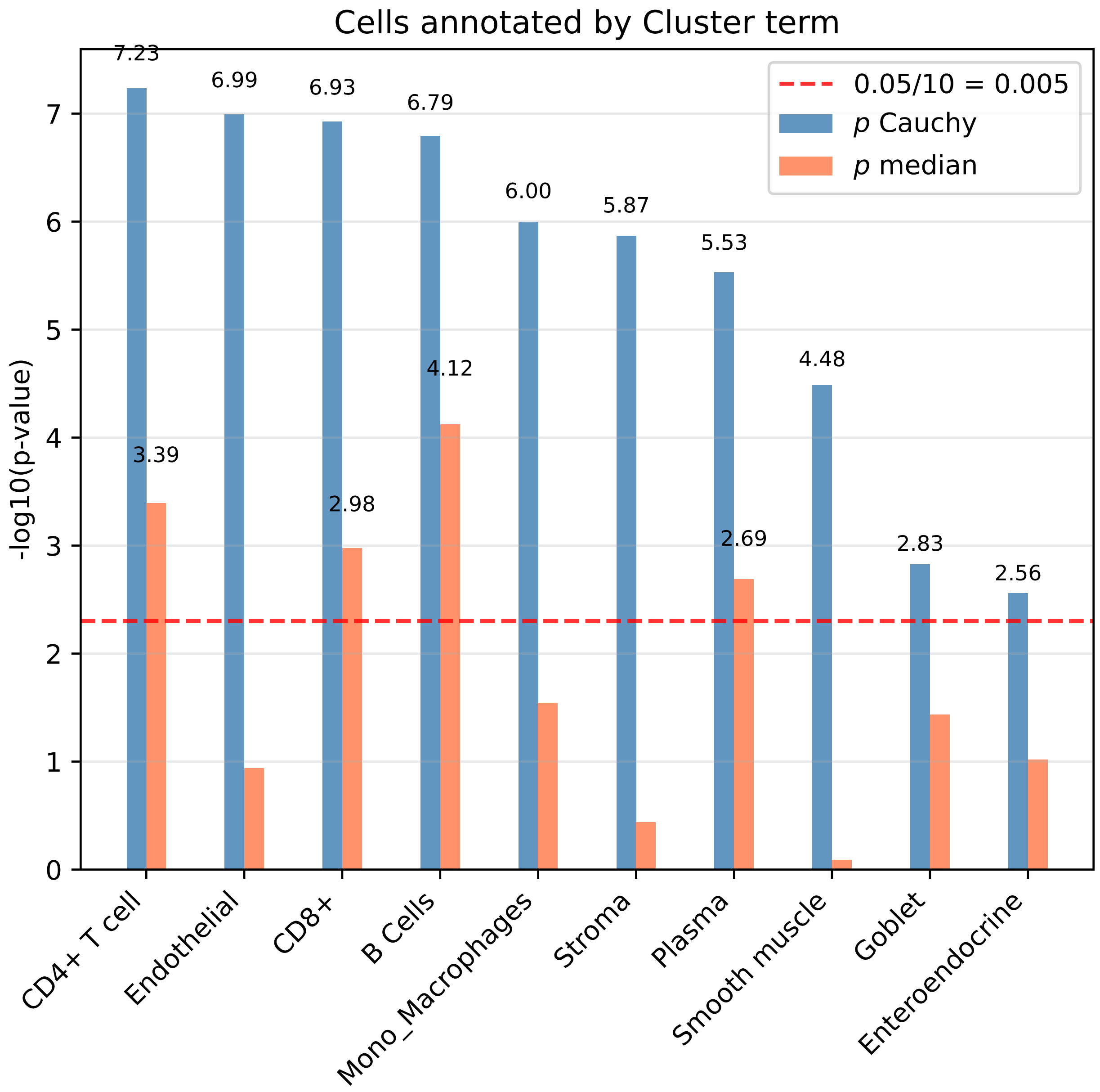

**Figure S32:** We show results of `gsMap` (*171*) for enrichments of selection signals in an intestinal single-cell, spatial transcriptomics dataset (*173*), when pooling data across 8 intestinal regions of the human gut and annotating cells by cell type (cluster term). See Supplementary Text C for details.

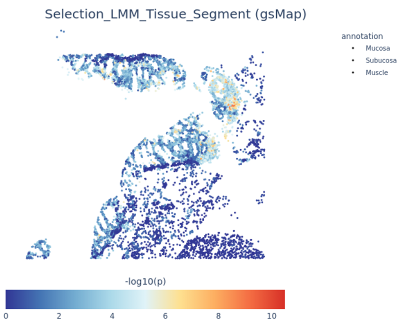

**(a)** B004_Ascending

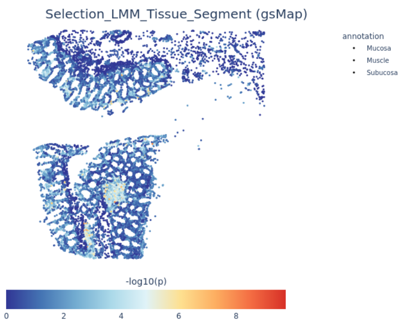

**(b)** B004_Descending

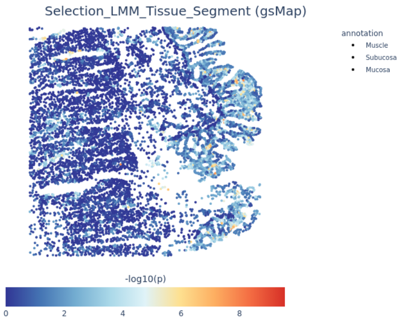

**(c)** B004_Descending-Sigmoid

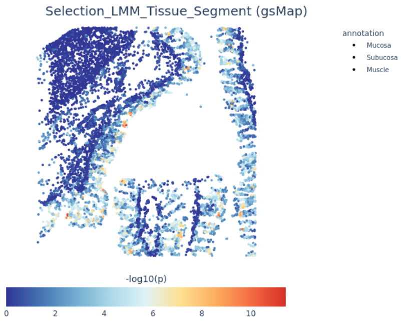

**(d)** B004_Transverse

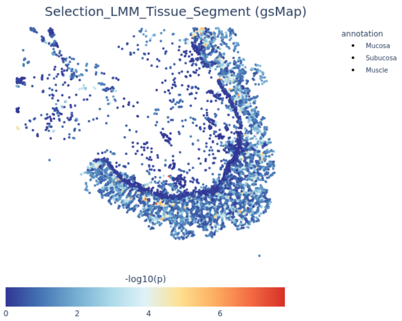

**(e)** B005_Ascending

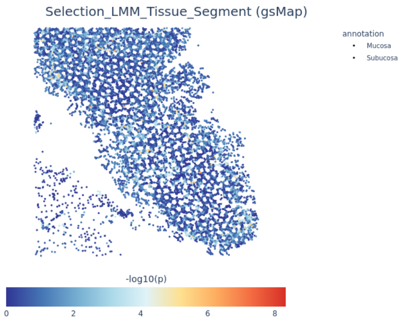

**(f)** B005_Descending

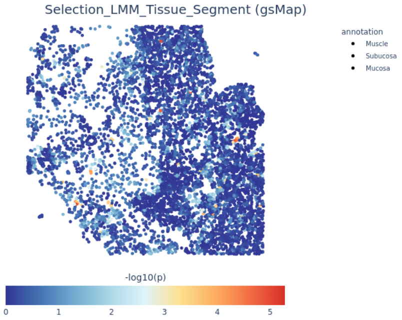

**(g)** B005_Descending-Sigmoid

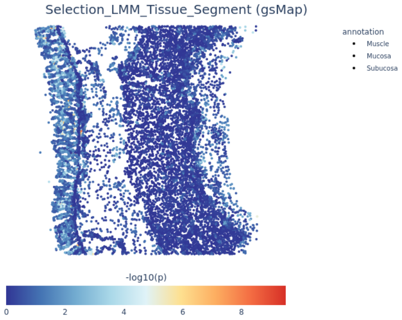

**(h)** B005_Transverse

**Figure S33:** We show results for cell-level enrichment in selection signals inferred using `gsMap`, for additional samples from (*173*) and cells annotated by tissue layer.

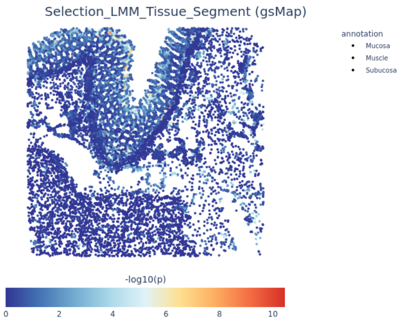

**(a)** B006_Ascending

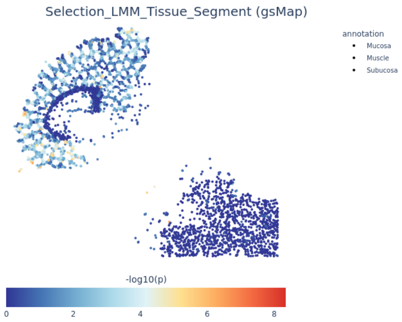

**(b)** B006_Descending

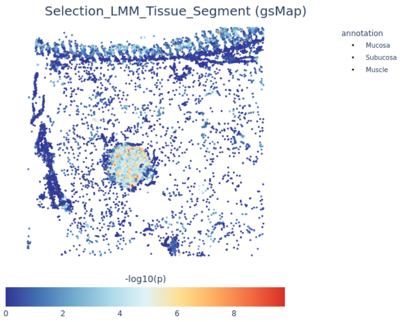

**(c)** B006_Descending-Sigmoid

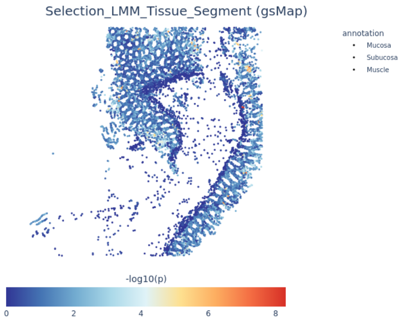

**(d)** B006_Transverse

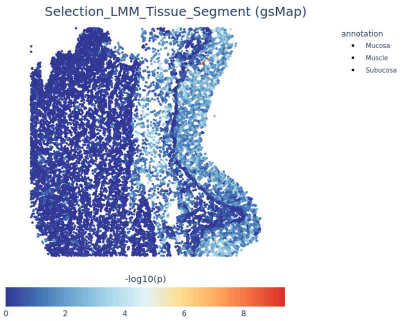

**(e)** B008_Left

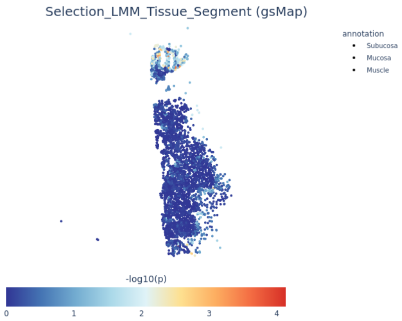

**(f)** B008_Right

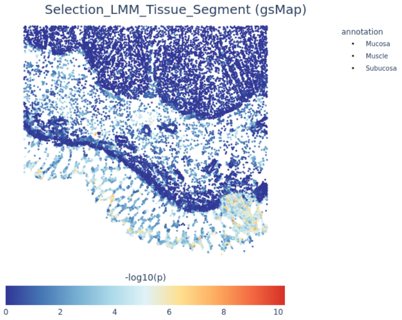

**(g)** B008_Sigmoid

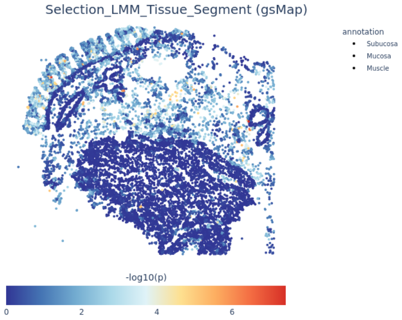

**(h)** B008_Trans

**Figure S34:** We show results for cell-level enrichment in selection signals inferred using `gsMap`, for additional samples from (*173*) and cells annotated by tissue layer.

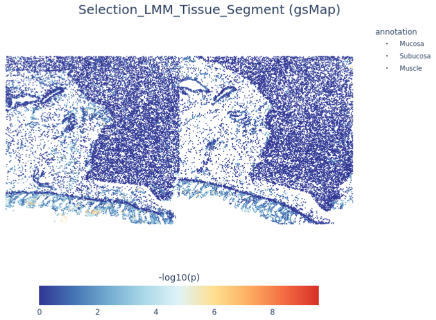

**(a)** B009_Left

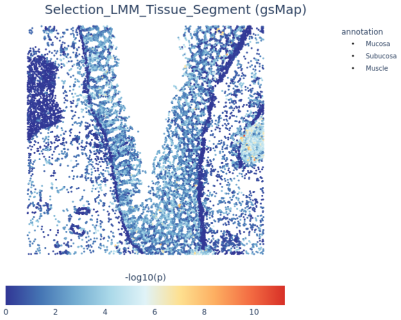

**(b)** B009_Right

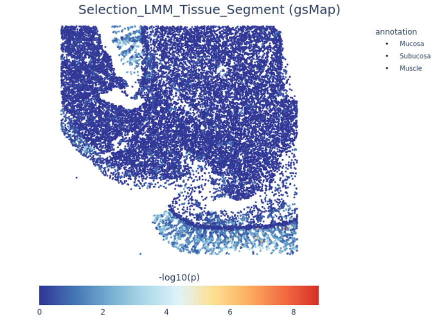

**(c)** B009_Sigmoid

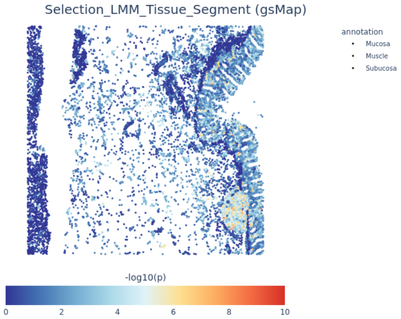

**(d)** B009_Trans

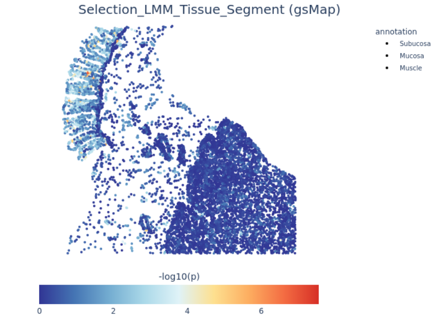

**(e)** B010_Left

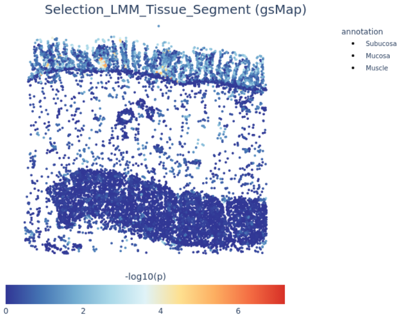

**(f)** B010_Right

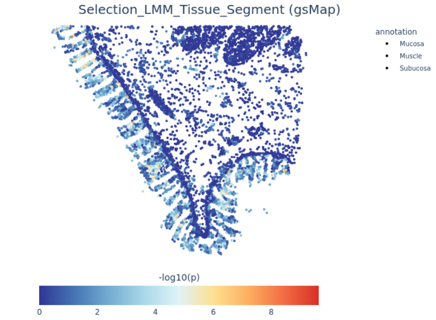

**(g)** B010_Sigmoid

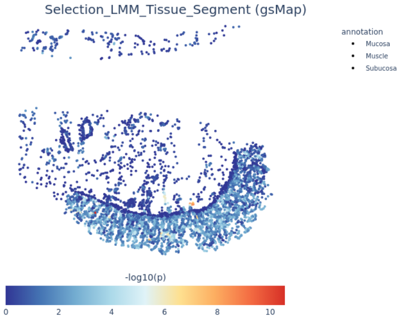

**(h)** B010_Trans

**Figure S35:** We show results for cell-level enrichment in selection signals inferred using `gsMap`, for additional samples from (*173*) and cells annotated by tissue layer.

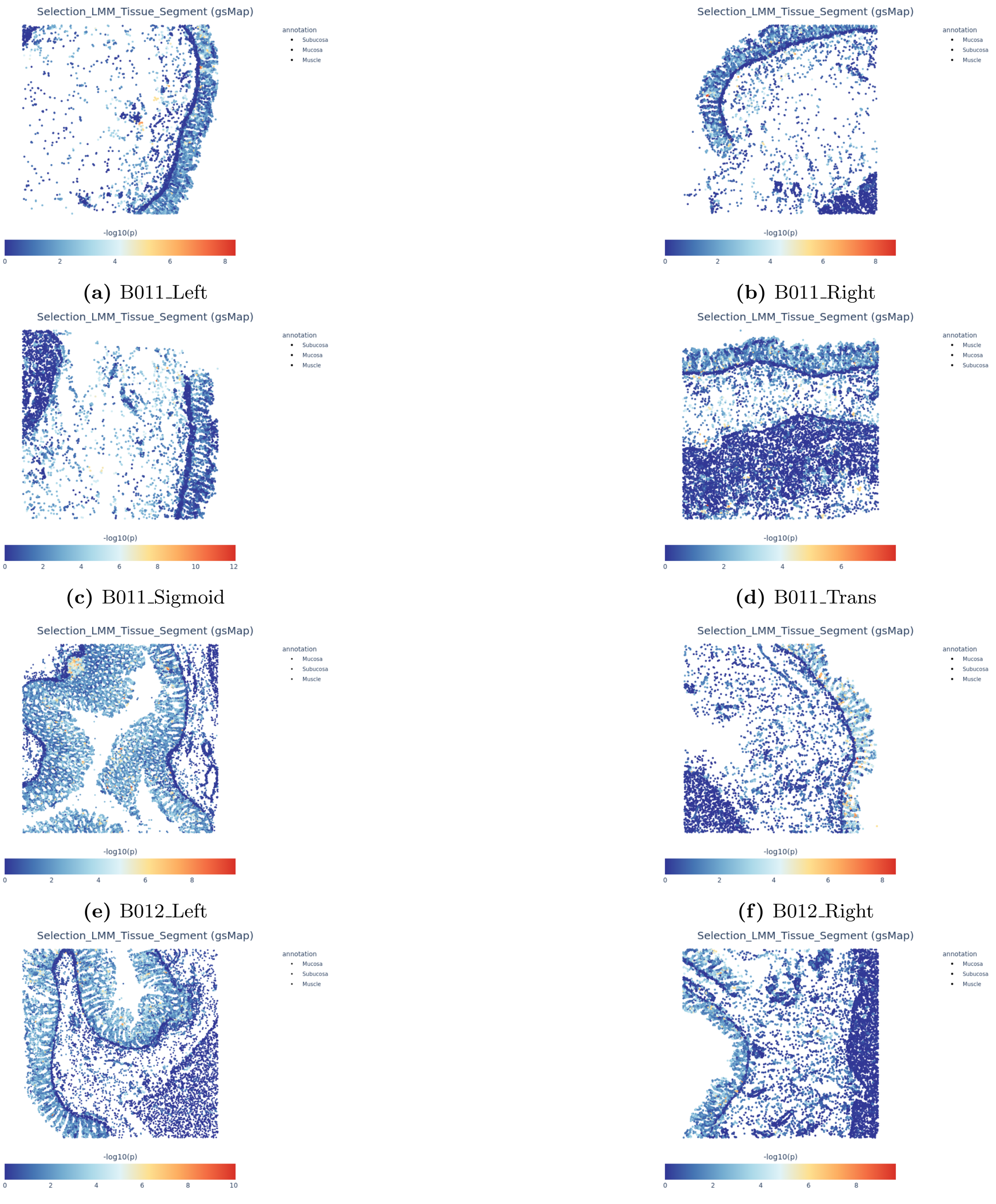

**(a)** B011_Left

**(b)** B011_Right

**(c)** B011_Sigmoid

**(d)** B011_Trans

**(e)** B012_Left

**(f)** B012_Right

**(g)** B012_Sigmoid

**(h)** B012_Trans

**Figure S36:** We show results for cell-level enrichment in selection signals inferred using `gsMap`, for additional samples from (*173*) and cells annotated by tissue layer.

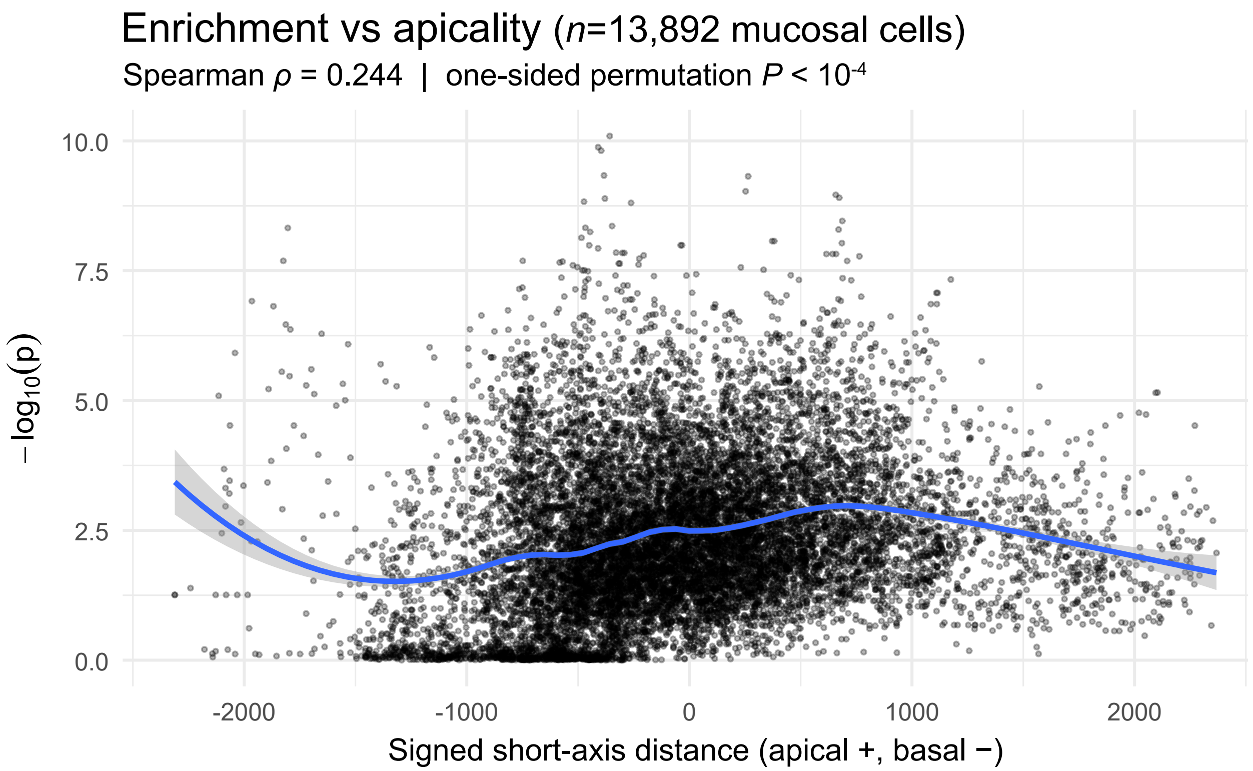

**Figure S37:** We show individual-cell -$\log_{10}(P\text{-value})$ for enrichment in signals of selection (as computed using `gsMap` (*171*) and an intestinal single-cell, spatial transcriptomics dataset (*173*)) against individual-cell position along an apical/basal (closer to/further from the intestinal lumen) axis. See Materials and Methods and Figure S27c,d for details.

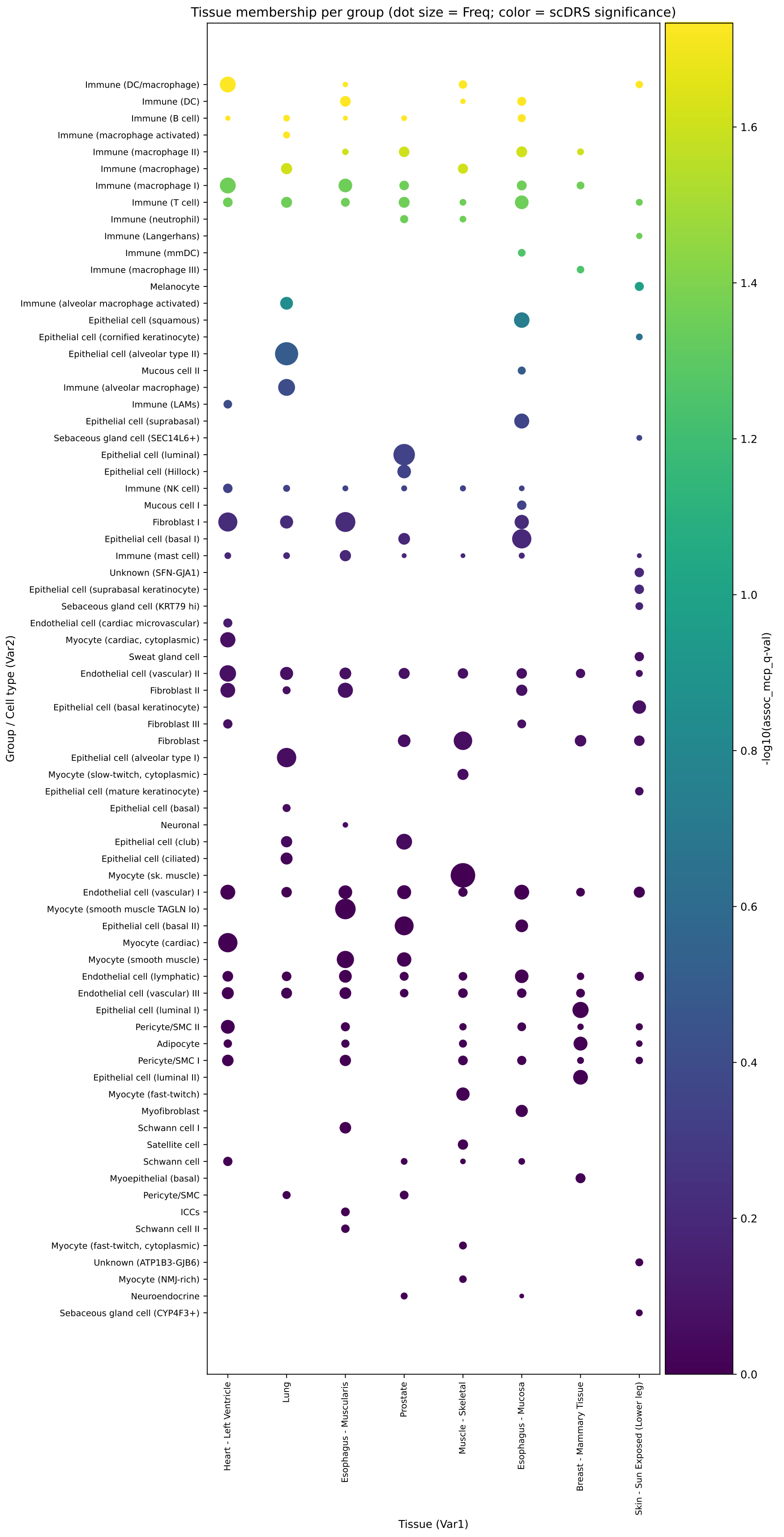

**Figure S38:** Relationship between labeled cell type and tissue of origin in the snGTEx (*176*) dataset. This dataset was analyzed with scDRS (*175*) to look for associations between cells and selection signals (Materials and Methods).

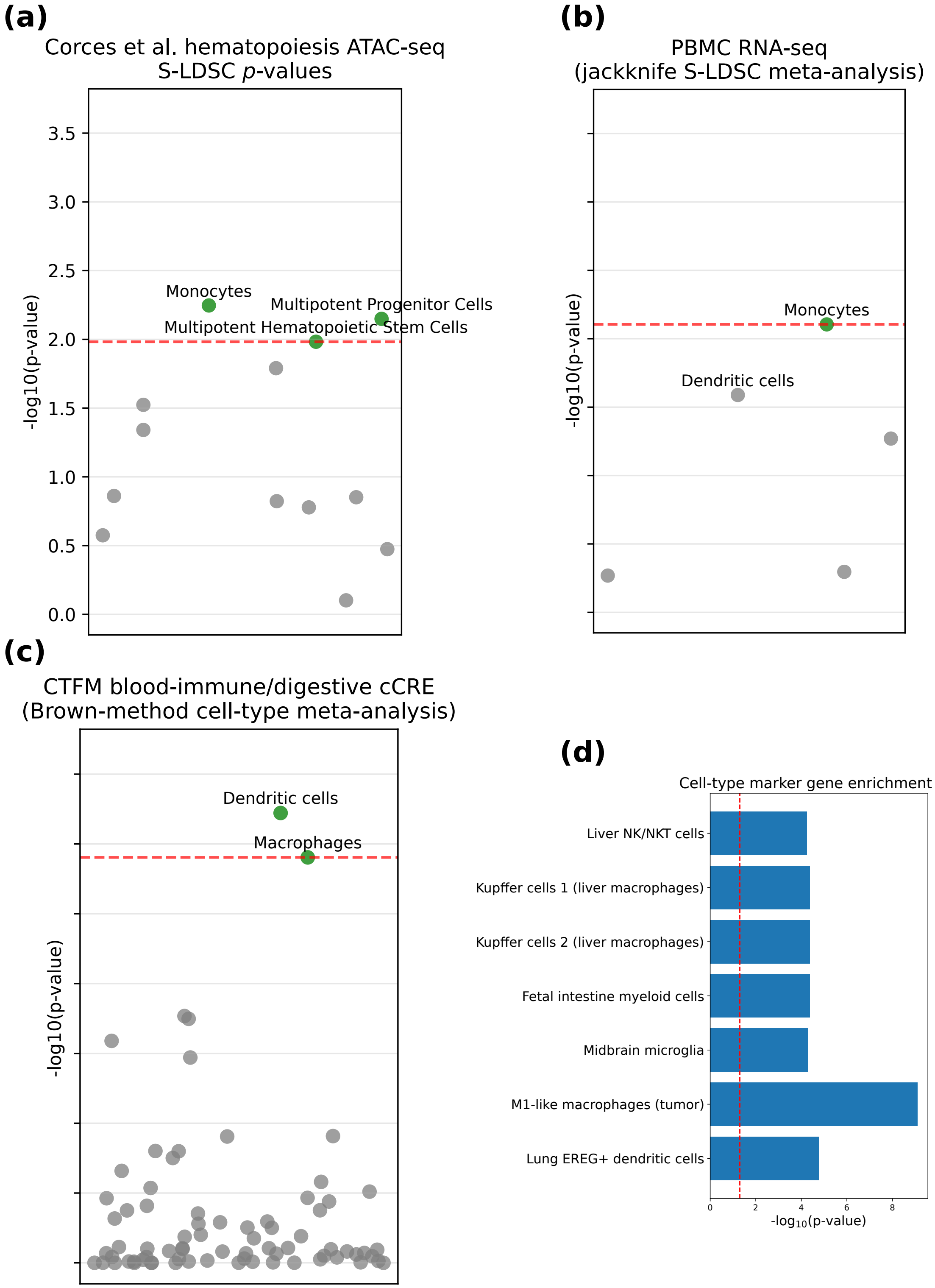

**Figure S39: Secondary cell-type analyses**. We show results of S-LDSC(*102*)-based tests for enrichment of specific cell types in signals of selection, based on **(a)** a chromatin accessibility-based hematopoiesis dataset(*167*), **(b)** single-cell gene expression datasets in blood (we meta-analyzed results for cell types present in all four such datasets analyzed by ref.(*190*) by jackknifing the average S-LDSC regression coefficient across the four), and **(c)** candidate cis-regulatory elements active in specific cell types analyzed by ref.(*192*) (we meta-analyzed results across annotations referring to the same cell type using the empirical Brown method(*193*), using the cross-annotation covariance matrix provided by ref.(*192*)). We also show results for **(d)** a MAGMA gene-set enrichment analysis (analogous to Figure S25c) for sets of genes curated as markers for a particular cell type(*153*). Full numerical results are reported in Data S6.

#### Supplementary Tables

| Regression model | Calibration | Experiment 1 | | Experiment 2.1 | | Experiment 2.2 | | Experiment 3 | |
|---|---|---|---|---|---|---|---|---|---|
| | | TP (%) | FP (%) | TP (%) | FP (%) | TP (%) | FP (%) | TP (%) | FP (%) |
| GLMM | GWAS enrichment | 25.5 | 0 | 59.25 | 0 | 89 | 0 | 22.25 | 0 |
| LMM | GWAS enrichment | 11.25 | 0 | 58.75 | 0 | 80.25 | 0 | 7 | 0 |
| GLMM | Genomic control | 0 | 0 | 28 | 0 | 67.5 | 0 | 0 | 0 |
| LMM | Genomic control | 10.75 | 0 | 58.25 | 0 | 82.25 | 0.25 | 9 | 0 |

**Table S1:** True positives (TP) and false positives (FP) for different regression models using a significance threshold of $|X| > 5.45$, calibrated using two approaches: GWAS enrichment and genomic control. Each experiment includes 800 simulations (400 with directional selection and 400 without). A detailed description of the simulation experiments is provided in Supplementary Information Section 2 of ref.(*86*). See also Supplementary Text A.2

.

| Locus | Putative favored mutation (hg19) | Present in the Akbari et al. scan | Belongs to 95% CS | Within-locus rank |
|---|---|---|---|---|
| *LCT* | 2_136608646_G_A | Yes | Yes | 2 |
| *TLR1* | 4_38798648_C_A | Yes | Yes | 1 |
| *SH2B3* | 12_111884608_T_C | Yes | Yes | 4 |
| *OCA2* | 15_28365618_A_G | Yes | Yes | 10 |
| *SLC45A2* | 5_33951693_C_G | Yes | Yes | 1 |
| *TYR* | 11_88911696_C_A | Yes | Yes | 1 |
| *KITLG* | 12_89328335_T_C | Yes | Yes | – |
| *FUT2* | 19_49206674_G_A | Yes | No | – |
| *CSK* | 15_75077367_C_A | Yes | Yes | 8 |
| *F12* | 5_176841339_T_C | No | – | – |
| *DDB1* | 11_61144652_C_A | No | – | – |
| *TNFSF13B* | 13_108993494_A_G | Yes | No | – |
| *MAPT* | 17_44012257_CG_C | Yes | No | – |

**Table S2:** Agreement between fine-mapping of causal selected variants in real data vs. putative favored mutations for well-characterized selective sweeps. See Supplementary Text A.3.1 for details.

| **Method** | **Level** | **Goal** | **Intuition** | **Reference** |
|---|---|---|---|---|
| cross-trait LDSC | Genome-wide | Estimate correlation between the genetic components of a pair of GWAS traits | Regress product of $Z-$scores on LD scores | Bulik-Sulivan et al. 2015b |
| Fine-mapping | Variant | Estimate probability of a variant having a non-zero effect | Approximate Bayes factor with marginal assoc. | Wakefield 2009 |
| coloc | Locus | Estimate probability of shared causal variants | Renormalize the product of Bayes factors and assess probability mass for each $H_i$ | Giambartolomei et al. 2013 |
| LCV | Genome-wide | Assess a causal relationship between traits | Variants affecting trait A also affecting trait B, but not vice versa (vertical pleiotropy) | O'Connor and Price 2018 |
| PHBC | Genome-wide | Estimate genetic variance of a trait shared with a set of auxiliary traits | cross-trait LDSC pairwise genetic correlations + bias correction | Zhao et al. 2025 |
| MAGMA | Genome-wide | Obtain gene-level association statistics from variant-level association statistics, pathway enrichment | Multiple regression of SNP-level effects on gene membership + covariates | de Leeuw et al. 2015 |
| PoPS | Genome-wide/Locus | Prioritize likely-causal genes | Polygenic enrichment of gene features | Weeks et al. 2023 |
| CALDERA | Locus | Estimate probability of a gene having a non-zero effect | Logistic regression including PoPS scores and physical distance | Schipper et al. 2024 |
| S-LDSC | Genome-wide | Test for signal enrichment in particular variant classes | Excess of heritability explained | Finucane et al. 2015 |
| LDSC-SEG | Genome-wide | Test for signal enrichment in particular tissues and cell-types | S-LDSC with tissue-specific annotations | Finucane et al. 2018 |
| gsMap | Genome-wide | Test for signal enrichment in individual and groups of spatially-arranged cells | S-LDSC with custom annotations | Song et al. 2025 |
| Epigenomic partitioning | Fine-mapped variants | Discover tissue-specific clusters among fine-mapped variants | Variants sharing tissue regulatory activity based on chromatin tracks | Kerner et al. 2026 |
| scDRS | Genome-wide | Test for signal enrichment in individual and groups of cells | Excess expression of GWAS-implicated genes | Zhang et al. 2022 |

**Table S3:** Overview of methods from the GWAS literature applied to analyzing LMM selection statistics in this study.

| Metric | Count |
|---|---|
| Variants with selection PIP $> 0.5$ | 32 |
| Variants with selection and GWAS PIP $> 0.5$ for at least one GWAS trait | 8 |
| Fine-mapped loci colocalizing with at least one GWAS trait | 30 |

**Table S4:** Summary of variants and loci meeting PIP and colocalization criteria. See Data S2 for numerical fine-mapping results and Data S3 for numerical colocalization results.

Legend (colour of Gene cell):
- Infection (antimicrobial)
- Infection (host-pathogen interface)
- Immune response (mucosal barrier)
- Immune response (immune-cell regulation)

| Gene | Notes | Chr (Mbp) |
|---|---|---|
| *BST2* | Interferon-induced tetherin restricting release of enveloped viruses | 19 (18) |
| *LYZ* | Lysozyme that degrades bacterial peptidoglycan (innate effector) | 12 (70) |
| *TMPRSS4* | Epithelial serine protease<br>Receptor for viral pathogens | 11 (118) |
| *FUT6* | Fucosyltransferase impacting surface glycans that influence microbial adhesion and immune recognition | 19 (6) |
| *FUT3* | Fucosyltransferase impacting fucosylated epitopes, altering mucosal glycan landscape and host–microbe interactions | 19 (6) |
| ■*CEACAM5* | Epithelial cell-surface protein at mucosal surfaces<br>Host–microbe interface/mucosal immune interactions | 19 (42) |
| ■*MUC2* | Main secreted intestinal mucin forming the lumen-facing mucus barrier that protects the epithelium from microbes | 11 (1) |
| ■*GP2* | M-cell apical (lumen-facing) glycoprotein binding microbes in the lumen and supporting antigen sampling in gut-associated lymphoid tissue | 16 (20) |
| ■*PIGR* | Transports IgA across mucosal epithelium to generate secretory antibodies that neutralize microbes in the lumen | 1 (207) |
| ■*REG1A* | Secreted protein induced during intestinal inflammation<br>Epithelial repair and mucosal barrier integrity | 2 (79) |
| ■*REG1B* | Secreted protein upregulated with gut epithelial inflammation<br>Mucosal regeneration and barrier protection | 2 (79) |
| *FAM3D* | Gut-secreted homeostatic factor expressed by colon epithelium<br>Colon homeostasis and host defense | 3 (59) |
| *ASAP1* | Cytoskeleton/trafficking regulator<br>Implicated in myeloid-cell motility and TB susceptibility | 8 (131) |
| *LTBR* | Organizes lymphoid tissue and supports inflammatory/antiviral programs | 12 (6) |
| *CXCL10* | Interferon-induced chemokine recruiting T/NK cells to sites of infection for antiviral defense | 4 (77) |
| *LILRA2* | Myeloid receptor modulating innate activation and inflammatory cytokine responses | 19 (55) |
| *CCL21* | Chemokine guiding dendritic cells and T cells<br>Immune-cell trafficking in lymphoid tissues | 9 (35) |
| *TNFRSF9* | T-cell costimulatory receptor<br>T/NK-cell survival and effector function | 1 (8) |

**Table S5: Notes on genes with QTL-colocalizations at loci highlighted in Figure 3a-f.** We provide notes on the function of genes for which the adaptive loci from Figure 3a-f colocalize (including as indicated in the caption of Figure 3), often with one of the candidate causal variants from Figure 3a-f as a fine-mapped QTL. Notes were adapted from protein-function descriptions in ref.(*181*). The ■ symbol indicates genes with a lumen-facing (oriented towards the inside space of bodily cavities) function, in particular in the gut. Full numerical details on colocalizations are given in Data S3.

- Infection (antimicrobial)
- Infection (host-pathogen interface)
- Immune response (mucosal barrier)
- Immune response (immune-cell regulation)

| Gene | Notes | Variant ID (hg19) | Annotation (VEP) | Selection PIP | GWAS/QTL (PIP) | Genomic window (hg19) | Colocalization ($PP_{H4}$) |
|---|---|---|---|---|---|---|---|
| *PLA2G2A* | Secreted phospholipase<br>Antimicrobial activity | 1_20306146_G_C | 5' UTR | 0.19 | *PLA2G2A* pQTL, plasma, UKBB (1)<br>*PLA2G2A* eQTL, prostate, GTEx (0.98)<br>*PLA2G2A* sQTL, prostate, GTEx (0.98) | chr1_18751343_21751343 | *PLA2G2A* pQTL, plasma, UKBB (0.99)<br>*PLA2G2A* eQTL, prostate, GTEx (0.99)<br>*PLA2G2A* sQTL, prostate, GTEx (0.99)<br>Other QTLs |
| *TRIM22* | Antiviral restriction factor<br>Interferon-stimulated | 11_5719228_A_C | intron | 0.07 | *TRIM22* sQTL, monocytes (0.99) | chr11_4177477_7177477 | *TRIM22* sQTL, monocytes (0.99)<br>*TRIM5* sQTL, artery aorta, GTEx (0.99) |
| ■*HSPA6* | Stress-response chaperone | 1_161500130_A_G | downstream | 0.26 | *FCGR3B* sQTL, monocytes (0.89) | chr1_159751343_162751343 | Autoimmune disease (0.99)<br>Immune-cell parameters (0.99)<br>Others |
| *TLR3* | Pattern-recognition receptor | 4_187009014_G_C | 3' UTR | 0.08 | *TLR3* pQTL, plasma, UKBB (0.1)<br>*IFNL1* pQTL, plasma, UKBB (0.98) | chr4_185068786_188068786 | *TLR3* pQTL, plasma, UKBB (0.99)<br>*IFNL1* pQTL, plasma, UKBB (0.99)<br>Others |
| *CLEC12A* | Myeloid inhibitory receptor | 12_10111199_A_G | intron | 0.0012 | *CLEC1B* eQTL, macrophages (0.99)<br>*CLEC12B* eQTL, macrophages (0.99)<br>*CLEC12A* eQTL, macrophages (0.99) | chr12_8170339_11170339 | *CLEC1B* eQTL, macrophages (0.86)<br>*CLEC12B* eQTL, macrophages (0.86)<br>*CLEC12A* eQTL, macrophages (0.86) |
| *TNFSF12* | TNF ligand<br>Inflammatory signaling | 17_7460957_G_T | 3' UTR | 0.73 | *TNFSF12* sQTL, aorta, GTEx (0.99)<br>*TNFSF13* sQTL, aorta, GTEx (0.99)<br>Other tissues | chr17_6000302_9000302 | *TNFSF12* sQTL, aorta, GTEx (0.99)<br>*TNFSF13* sQTL, aorta, GTEx (0.99)<br>Many others |
| *SIGLEC10* | Myeloid/NK activation regulation | 19_51918135_G_T | missense | 0.006 | *SIGLEC10* pQTL, plasma, UKBB (0.78) | chr19_50240554_53240554 | *SIGLEC10* pQTL, plasma, UKBB (0.94)<br>Neutrophil percentage (0.99)<br>Others |
| *ABO* | Glycosyltransferase<br>*ABO* blood group antigens | 9_136128000_G_C | 3' UTR | 0.03 | *ABO* eQTL, skin sun exposed, GTEx (0.99) | ■chr9_136125097_137125097 | *ABO* sQTL, thyroid, GTEx (0.93)<br>*ABO* eQTL, regulatory T cells (0.95)<br>*ABO* eQTL, skin sun exposed, GTEx (0.99)<br>Monocyte percentage (0.99) |
| *ABO* | Glycosyltransferase<br>*ABO* blood group antigens | 9_136132954_T_C | 3' UTR | 0.01 | *ABO* sQTL, thyroid, GTEx (0.93)<br>*ABO* eQTL, regulatory T cells (0.95) | ■chr9_136125097_137125097 | *ABO* sQTL, thyroid, GTEx (0.93)<br>*ABO* eQTL, regulatory T cells (0.95)<br>*ABO* eQTL, skin sun exposed, GTEx (0.99)<br>Monocyte percentage (0.99) |
| *FUT2* | Secretor fucosyltransferase<br>Mucosal glycan synthesis | 19_49206674_G_A | synonymous | 0.34 | *FUT2* sQTL, minor salivary gland, GTEx (0.99)<br>*TNFRSF12A* pQTL, plasma, UKBB (0.60) | ■chr19_47240554_50240554 | *TNFRSF12A* pQTL, plasma, UKBB (0.99)<br>Type 1 diabetes (0.99)<br>Many others |
| *FAM26F* | Interferon-induced<br>T-cell activation marker | 6_116783330_G_A | upstream | 0.03 | *FAM26F* eQTL, stimulated T cells (0.99) | chr6_115192106_118192106 | *FAM26F* eQTL, stimulated T cells (0.99)<br>Ulcerative colitis (0.97)<br>Others |
| *KLRC2* | NK receptor<br>Antiviral/activation signaling | 12_10581650_G_C | downstream | 0.009 | *KLRC4-KLRK1* eQTL, T cells (0.60) | chr12_9170339_12170339 | *KLRC4-KLRK1* eQTL, T cells (0.97) |
| *SH2B3* | Immune signaling adaptor<br>Lymphocyte-pathway regulation | 12_111884608_T_C | missense | 0.08 | Lymphocyte count (0.99)<br>Autoimmune disease (0.99)<br>Others | chr12_110170339_113170339 | Lymphocyte count (0.99)<br>Autoimmune disease (0.99)<br>Others |
| | | 13_108993494_A_G | intergenic | 0.002 | Leucocyte count (0.85)<br>*CD27* pQTL, plasma, UKBB (0.90)<br>*CR2* pQTL, plasma, UKBB (0.86) | chr13_107046435_110046435 | Leucocyte count (0.97)<br>*CD27* pQTL, plasma, UKBB (0.94)<br>*CR2* pQTL, plasma, UKBB (0.94) |
| *PRKCH* | Immune-cell signaling | 14_61810980_A_G | intron | 0.003 | *PRKCH* eQTL, stimulated macrophages (1)<br>*PRKCH* eQTL, stimulated monocytes (0.99) | chr14_60398369_63398369 | *PRKCH* eQTL, stimulated macrophages (0.94)<br>*PRKCH* eQTL, stimulated monocytes (0.94)<br>Lymphocyte percentage (0.98)<br>Others |
| *KSR1* | Signaling in inflammation | 17_25869033_A_C | intron | 0.1 | *KSR1* eQTL, stimulated monocytes (0.98) | chr17_24000302_27000302 | *KSR1*, stimulated monocytes (0.99)<br>Psoriasis (0.97) |
| *TNFRSF13C* | BAFF receptor<br>B-cell survival and maturation | 22_42322716_G_C | missense | 0.1 | *TNFRSF13C* pQTL, plasma, UKBB (1) | chr22_41070120_44070120 | *TNFRSF13C* pQTL, plasma, UKBB (0.99)<br>Chronic tonsil/adenoid disease (0.99)<br>Many others |

**Table S6: Additional fine-mapped candidate causal selected variants**. We report additional fine-mapped candidate causal selected variants inferred from selection–immune gene QTL colocalization (often with GWAS colocalizations for immune-related traits that are not infectious diseases) together with QTL and/or immune-trait GWAS fine-mapping. Notes were adapted from protein-function descriptions in ref.(*181*). Variants were annotated using the Ensembl variant effect predictor(*194*). The ■ symbol indicates ambiguity of some sort: 1_161500130_A_G is annotated as downstream of *HSPA6*, but it more plausibly acts on immunity through *FCGR3B*, a neutrophil antibody receptor for which it is a fine-mapped QTL; chr9_136125097_137125097, containing the *ABO* genes, implicated two different candidate causal variants, with conflicting QTL fine-mapping results in different tissues; and chr19_47240554_50240554, containing the *FUT2* gene, implicated a synonymous SNP distinct from the well-known stop-gain SNP for this gene, previously linked to genetic adaptation to infections(*195*). Full numerical details are given in Data S3.

- Pigmentation
- Metabolism/Transport
- Cardiovascular/Blood

| Gene | Notes | Variant ID (hg19) | Annotation (VEP) | Selection PIP | GWAS/QTL (PIP) | Genomic window (hg19) | Colocalization ($PP_{H4}$) |
|---|---|---|---|---|---|---|---|
| *RMDN2* | Microtubule dynamics | 2_38298139_T_C | downstream | 0.81 | Skin color (0.99) | chr2_37011486_40011486 | Skin color (0.99) |
| *SLC45A2* | Melanosomal transporter | 5_33951693_C_G | missense | 1 | Skin color (1)<br>Others | chr5_32021182_35021182 | Skin color (1)<br>Others |
| *TYRP1* | Melanin pathway protein | 9_12712157_G_A | downstream | 0.002 | Skin color (0.96) | chr9_11125097_14125097 | Skin color (0.99) |
| *TYR* | Tyrosinase<br>Melanin biosynthesis | 11_88911696_C_A | missense | 0.61 | Black hair color (0.99) | chr11_87177477_90177477 | Black hair color (0.99) |
| | | 12_89328335_T_C | regulatory | 0.001 | Skin color (1) | chr12_88170339_91170339 | Skin color (0.94) |
| *EGLN3* | Hypoxia response pathway | 14_34672770_T_C | intron | 0.2 | *EGLN3* eQTL, skin, GTEx (0.99) | chr14_34398369_35398369 | *EGLN3* eQTL, skin, GTEx (0.99)<br>Skin color (0.99) |
| *SLC24A5* | Cation exchanger | 15_48426484_A_G | missense | 0.32 | Skin color (0.93) | chr15_47150423_50150423 | Skin color (0.99) |
| *MARC1* | Mitochondrial enzyme | 1_220970028_A_G | missense | 0.34 | Cholesterol (0.99)<br>Liver fat (0.80) | chr1_219751343_222751343 | Cholesterol (0.99)<br>Liver fat (0.99) |
| *SLC39A8* | Zinc transporter | 4_103188709_C_T | missense | 0.16 | Waist-to-hip ratio (0.99)<br>Many others | chr4_102068786_105068786 | Waist-to-hip ratio (0.99)<br>Many others |
| *SLC22A4* | Organic cation transporter | 5_131640536_A_G | intron | 0.98 | *SLC22A4* eQTL, fibroblasts, GTEx (0.99) | chr5_130021182_133021182 | *SLC22A4* eQTL, fibroblasts, GTEx (0.99)<br>Others |
| *SERPINA1* | Alpha-1 antitrypsin<br>Liver/lipid traits | 14_94847262_T_A | missense | 0.05 | Lipid metabolism disorders (0.84)<br>*SERPINA1* pQTL, plasma, UKBB (1) | chr14_93398369_96398369 | Lipid metabolism disorders (0.99)<br>*SERPINA1* pQTL, plasma, UKBB (0.99)<br>Many others |
| *F12* | Coagulation factor XII | 5_176839890_T_G | upstream | 0.55 | *F12* pQTL, plasma, UKBB (0.98) | chr5_175021182_178021182 | *F12* pQTL, plasma, UKBB (0.99)<br>Many others |
| *PKD2L1* | Cation channel | 10_102075479_G_A | intron | 0.3 | Red Cell Distribution (0.99)<br>Many others | chr10_100083359_103083359 | Red Cell Distribution (0.99)<br>Many others |
| *RIN3* | Endocytosis regulator<br>Platelet traits | 14_93118229_C_T | missense | 0.005 | Platelet crit (0.96)<br>*RIN3* apaQTL, whole blood, GTEx (0.67) | chr14_91398369_94398369 | Platelet crit (0.94)<br>*RIN3* apaQTL, whole blood, GTEx (0.90)<br>Many others |
| *ARNT2-DT* | lncRNA near ARNT2 | 15_80676925_A_G | intron | 0.003 | Atrial fibrillation (1) | chr15_78150423_81150423 | Atrial fibrillation (0.94)<br>Others |
| *EPS15L1* | Endocytosis adaptor | 19_16499569_C_T | intron | 0.005 | Lymphocyte count (1) | chr19_15240554_18240554 | Lymphocyte count (0.99)<br>Ischemic heart disease (0.99)<br>Blood pressure (0.99)<br>Others |
| *MYO9B* | Motor protein | 19_17219105_G_A | intron | 0.06 | Blood pressure (0.98)<br>*MYO9B* eQTL, fibroblasts, GTEx (0.98) | chr19_15240554_18240554 | Blood pressure (0.99)<br>*MYO9B* eQTL, fibroblasts, GTEx (0.99)<br>Others |

**Table S7: Additional candidate causal selected variants not primarily related to immunity, inferred from selection–GWAS and selection–QTL colocalizations**. We report additional fine-mapped candidate causal selected variants inferred from integration of selection, QTL, and GWAS colocalization and/or fine-mapping, at loci not primarily related to immunity. Notes were adapted from protein-function descriptions in ref.(*181*). Variants were annotated using the Ensembl variant effect predictor(*194*). Full numerical details are given in Data S3.

| Parameter | Definition | Estimate | SE | $z$ | $p$ |
|---|---|---|---|---|---|
| $\mathrm{cor}(S, U)$ | selection–UC correlation | 0.00847 | 0.00407 | 2.08 | 0.040 |
| $\mathrm{cor}(S, Z_{\mathrm{inf}})$ | selection–infection $Z$ correlation | −0.00919 | 0.00417 | −2.20 | 0.030 |
| $b$ | slope in $U = \alpha + bT$ ($T = -Z_{\mathrm{inf}}$) | −0.02669 | 0.00346 | −7.71 | $1.0 \times 10^{-11}$ |
| $\mathrm{Cov}(S, U_T)$ | infection-linked component ($= b\,\mathrm{Cov}(S, T)$) | $-3.03 \times 10^{-4}$ | $1.43 \times 10^{-4}$ | −2.12 | 0.036 |
| $\mathrm{Cov}(S, U_{\mathrm{res}})$ | residual component | 0.01123 | 0.00524 | 2.14 | 0.035 |
| $f_T$ | fraction $\mathrm{Cov}(S, U_T)/\mathrm{Cov}(S, U)$ | −0.0277 | 0.0211 | −1.32 | 0.191 |

**Table S8:** Selection–ulcerative colitis (UC) decomposition with respect to protection against intestinal infections. $S$ is the selection statistic ($Z_s$; A1 allele favored if $S > 0$), $U$ is the UC GWAS $Z$-score, and $Z_{\mathrm{inf}}$ is the intestinal infections GWAS $Z$-score; $T \equiv -Z_{\mathrm{inf}}$ just recodes the $Z$-score in the protective direction. The negative $\mathrm{Cov}(S, U_T)$ indicates that the selection component aligned with infection protection is UC-protective, while the residual component is UC-risk increasing. See Supplementary Text D for full details.

### Supplementary Data

Provided as Excel tables.

I. Data S1: GWAS analyses.

**Sheet 1.** Source of GWAS data analyzed.

**Sheet 2.** Genetic correlations between LMM selection statistics and GWAS summary statistics, estimated by cross-trait LDSC.

**Sheet 3.** Genetic correlations among immune-related GWAS traits having a non-zero genetic correlation with LMM selection statistics (FDR < 20 %).

**Sheet 4.** Results of a test for significant increases or decreases over time of genetically-predicted GWAS trait values ($\gamma_{\text{sign}}$).

**Sheet 5.** LCV results between LMM selection statistics and GWAS summary statistics.

**Sheet 6.** PHBC results between LMM selection statistics and sets of GWAS auxiliary traits.

**Sheet 7.** Functional enrichment of fine-mapped selected variants.

**Sheet 8.** Results of applying S-LDSC with the baselineLD model to LMM selection statistics.

II. Data S2: Fine-mapped selected variants and gene-level analyses:

**Sheet 1.** Complete selection fine-mapping results with credible sets.

**Sheet 2.** MAGMA results for selection.

**Sheet 3.** PoPS results for selection.

**Sheet 4.** CALDERA results for selection.

**Sheet 5.** S-LDSC results including immune-genes and all-genes annotation with the baselineLD v2.2 model.

III. Data S3: Summary of colocalizations and fine-mapping convergence:

**Sheet 1.** Variant-level summary for variants with convergent evidence from selection and GWAS/QTL fine-mapping.

**Sheet 2.** Variant annotation by Ensembl-VEP.

**Sheet 3.** Locus-level results of colocalization tests for loci with convergent evidence for selection and GWAS/QTL association.

**Sheet 4.** Immune-cell QTL studies retrieved from the eQTL catalog.

IV. Data S4: Pathway/gene-set analyses:

**Sheet 1.** Overlap between CALDERA fine-mapped genes for selection and selection-QTL colocalized genes.

**Sheet 2.** Formal assessment of immune-related pathways being more likely to be enriched for selection genes after adjusting for pathway size.

**Sheet 3.** Results of pathway-enrichment analyses for CALDERA>0.1 or QTL-colocalized genes for selection.

**Sheet 4.** Results of tests for a directional effect of positively-selected alleles on pathway activity.

**Sheet 5.** Overlap between pairs of positive/negative regulators of the same biological process.

**Sheet 6.** S-LDSC results for blood cellular processes from (Jagadeesh, Key et al. 2022, *Nat Gen*).

**Sheet 7.** MAGMA gene-set enrichment results for selection.

**Sheet 8.** Results for MAGMA gene-set enrichment for *Inborn Errors of Immunity* gene sets and tuberculosis GWAS.

#### V. Data S5: Tissue analyses.

**Sheet 1.** Orthogonal clusters of gene-level features selected by `PoPS` for selection.

**Sheet 2.** Contribution of each cluster to `PoPS` scores for selection.

**Sheet 3.** Manually-chosen labels for `PoPS` clusters for selection.

**Sheet 4.** Ranked gene-set enrichment results for human-airway gene features used to annotate ambiguous `PoPS` clusters.

**Sheet 5.** LDSC-SEG results for the `Multi_tissue_gene_expr` dataset.

**Sheet 6.** LDSC-SEG results for the `Multi_tissue_chromatin` dataset.

**Sheet 7.** Epigenomic partitioning results for selection (matrix $H$, cluster-to-tissue).

**Sheet 8.** Epigenomic partitioning results for selection (matrix $W$, variant-to-cluster).

**Sheet 9.** `gsMap` results for LMM selection statistics with the human-gut single-cell, spatial transcriptomics dataset by (Hickey et al. 2023, *Nature*).

**Sheet 10.** Per-cell apicality for mucosal cells from sample B012 from (Hickey et al. 2023, *Nature*).

#### VI. Data S6: Cell-type analyses.

**Sheet 1.** `scDRS` results for LMM selection statistics for both group-level association (`assoc_mcp`) and heterogeneity (`hetero_mcp`) in snGTEx.

**Sheet 2.** `scDRS` results for LMM selection statistics for individual cells in snGTEx.

**Sheet 3.** LDSC-SEG results for LMM selection statistics and the Corces et al. ATAC-seq dataset.

**Sheet 4.** S-LDSC results for LMM selection statistics and meta-analyzed cell-types across the 4 blood-related datasets from (Jagadeesh, Key et al. 2022, *Nat Gen*).

**Sheet 5.** S-LDSC results for LMM selection statistics and blood and digestive CTFM annotations.

**Sheet 6.** Empirical-Brown-method meta-analysis results across cell types from blood and digestive CTFM annotations.

**Sheet 7.** MAGMA gene-set enrichment results for sets of cell-type marker genes.
